# Integrated miRNA-protein profiling of small extracellular vesicle cargo reveals dynamic cargo remodelling during tissue repair in UV-C-damaged *C. elegans*

**DOI:** 10.64898/2026.09.23.753761

**Authors:** Mukesh Behera, Srianshu Kumar Panda, Jaspreet Kaur, Gaurav Sharma, Manshi Sharma, Abhay Kumar Sharma, Parmeshwar Sahu, Sundeep Singh Saluja, Jaswinder Singh Maras, Narender K Dhania

## Abstract

Ultraviolet-C (UV-C) radiation is a potent environmental stressor that induces DNA damage, accumulation of reactive oxygen species (ROS), and cellular dysfunction. This also initiates coordinated molecular responses to regulate cellular adaptation and recovery. Small extracellular vesicles (sEVs) have emerged as key mediators of intercellular communication through the selective incorporation of microRNAs (miRNAs), proteins, and lipids. However, the temporal changes of sEV-associated cargo during the progression from UV-C-induced damage to recovery remain uncharacterized. We hypothesised that UV-C exposure induces stage-specific changes in sEV cargo (miRNA and proteins) that promote the transition of cells from damage to the response and recovery phases. To examine this, a time-resolved UV-C exposure model was established using *Caenorhabditis elegans* (*C. elegans*). In this paper, we studied 2.5 hours of UV-C exposure, including cellular damage, intracellular ROS levels, and DNA integrity to define the damage and repair state. Based on these temporal variations, samples were collected: i) immediately after exposure, ii) 6 hours post-exposure, and iii) 16 hours post-exposure were designated as damage, response, and recovery conditions, respectively. sEVs isolated from whole-worm preparations were characterised using biophysical and molecular approaches, followed by miRNA sequencing and proteomic analyses. Integrative analysis further identified miRNA–protein associations and associated them with pathways involved in stress adaptation, metabolism, signal transduction, and cellular homeostasis. Collectively, these findings provide a temporal molecular framework of sEV cargo associated with UV-C-induced damage, response, and recovery in *C. elegans*. Overall, this study identified candidate molecular signatures and miRNA– protein associations that underlie the temporally selective packaging of sEV-associated cargo following UV-C-induced damage, providing a basis for future tissue-specific and mechanistic investigations.

## Introduction

UV radiation is an inevitable part of the environment and is associated with DNA damage, ROS production, immunosuppression, and cell death. Skin, being the outermost organ, is continuously exposed to UV radiation, leading to various uninvited consequences. UV radiation is divided into UV-A (320-400nm), UV-B (280-320nm), and UV-C (100-280nm). UV-A and UV-B can enter the Earth’s surface, whereas UV-C cannot penetrate it[1]. UV-C exposure of Earth’s surface occurs via artificial sources used for disinfection, laboratory research, and industrial welding. Around 45% of the global population (∼3.6 billion people) receives high-to extreme-level UV (UV Index above 8)[2, 3]. Metalworkers are more prone to UV-C irradiation and its associated damage, which ultimately leads to lasting chronic issues. UV damage causes oxidative stress via ROS production; it also initiates DNA damage by generating photodimers at di-pyrimidine sites, thereby stressing cells and organisms by obstructing DNA replication and transcription. Cells also undergo cell death via cell cycle arrest and apoptosis to cease proliferation of the damaged cells[4]. UV-induced skin inflammation stimulates expansion of immunosuppressive regulatory T cells. WHO has listed UV radiation as a distinguished carcinogen because it is both a mutagen and a nonspecific damaging agent with qualities of a tumour initiator and promoter[3]. UV radiation also promotes pterygium and cataract in the eyes[3, 5], follicular damage of hair, altered skin flora, vasodilation and leukaemia[6]. Additional problems associated with skin include erythema, tanning, photoaging, and wrinkling [7, 8]. Moreover, understanding how organisms respond and recover, and how cells communicate during UV exposure, remains a biological question.

While previous research has elucidated the mechanism of UV-induced damage in organisms, a research gap persists regarding the sequential approach to the repair mechanism following UV-induced damage. Elucidating these sequences is important to overcome the challenges associated with cellular repair. Over the years, various regulatory molecules (cytokines, chemokines, growth factors, phytochemicals, sEVs) have been studied for their role in damage repair[9–11]. Although sEVs are known mediators of intercellular communication, their communicative role has attracted significant attention in recent years. The efficient role of small sEVs in UV-damaged tissue repair remains largely unexplored. Also, variations in sEVs cargo (miRNA, proteins) during UV-C-induced tissue damage and the subsequent repair phases remain unclear. sEVs are lipid bilayered, nanosized vesicles that play a vital role in cell signalling, acting as carriers of bioactive molecules such as proteins, lipids, and nucleic acids^11^. sEVs, which were initially considered as cellular “trash bins”, are now widely known for their role in cellular communication. For example, (i) in response to tissue damage, Mesenchymal Stem Cells (MSCs) derived EVs of human umbilical cord attenuate H_2_O_2_-induced epithelial cell death in mice[12], (ii) Programmed death-ligand 1 (PD-L1) is secreted from human lung fibroblast EVs in response to TGF-β, which helps in tissue remodeling[13], (iii) muscle-derived EVs are important for modulating cell metabolism, differentiation, regeneration and repair[14, 15]. Previous reports also suggest that EV-induced damage can be mitigated by the intervention of sEVs through reducing γH2AX foci, reducing matrix metalloproteinase 1 (MMP-1), increasing Glutathione Peroxidase-1 (GPX-1) activity and increasing collagen deposition. In the last decade, there has been rapid progress in the therapeutic/diagnostic implementation of sEVs. However, the association of sEVs and UV-C-damaged tissue repair is yet to be established. In particular, the dynamic changes in sEV-associated miRNAs and proteins during damage and subsequent processes have not been studied together. In this paper, we hypothesised that sEVs’ molecular signatures (miRNA and protein composition) undergo temporal shifts in UV-C-induced damage and subsequent repair processes. We profiled whole-worm population-derived sEVs, miRNAs, and proteins during damage, response, and recovery stages post-UV-C exposure. sEVs were isolated by size exclusion chromatography combined with an ultrafiltration method (SEC-UF), followed by characterisation. Further, miRNA and proteins were isolated and analysed. We reported miRNAs and proteins at different time points of tissue repair. Our results identified differentially expressed miRNAs and proteins in sEVs that were dynamically adaptive across different phases of the recovery process.

## Results

### Effects of UV-C exposure on *C. elegans*

To assess biological impact of UV-C exposure using the touch response assay, *C. elegans* were continuously exposed to UV-C (0.728 mW) for 4 hours. After 1 hour of UV-C exposure, about 10-15% of worms were dead (Figure 1a). Dead count increased to ∼20% after 1.5 hours of exposure (Figure 1a). Mortality increased significantly after 2.5 hours, reaching ∼50%. Prolonged exposure for more than 3.5 hours resulted in severe damage, with ∼90% mortality. (Figure 1a). Further, cuticle exposed to UV-C for 2.5 hours showed no evident alterations in surface morphology compared with a non-exposed control worm (Figure 1b and 1c).

**Figure 1.**
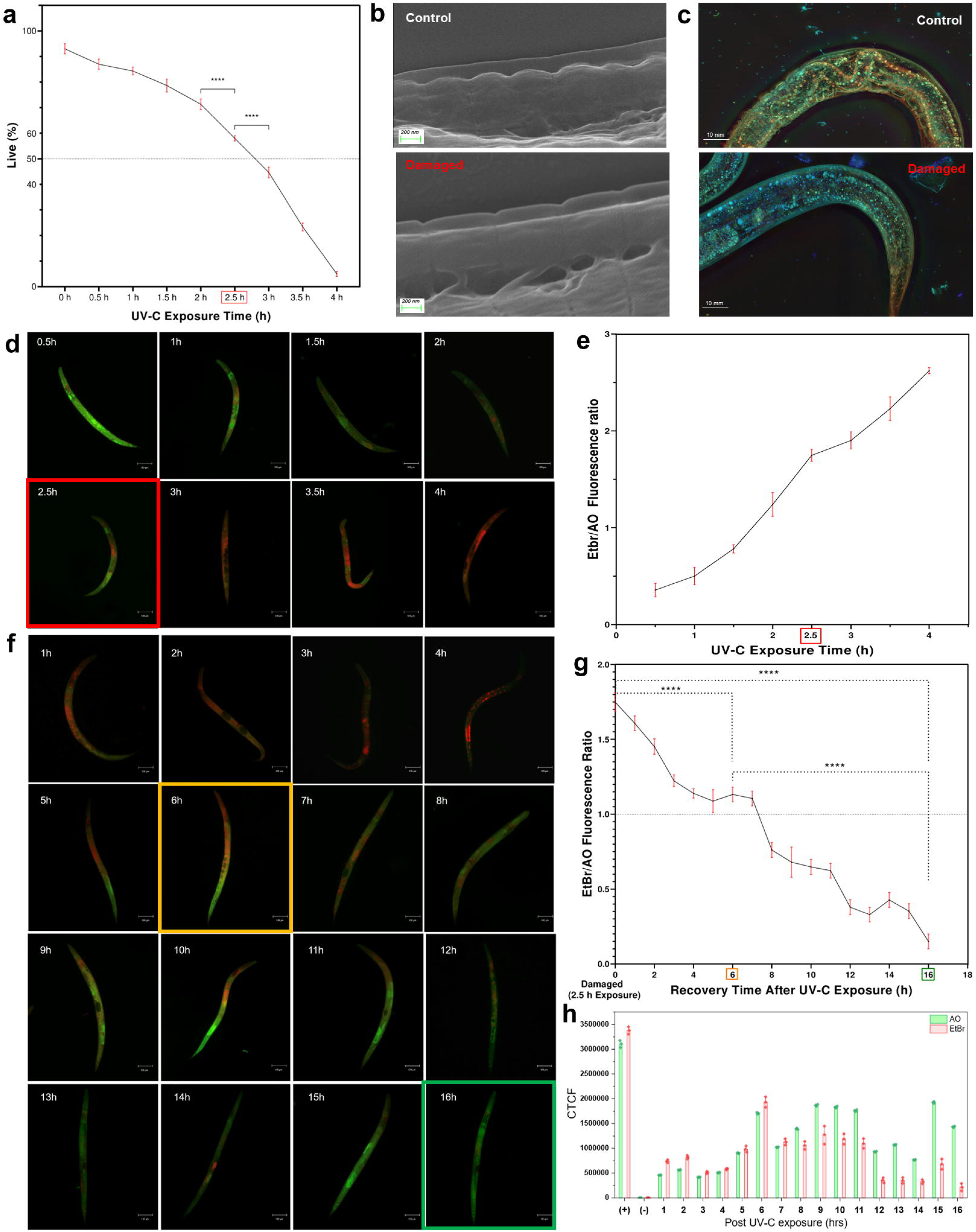
Phenotypic and morphological characterisation of UV-C-induced damage in *C. elegans*. **a**) Touch response assay showing live (%) worms during 4 hours of UV-C exposure (X-axis represents the time period of UV exposure, and the y-axis represents the percentage of live worms). **b**,**c)** Ultrastructure analysis of *C. elegans* epidermis via FESEM and Holotomography, respectively for Control and damaged (2.5 hours of exposure). **d)** Fluorescence microscopy image showing dual AO and EtBr-stained worms after every 30 mins of UV-C exposure (green represents viable cells, whereas red/orange represents dead cells) (scale bar = 100 μm). **e)** Graph showing EtBr/AO fluorescence with respect to continuous 4-hour exposure to UV-C. **f)** Fluorescence images indicating cellular recovery using dual AO and EtBr staining every hour post 2.5 hours of UV-C exposure. **g)** Graph showing EtBr/AO fluorescence ratio for dead and live cells post 2.5 hours of exposure. **h)** Graph showing AO and EtBr CTCF values measured at different post-UV-C exposure time points. (+), positive control; (−), exposed negative control.

We also studied tissue damage using dead and live cell staining with ethidium bromide (EtBr) & acridine orange (AO) dual staining at different time points. As AO binds to both live and dead cells and EtBr enters non-viable cells due to loss of their membrane integrity, we were able to observe uniformly green viable cell and orange/red apoptotic/necrotic cells. After 1.5 hours of UV-C exposure, EtBr-labeled cells were prominently observed. A significant increase in non-viable cells was observed after 2.5 hours (Figure 1d), and EtBr/AO ratio was ∼1.748 (Figure 1e). Followed by elevated dichromatic staining in worms at 3 hours post damage (Figure 1d). The positive control group used for the assay validation exhibited an increase in EtBr-associated red fluorescence relative to the control condition, confirming the EtBr and AO dual staining method. (Supplementary Figure S1a).

In a follow-up experiment, worms were continuously monitored over a period of 16 hours using fluorescence imaging and analysed using fiji/imageJ (Figure 1f). The integrated fluorescence intensity of AO and EtBr was measured for three worms at each one-hour interval from 1 hour to 16 hours, and the EtBr/AO fluorescence ratio was calculated as an indicator of the relative proportion of dead/live cells. The graph showed the ratio of EtBr to AO and corresponding recovery time points after a 2.5-hour exposure. The EtBr/AO ratio increased evidently post 2.5 hours of UV-C exposure, indicating enhanced cellular damage compared with control condition (non-UV-C exposure). Following UV-C exposure, at 0 hours (post-exposure), EtBr/AO ratio was ∼1.748 (Figure 1g); then the ratio progressively declined during subsequent response and repair phases. i.e. at 6 hours, this ratio decreased to around 1.132, and by 16 hours, dropped significantly to 0.15, suggested a gradual reduction in cellular damage and recovery of tissue integrity. Also, AO and EtBr CTCF profiles showed distinct temporal patterns following UV-C exposure, with the relative fluorescence signals changed substantially across the post-exposure period (Figure 1h).

Moreover, ROS levels were assessed to evaluate oxidative stress. The continuous UV-C exposure (0 to 4 hours) experiment demonstrated clear time-dependent increase in intracellular ROS levels in *C. elegans* (Figure 2a). Fluorescent dichlorofluorescein (DCF) increased progressively during initial period from 0.5 hours to 2 hours, followed by elevation after 2 hours. A noticeable increase in DCF was observed between 2 and 3 hours of continuous UV-C exposure, with fluorescence reaching ∼1.5 × 10 CTCF at 3 hours (Figure 2b). DCF levels continued to increase at 3.5 hours and reached the highest value at 4 hours, at ∼2.1 × 10 CTCF (Figure 2b). Overall, the progressive increase in CTCF indicated accumulation of intracellular oxidative stress with increasing duration of continuous UV-C exposure.

**Figure 2.**
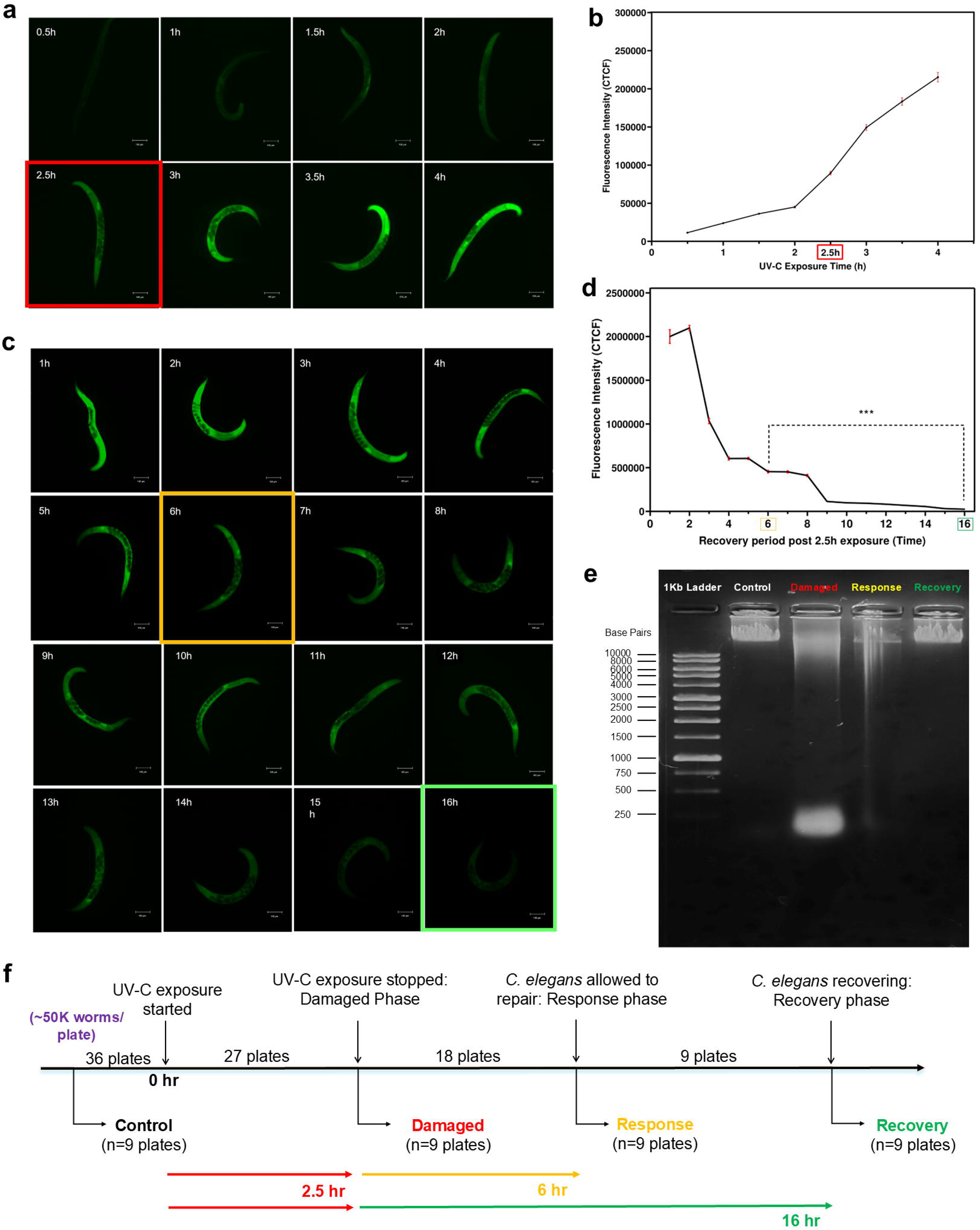
Oxidative stress and molecular characterisation of UV-C-induced damage in *C. elegans*. **a)** Representative DCF fluorescence images at the indicated time points following UV-C exposure. The 2.5-h time point is highlighted in red. Scale bars, 100 μm**. b)** Quantification of DCF fluorescence intensity corresponding to the time points shown in a. **c)** Representative DCF fluorescence images showing ROS activity at the indicated time points following the 2.5-h UV-C exposure. **d)** Quantitative analysis of DCF fluorescence intensity during the recovery period following 2.5-h UV-C exposure. *P* < 0.001. **e)** Agarose gel electrophoresis of control, damaged, response and recovery samples following UV-C exposure. **f)** Experimental workflow showing UV-C exposure and the collection of *C. elegans* samples at the indicated damage, response and recovery time points.

We also studied ROS activity after 2.5 hours of continuous exposure. Following cessation, DCF showed a progressive decline during the recovery period (Figure 2c). DCF intensity was highest during the early recovery phase (at 1–2 hours), ∼2.0–2.1 × 10 CTCF (Figure 2d), demonstrated substantial persistence of oxidative stress immediately following UV-C exposure. Subsequently, DCF fluorescence reduced sharply, reaching ∼1.0 × 10 CTCF at 3 hours and ∼6.0 × 10 CTCF at 4–5 hours (Figure 2d). Further, gradual decline was observed at 6-8 hours, followed by reduction at 9 hours. By 16 hours of recovery, decrease in DCF intensity level suggested a substantial change in UV-C-associated oxidative signal. The positive control group showed a clear increase in DCF-derived green fluorescence compared to control, confirmed the ability of the presence of elevated intracellular ROS (Supplementary Figure S1b).

Based on all experimental data, we classified samples into three distinct phases: i) “Damaged” phase, which refered to a continuous exposure to UV-C for 2.5 hours; ii) “Response” phase, which included samples taken 6 hours after exposure; and iii) “Recovery” phase, which comprises samples taken 16 hours post-exposure (Figure 2f). These time periods aligned with the recent study where comprehensive profiling of 12 time points (0.25 to 24 hours), portrayed three main events as Response (0.25 hours), Repair (0.5–4 hours), and remodelling (6–24 hours), 16 hours post-damaged (h.p.d) clearly falls within late remodelling stage where extracellular matrix (ECM) reconfiguration and epidermal function are restored [16].

Further, Genomic DNA integrity was assessed to determine UV-induced cellular damage. The control sample exhibited high-molecular-weight DNA retained close to loading well (right to 1kb ladder) (Figure 2e). However, damaged sample showed a diffused DNA signal and downward migration along the well. The response sample also demonstrated a smear extended from high-molecular-weight region to smaller molecular sizes, represented DNA fragmentation in the early post exposure phase. But remarkably, the recovery sample showed a high-molecular-weight DNA profile with reduced smear of low-molecular-weight DNA, which resembled control group pattern. These patterns in gel electrophoresis complemented AO/EtBr and ROS observations and supported transition from UV-C-induced cellular damage to recovery phase.

### Characterisation of *C. elegans-*derived sEVs

Dynamic light scattering (DLS) analysis revealed the hydrodynamic diameter of 126 nm (Figure 3a), and polydispersity index (PI) of 0.136. This showed that sEVs were monodisperse and had low variation in size. Subsequently, nanoparticle tracking analysis (NTA) revealed that the elution had a homogeneous population, and sEV concentration was 1.10±0.04×10^13^ particles/mL (mode:162.0 nm, mean:218.1 nm, D50:198.6±3.8 nm) (Figure 3b, Supplementary Figure S1e).

**Figure 3.**
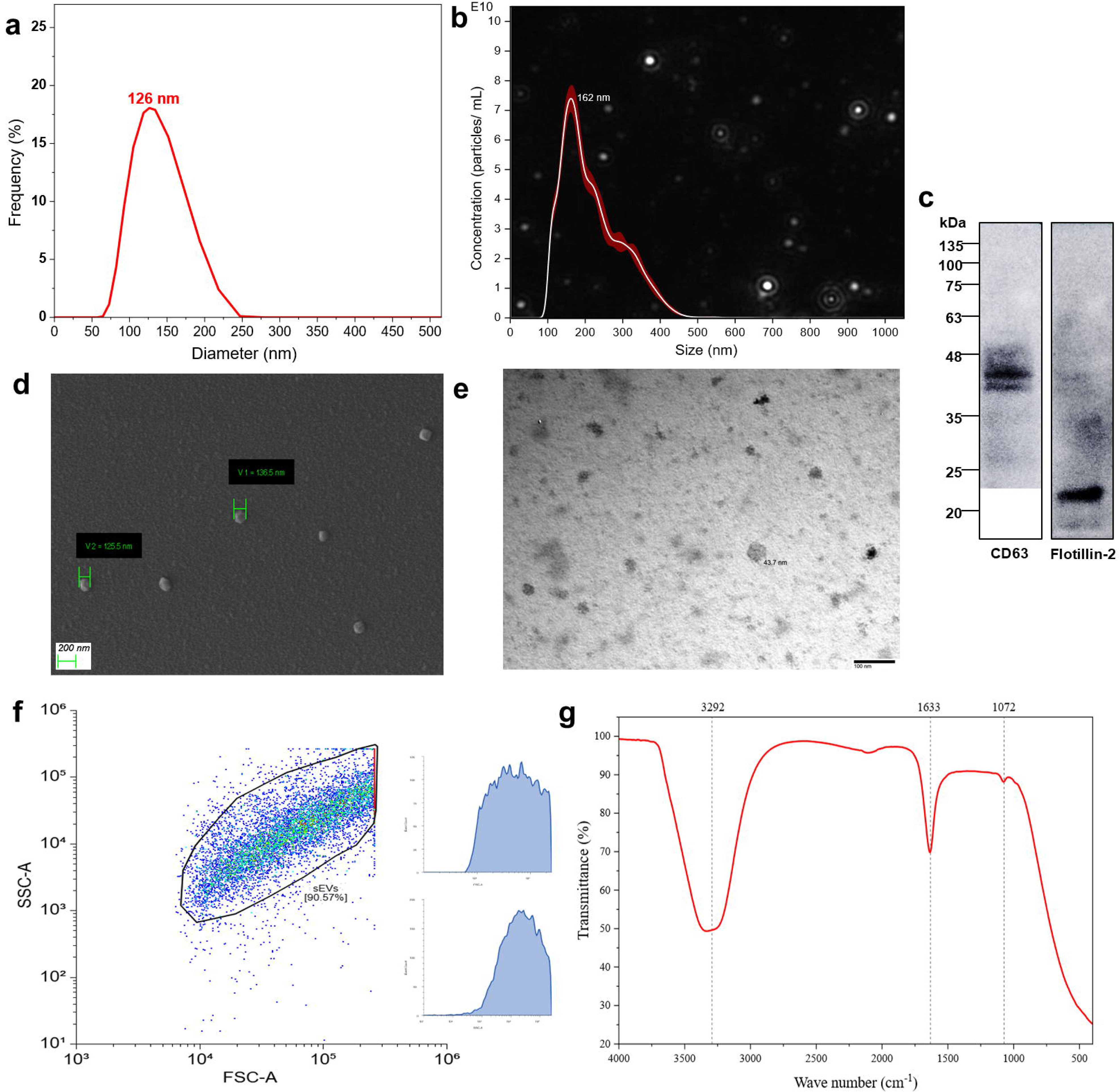
Characterisation of small extracellular vesicles (sEVs). **a)** Dynamic light scattering analysis (DLS) of sEVs showing 126 nm as the mean hydrodynamic diameter (x-axis = Diameter [nm]; y-axis = % frequency). b) Nano tracking analysis (NTA) of sEVs depicting 162 nm as the mean hydrodynamic diameter. The graph represents particle size (nm) on the x-axis and particle concentration (particles/mL) on the y-axis. **c)** Western blot analysis of sEVs with positive signals for CD63 (63 kDa) and anti-flotilin-2 (42 kDa). **d)** Ultrastructure of sEVs observed under FESEM (scale bar = 200 nm). **e)** TEM image of sEVs (scale bar = 100 nm). **f)** FSC vs SSC scatter plot of homogeneous sEVs population from flow cytometry analysis. g FTIR represents three distinct peaks at 3292 cm-1 [O-H, N-H], 1633 cm-1 [amide-I vibration from C=O stretching with a minor N-H bending reflecting mostly α-helical and the disordered secondary structures in membrane-associated, along with luminal proteins, a signal] and 1072 cm-1 [emerged from C-O stretching and C-O-C glycosidic linkages in polysaccharide, glycoproteins in addition to that PO₂⁻ asymmetric stretching from phosphodiester bonds in RNA or nucleic acid in the cargo in sEVs] (X-axis = wavenumber [cm-1] Y-axis = transmittance %).

Western blot analysis confirmed that sEV population had positive signals for CD63 (63 kDa) cell surface protein (tetraspanin) involved in vesicle transport, and anti-flotillin-2 (42 kDa) antibodies, a scaffolding protein associated with protein trafficking (Figure 2C, Supplementary Figure S1d).

Morphological analysis of by FESEM and TEM demonstrated that intact vesicles (∼110 nm to ∼140 nm) were present in concentrated elution (Figure 2d, e). Ultrastructure analysis also confirmed the cup-shaped double-layer vesicle.

A homogeneous sEV population was confirmed using flow cytometry analysis (10000 events). Post-gated analysis revealed that 90.57% of the population (9057 events) belonged to sEVs (Figure 2f).

Fourier-transformed infrared (FTIR) spectroscopy revealed biological composition of sEVs (Figure 2g). An absorption peak at 3292 cm^-1^ was attributed to O-H and N-H stretching vibrations, characteristic of hydrogen-bonded hydroxyl groups in carbohydrates or water and primary/secondary amines derived from proteins, which marks the hallmark of hydrated biomolecular structures in sEVs (Figure 2g). Absorption peak at 1633 cm^-1^ indicated the amide I vibration from C=O stretching with minor N-H bending, reflecting mostly α-helical and disordered secondary structures in membrane-associated, along with luminal proteins (Figure 2g). Similarly, signal at 1072 cm⁻¹ was observed, from C-O stretching and C-O-C glycosidic linkages in polysaccharide, glycoproteins. (PO₂⁻ asymmetric stretching from phosphodiester bonds in RNA or nucleic acids) in sEVs cargo.

Subsequently, Raman spectroscopy of sEVs revealed similar prominent peaks in the spectra. Peaks at 676 cm^-1^ correspond to C-S stretching in the proteins (methionine/cysteine) as well as guanine rings modes in nucleic acid (∼668-683 cm⁻¹) (Supplementary Figure S1f), 892 cm^-1^ specified for symmetric stretching of C-O-C in glycoproteins/ proteins (∼852-883 cm⁻¹ tyrosine/amino acids), and resonance peak at 977 cm^-1^ reflected C-C skeletal vibration in lipid-protein complexes and lipid vibrations near cholesterol regions (∼700-717 cm⁻¹). These results aligned with minimal information on sEVs for studies of extracellular vesicles guidelines[11]. Upon successful confirmation, isolated sEVs from all four experimental groups were subjected to miRNA and protein study. (Figure 4a).

**Figure 4.**
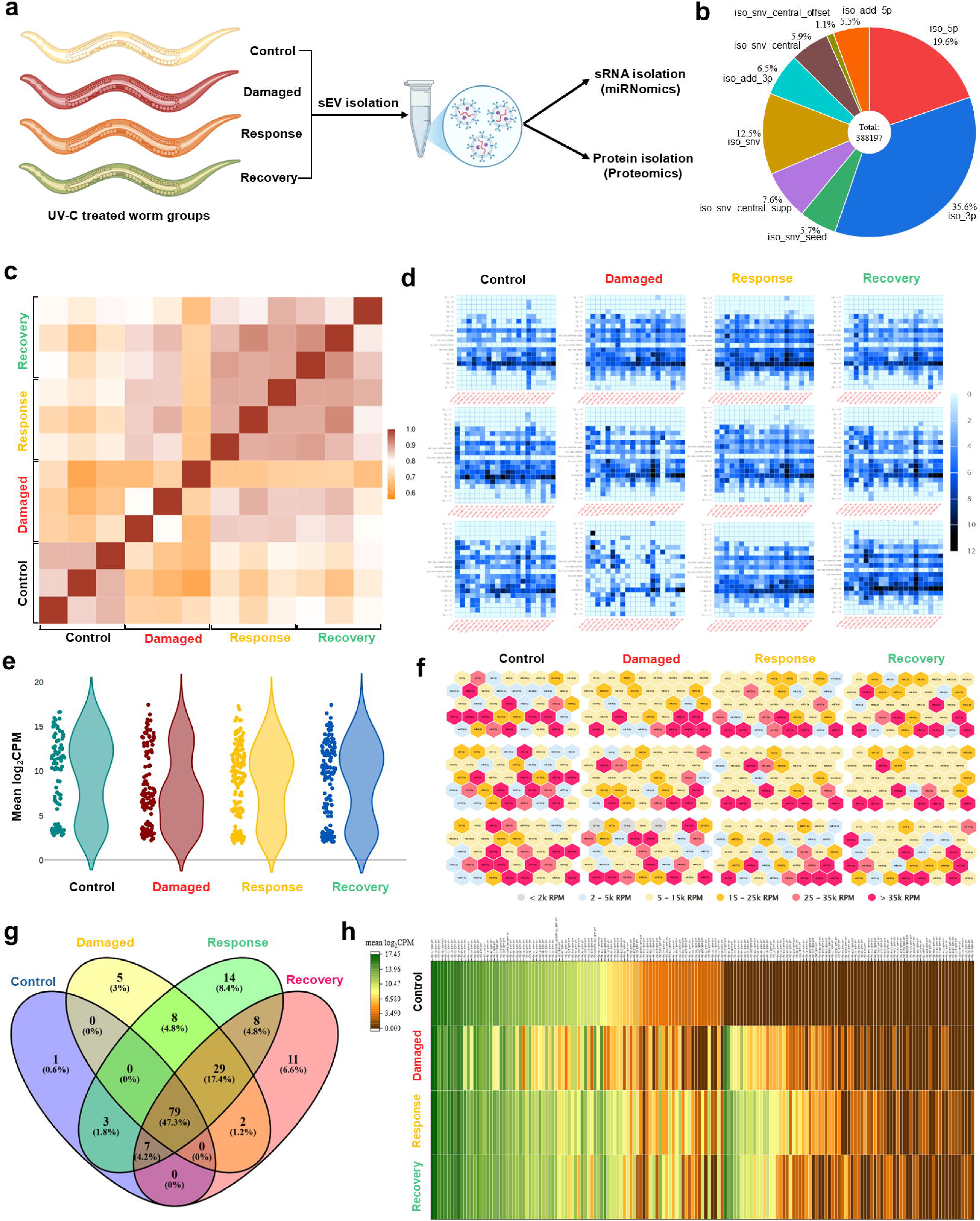
sEVs miRNA transcriptome profile post UV-C damage. **a)** Depiction of sample nomenclature used for sEVs transcriptomics analysis. **b)** Small RNA-Seq analysis of sEVs across all samples indicating extensive isomiR heterogeneity. **c)** Heatmap representing the correlation of expressed miRNAs across samples (colour gradient correlation level: brown [1] to orange [0.6]). **d)** The relative abundance heatmaps of isomiRs in the top 20 miRNAs. Represented isomiR types are canonical miRNAs, 5’ and 3’ truncations or additions representing length shifts at arm ends, iso_snv (central/central_supp/central_offset/seed) (scale bar: 0-12 of log2[read counts]). **e)** Violin plot illustrates expression distribution and statistical significance of miRNAs, highlighting dynamic shifts in expression (x-axis = sample; y-axis = mean log_2_CPM). **f)** Hexabin plot representing 40 abundant miRNAs in different samples; colour intensity represents the abundance in reads per million (Light grey for <2k RPM, light blue for 2-5k RPM, light yellow for 5-15k RPM, orange for 15-25k RPM, light pink for 25-35k RPM, and pink for >35k RPM). **g)** Venn diagram showing the number and percentage of expressed miRNAs in four different samples. **h)** Heat map of 167 differentially expressed miRNAs across four experimental groups (mean log_2_CPM colour gradient scale bar: 0 to 17.45).

### sEVs miRNA transcriptome profile

Small RNA-Seq analysis of sEVs revealed isomiR (isoforms of canonical mature miRNAs differ in sequence or length) heterogeneity, with the distribution of variant types as shown in Figure 4b. Out of 388197 reads, iso_3p contributed largest fraction (35.6%), followed by 5’-end variation of mature miRNAs (iso_5p,19.6%), Iso_snv (12.5%) and Iso_snv_central_offset (1.1%) (Figure 4b). These patterns were quantified by miRge3.0[17] and DESeq2[18], and represented dominant post-transcriptional modifications during sEV packaging. Expression patterns of miRNAs across three replicates were represented by a clustered heatmap (Figure 4c). The relative abundance heatmaps indicated variation among different types of isomiRs in the top 20 abundant miRNAs (Figure 4d, Supplementary file 2). The violin plot (density plot) showed the expression distribution of miRNAs, highlighting dynamic shifts in expression post-UV-C exposure (Figure 4e). The distribution patterns of control and damaged were relatively different, whereas compared to response and recovery groups were found similar. This represented the restoration of miRNA expression upon onset of repair phase (Figure 4e). The hexagon plot displayed normalised expression levels of 40 most abundant miRNAs (Figure 4f). Across our 12 hexagon plots, clear miRNA RPM patterns mark *C. elegans* recovery progression in sEVs. Control samples showed balanced miR-35 family/let-7 prominence with moderate miR-34/71, damaged phase triggered miR-34/71 explosion (>25k RPM, darkest tiles) while miR-35 shrinks, response sustained miR-34 peaks with miR-1/64, and repair was observed for partial return to control-like miR-35/1 but miR-34/71 stays elevated (>25k RPM) for matrix remodeling and scar resolution (Figure 4f, Supplementary file 3). Overall, miR-34/71 acted as conserved “stress-to-repair” hubs, dynamically reprogrammed EV miRNA cargo for intercellular signalling. The upset plot illustrated intersections of all miRNAs across four samples; herein, 79 miRNAs (47.3%) were common in groups; however, 1(0.6%), 5 miRNAs (3%), 14 miRNAs (8.4%) and 11 miRNAs (6.6%) were unique in control, damaged, response and recovery groups, respectively. A clustered heatmap of miRNAs (n=167 out of 431 mapped miRNAs) revealed differential miRNA expression post UV-C damage (Figure 4h). We observed significant variation between control and damaged groups. However, expression patterns in response and recovery were similar to the control group (Figure 4h), which complemented violin plot presentation (Figure 4e). Interestingly, a few sets of miRNAs with lower expression remained upregulated in a selected time frame of the recovery phase. This suggested memory association of miRNAs, and clustered heatmap complemented miRNA abundance and orchestration during the response and recovery phase.

### Differentially Expressed miRNAs (DE miRNAs)

Abundance analysis of phase-specific DE miRNAs demonstrated distinct temporal expression. The damaged phase exhibited elevated medians with broad right-skewed distributions, while response phase showed intermediate bimodal profiles (Figure 5a). Recovery samples showed baseline distributions, confirmed post-transcriptional recovery. Volcano plot analysis across damaged, response and recovery phases identified distinct miRNA regulatory signatures (log2FC 1.5 and - log_10_*Padj* 1.33) (Figure 5b). It showed that 28 (22 ↑ and 6 ↓), 43 (43 ↑ only) and 33 (33 ↑ only) miRNAs were present in damaged, response and recovery samples, respectively (Figure 5b). Out of total of 51 upregulated DE miRNAs, 18 miRNAs were found in all samples; however, we observed that 6 downregulated DE miRNAs were found exclusively in the damaged sample (Figure 5c).

**Figure 5.**
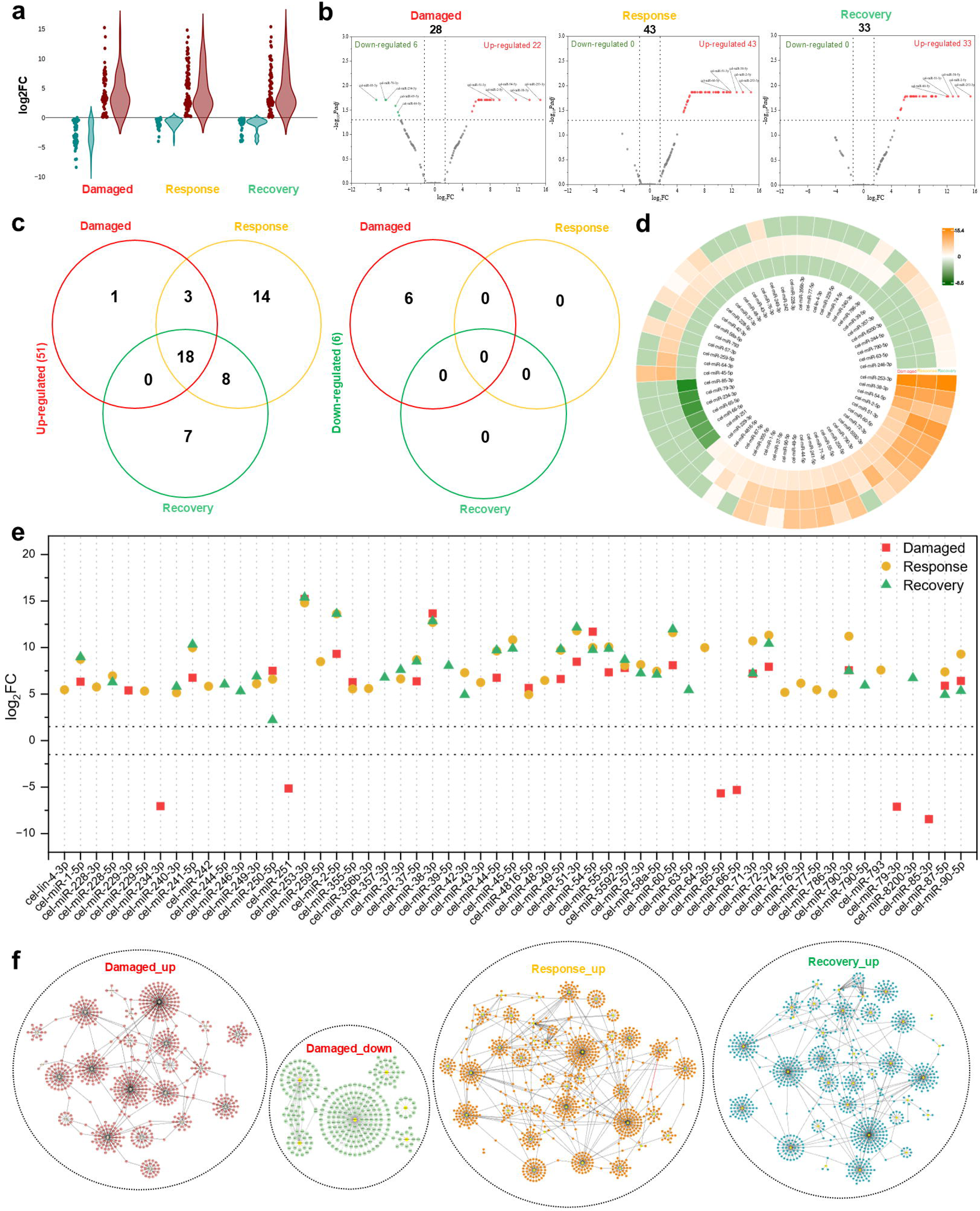
Differential microRNA expression analysis. **a)** Violin plots of log₂FC values for phase-specific differentially expressed miRNAs (n=57, *Padj* <0.05 and |log2fold change| >1.5). **b)** Scatter plot representation of differentially expressed miRNAs in damaged, response and recovery samples (cutoff, log2FC = ±1.5, - log_10_*Padj* = 1.33). **c** Venn diagram comparing the differentially upregulated and downregulated miRNAs. **d,e)** Circos plot and scatter plot illustrating the expression profile of differentially expressed miRNAs (scale bar=-8.5 to 15.4). **f** Target network analysis of miRNAs and their respective protein target; each miRNA is positioned at the centre, with surrounding nodes representing its target proteins.

We reported that only cel-miR-229-3p was upregulated in damage phase and previously associated with stress-response and longevity-related pathways in *C. elegans*[19, 20]. These 6 miRNAs have previously implicated in stress and heat-shock responses[21, 22] (cel-miR-85-3p, cel-miR-65-5p, and cel-miR-66-5p), neuronal migration[23] and neuromuscular signalling[24] (cel-miR-79-3p and cel-miR-234-3p), and innate immune responses[25] (cel-miR-251). We also reported that a subset of the differentially expressed *C. elegans* miRNAs had sequence relationships with human miRNAs, based on the criteria of Ibáñez-Ventoso et al.[26]. These sequence-based relationships and reported functions are summarised in Supplementary Table S5.

DE miRNAs shown in the circus plot and scatter plot clearly represented variability and complemented Venn analysis (Figure 5d,e). miRNA-target protein networks identified by miRanda and analysed in Cytoscape for phase-specific regulated and downregulated miRNAs, which generated potential hubs (Figure 5f; Supplementary figure S1h, Supplementary figure S1i). Hubs connected to ≥ 50 potential target genes in damaged_up, included cel-miR-60-5p, cel-miR-51-3p, cel-miR-5592-3p, cel-miR-54-5p, cel-miR-49-5p, cel-miR-241-5p, and cel-miR-2-5p (Figure 5f). Only two potent hubs (cel-miR-85-3p and cel-miR-66-5p) were found in damaged_down. cel-miR-241-5p, cel-miR-2-5p, cel-miR-43-3p, cel-miR-49-5p, cel-miR-51-3p and cel-miR-60-5p were found in response_up (Figure 5f). Additionally, recovery_up displayed cel-miR-60-5p, cel-miR-51-3p, cel-miR-54-5p, cel-miR-49-5p, cel-miR-241-5p, cel-miR-244-5p and cel-miR-2-5p as potential hub (Figure 5f).

### Temporal Profiling of sEV-Associated Proteins

We have also sequenced sEV-packaged proteins using LC-MS. The proteomic data were analysed using abundance values. sEV protein profiling revealed a distribution pattern across damaged, response and recovery samples of 3794 proteins (Figure 6a). Principal component analysis (PCA) exhibited strong clustering and distribution among samples (Figure 6b). Significant variability was observed between control and damaged samples (∼ 23%) and between damaged and response/recovery samples (∼35%). This showed relatedness between response and recovery samples (Figure 6b). Sample triplicates showed strong correlation and internal consistency (deep red cluster), and we also observed an inverse relationship (blue circle) between control and damaged/response/recovery samples (Figure 6c). DE proteins in sEVs were plotted using volcano plot (log2FC ≥ 1.5 and-log10*Padj* ≥ 1.33) (Figure 6d). Out of total 451 DE-damaged proteins, 229 (↓), 222 (↑) and top 20 were labelled in Figure 5d. The response phase (out of 116) showed 98 ↓ and 18 ↑ proteins. In addition, recovery phase (187 DE proteins) showed 148 ↓ and 39 ↑ (Figure 6d). We observed that response phase had lower number of DE proteins compared to recovery sample. Commonly and uniquely expressed proteins among three groups were identified using Venn diagrams (Figure 6e). Total of 556 DE proteins (249 ↑ & 307 ↓) were found across samples, out of which 200, 6 and 18 proteins were upregulated in damage, response and recovery, respectively. Similarly, 116, 23 and 37 proteins were downregulated in damage, response and recovery groups, respectively. We have also observed 5 ↑ and 37 ↓ proteins common to all samples. The resulting *C. elegans*–human orthologous protein pairs were presented in Supplementary file 4. Further, differentially regulated proteins of the damage, response and recovery groups were categorised into biological processes (BP), cellular components (CC) and molecular functions (MF) for gene ontology (GO) (significant threshold:-log10(*Padj*) ≥ 1.301) (Figure 6a). GO: BP enrichment revealed a stage-dependent shift in biological processes post UV-C exposure. The damaged stage was predominantly associated with multicellular development, anatomical organisation, and cellular developmental processes, whereas the response stage showed enrichment of locomotion, cell motility and developmental-regulatory processes. Recovery-associated proteins were mainly linked to multicellular growth, developmental processes, anatomical organisation and cell-fate regulation (Figure 7a). Overall, these patterns indicated progressive cellular remodelling, involvement of calcium signalling, ROS and cytoskeletal changes in *C. elegans*[27–29].

**Figure 6.**
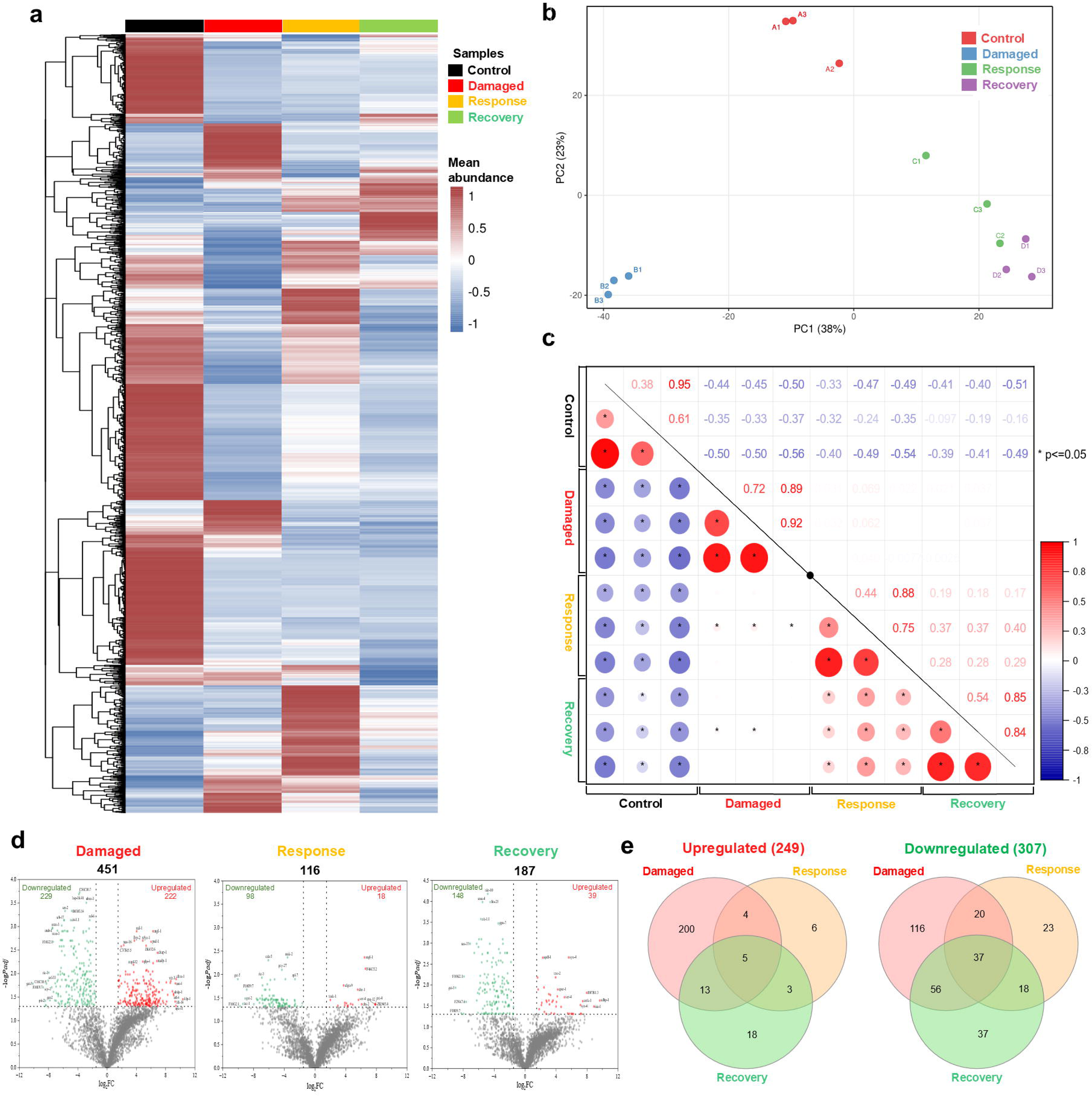
Differential protein expression analysis. a,b) Sample-wise PCA distribution and correlation analysis of proteomic data (p <= 0.5). **c)** Heatmap representation of mean abundance values of differentially expressed proteins (scale bar=-1 to 1). **d)** The volcano plots showing 451, 116 and 187 differentially expressed miRNAs in the damage, response and repair phases, respectively (cutoff, log2FC = ±1.5,-log_10_*Padj* = 1.33). A Venn diagram comparing the differentially upregulated and downregulated proteins.

**Figure 7.**
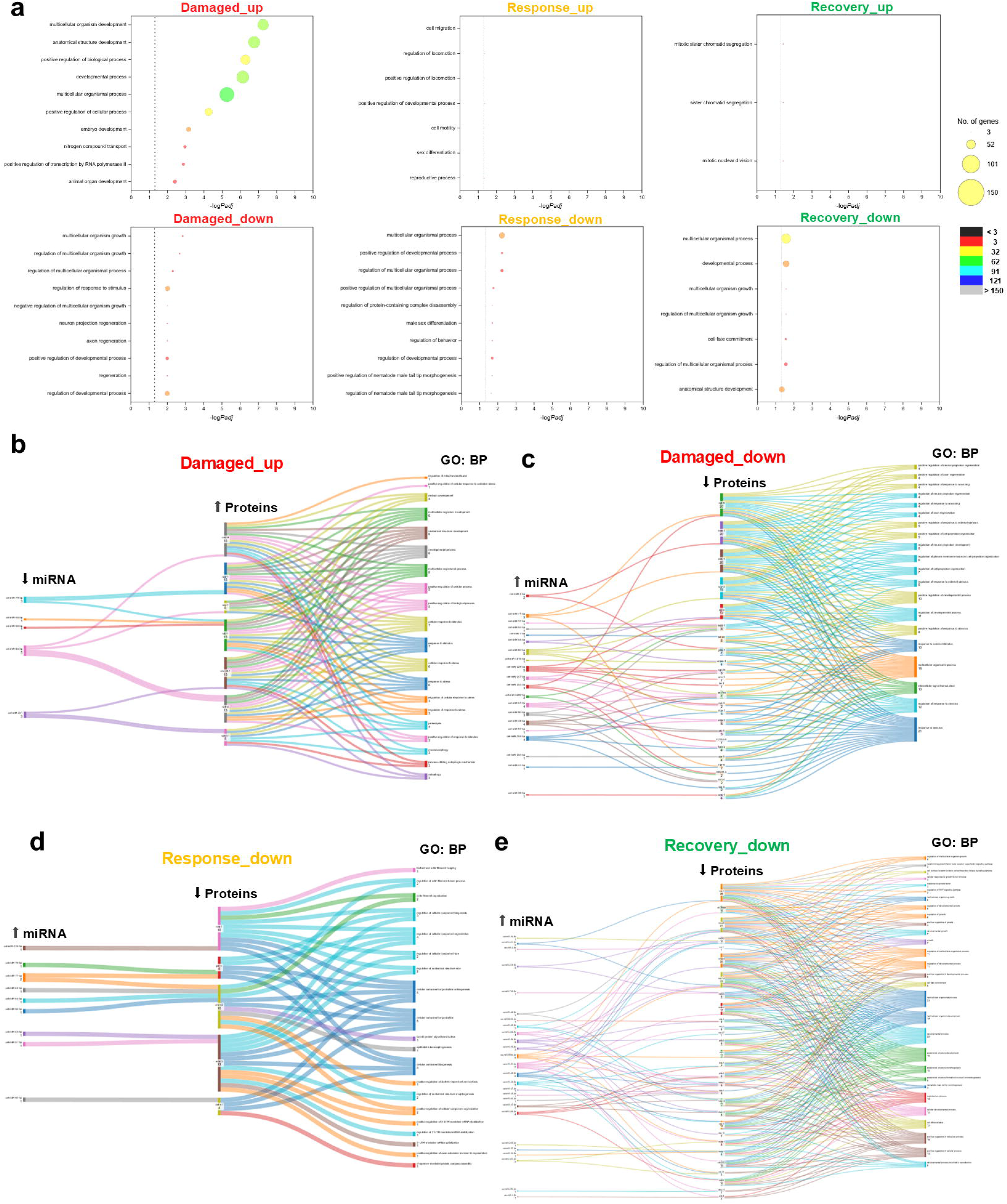
Enrichment analysis of differentially expressed proteins. **a)** Gene ontology (GO) enrichment analysis of differentially expressed proteins in the damage, response and repair phases. The x-axis represents-log*Padj* value, and y axis represent the cellular process. **b)** Sankey diagram of relationships between miRNA and upregulated proteins in the damage phase, **c)** miRNA and downregulated proteins in the damage sample, **d)** miRNA and downregulated proteins in the response phase, **e)** miRNA and downregulated proteins in the recovery phase. The heights of the coloured boxes and connecting bands represent the relative numbers of proteins and biological pathways regulated by a particular miRNA.

### Inverse miRNA-Protein analysis

To integrate temporal changes in miRNA and protein abundance, inverse analysis was performed to identify associations across damage, response, and recovery samples (Figure 7b-e, Supplementary Figure S1j). Downregulated miRNAs from damaged samples (cel-miR-79-3p, cel-miR-85-3p, cel-miR-65-5p and cel-miR-66-5p) were associated with increased abundance of stress/response-related proteins (SKN-1, CED-9 and SVH-2) (Figure 7b). Upregulated miRNAs from damaged sample (cel-miR-71-3p, cel-miR-60-5p, cel-miR-229-3p, cel-miR-253-3p and cel-miR-51-3p) were associated with decreased abundance of EGL-8, MEK-1 and MLK-1 (Figure 7c). Upregulated miRNAs from response sample (cel-miR-51-3p, cel-miR-228-3p and cel-miR-77-5p) were associated with reduced abundance of CDC-42, PFN-1, and MAK-2, proteins linked to cellular and cytoskeletal remodelling (Figure 7d). Similarly, upregulated miRNAs from recovery sample were associated with reduced abundance of KIN-29, SMA-4 and ELT-2 proteins, reflecting changes in growth/development/stress-associated protein networks (Figure 7e). Overall, Sankey diagram revealed a temporal shift in miRNA–protein association.

### Pseudotime trajectory analysis

Based on pseudotime trend profiles (Figure 8b-f), proteins from individual clusters were enriched for top 15 *C. elegans* phenotype ontology annotations using STRING database[30], integrated from MONARCH initiative[31] and presented in the right panel (Figure 8).

**Figure 8.**
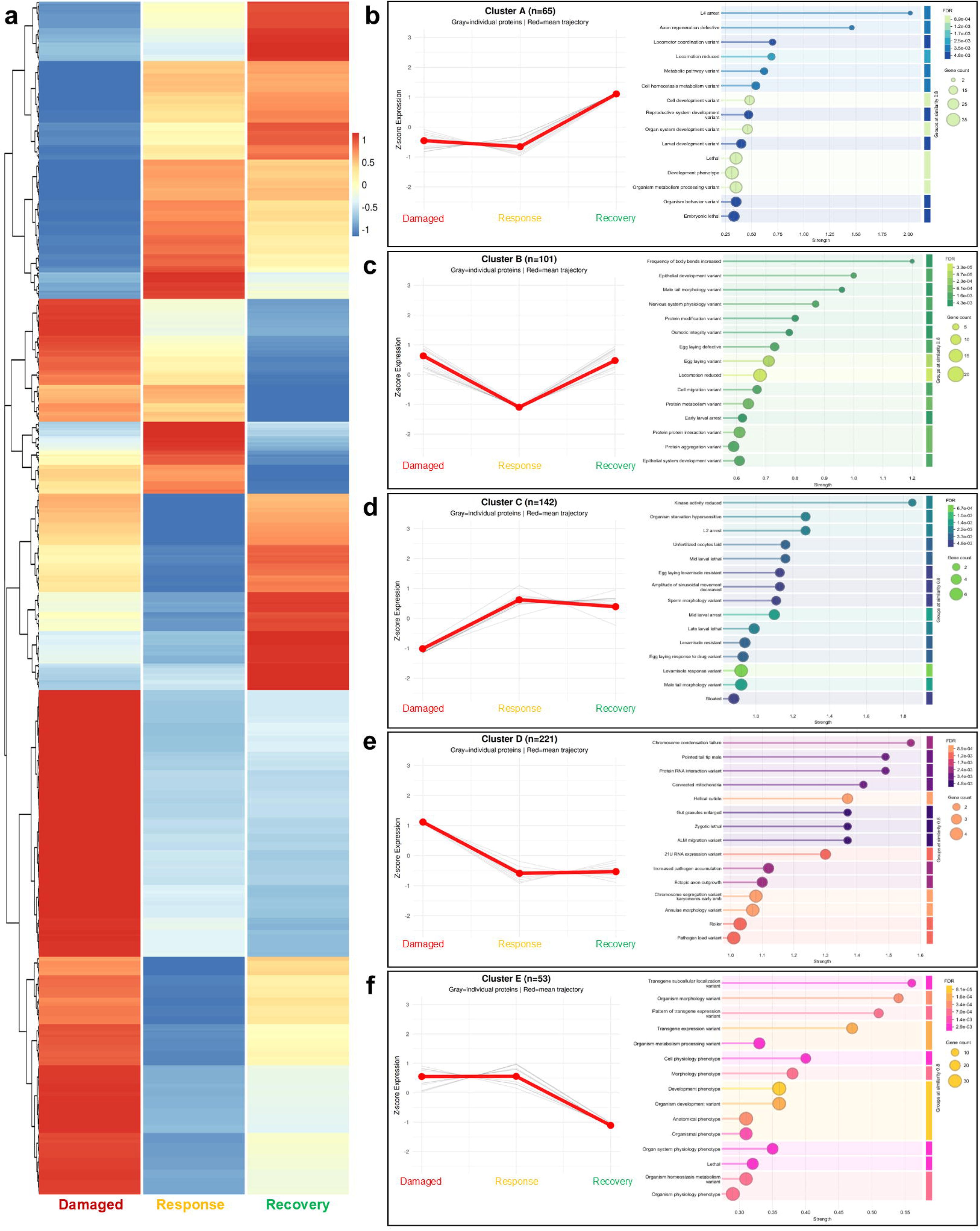
Differential protein expression profiles along tissue recovery. **a)** Protein expression heatmap of 582 DEPs (q-value <0.05) across the different samples was clustered hierarchically into five clusters. The colour indication from blue to red indicates relative expression levels from low to high. **b-e)** Pseudotime analysis of protein expression dynamics was classified into 5 clusters (A-E). The grey line represents the individual protein trajectory, whereas the red line represents the mean trajectory of the total proteins in each cluster.

Cluster A: Late upregulation during the Recovery phase.

The protein cluster A (n=65) in this pattern corresponds to early and mid-downregulation with upregulation in recovery phase. Axon regeneration defects, cell development variants, and lethal and developmental phenotypes were major contributors to the monarch phenotype study (Figure 8b).

Cluster B: Biphasic expression with mid-response suppression.

A sharp decline in the response phase compared to damage and repair phase was contributed by n=101 in cluster B. Locomotion reduction, cell migration, and protein aggregation variants were prominent pathways observed in this cluster (Figure 8c). Cluster C: Initial repression followed by sustained response-to-repair activation. (n=142)

The damage phase showed downregulation of proteins and consistent upregulation in later response and repair phase for homeostasis. In this case, one of the notable pathways observed in the damage phase was reduced kinase activity, which results in accelerated ageing and shortened lifespan (Figure 8d).

Cluster D: Early activation with progressive mid-to-late downregulation (n=221). Regulated protein clusters in damaged/response/recovery phase were crucial for survival and homeostasis. It was observed that chromosome condensation failure was regulated, where condensin subunits (smc-4, smc-5, and mix-1) might be increased to handle replication stalling upon UV exposure. Helical defects in response to environmental stressors cause upregulation of collagen genes in damaged state before recovery phase (Figure 8e).

Cluster E: Early-to-mid expression peak with late-phase decline. (n=53)

These protein clusters showed major contributions in different pathways by upregulation in damage and response phase (mid transition) but downregulation in recovery phase. These included organismal phenotypes (life span/motility), developmental phenotypes (worm stage delays), cell physiology (apoptosis/stress), and anatomical phenotypes (germ cell/growth defects) (Figure 8f).

### Cross-Species Comparison of miRNAs and Human Orthology of Proteins

Cross-species sequence comparison reported human sequence relationships for 57 differentially abundant *C. elegans* miRNAs. Supplementary Table S5 summarises the corresponding sequence relationships and reported biological associations. Similarly, human orthology analysis of DE proteins identified corresponding human orthologs for a substantial subset of the *C. elegans* proteins. Multiple human orthologs were identified for several *C. elegans* proteins, reflecting one-to-many orthologous relationships and resulting protein orthology was summarised in Supplementary file 4.

### Orthogonal validation using Stem-loop RT-qPCR

To validate miRNA sequencing data, we selected 10 phase-specific miRNAs from DE miRNAs and 1 from non-DE miRNAs. Figure 9a shows the selection strategy of miRNAs for stem-loop RT-qPCR. Compared to control, we observed upregulation of cel-miR-253-3p, cel-miR-85-3p, cel-miR-2-5p, and cel-miR-250-5p in all samples (Figure 9b). Additionally, cel-miR-79-3p, cel-miR-90-5p, cel-miR-794-5p, cel-miR-249-3p, and cel-miR-60-5p were downregulated in damage but upregulated in response and recovery phases (Figure 9b). Significant correlation was observed between RNA-seq (normalised miR count) and stem-loop RT-qPCR results (2^-ΔΔCt^) (Figure 9c, melting curve Supplementary Figure S1k). A mean correlation coefficient value among compared groups was 0.54 (±0.15), indicated a positive relation (Figure 9c).

**Figure 9.**
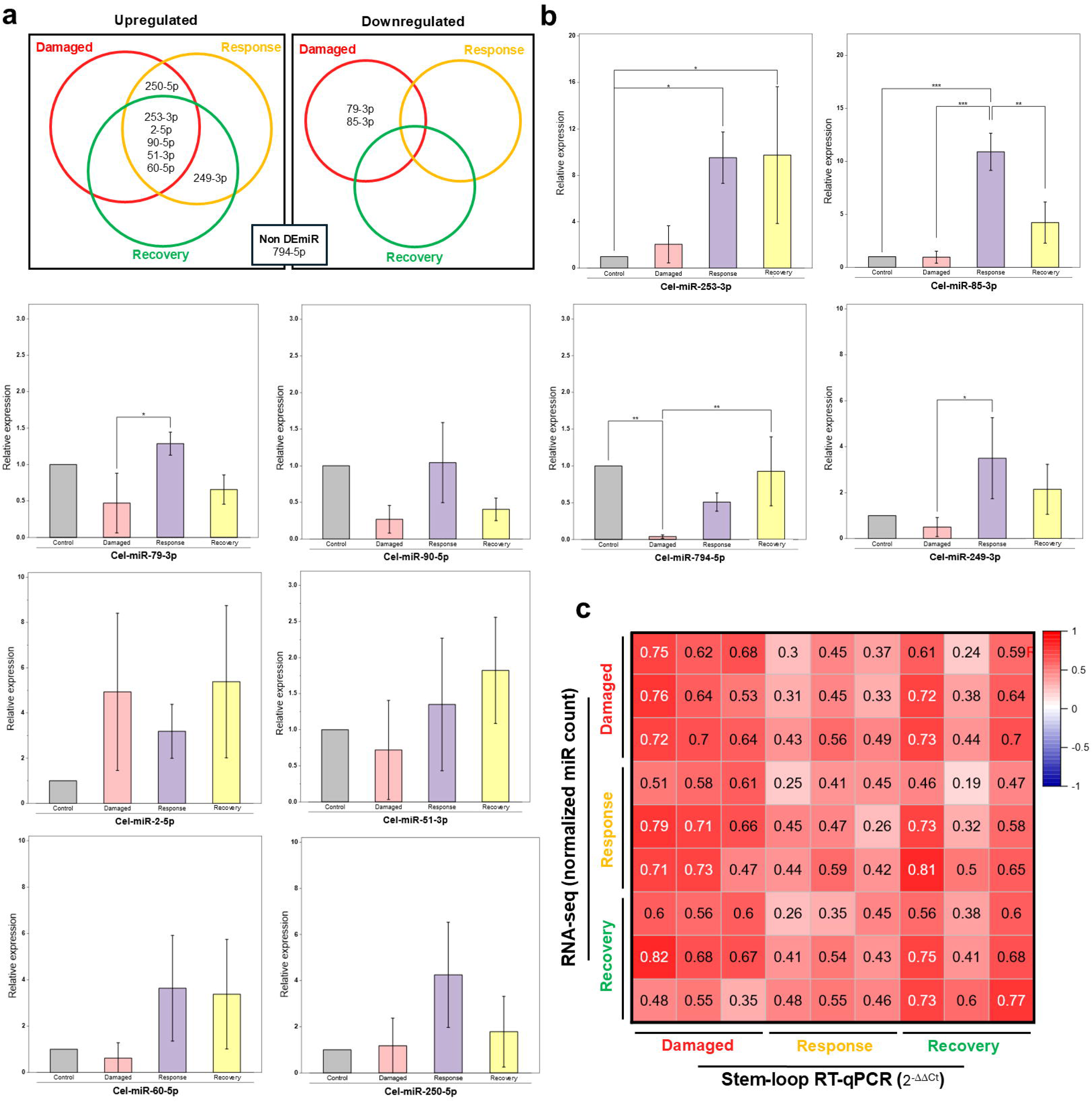
Stem-loop RT-qPCR validation. **a)** VENN diagrams representing phase-specific DE miRNAs and a list of non-DE miRNAs selected for orthogonal Stem-loop RT-qPCR validation. **b)** Stem-loop RT-qPCR validation of 10 miRNAs across all the samples, where the Y-axis represents their relative expression (2^-ΔΔCt^) and the expression of Control is always 1. One-way ANOVA has been performed across the four groups of each miRNA, followed by Tukey’s post-hoc pairwise test comparing each group to the control. Significant pairs have been marked with horizontal lines having scale (* p<=0.05 * p<=0.01 ** p<=0.001). **c)** Correlation plot of 10 miRNAs between their RNA-seq miRNA values (Normalized read counts of damage, response and Recovery in triplicates) and Stem-loop RT-qPCR relative expression values (2^-ΔΔCt^ of damage, response and Recovery in triplicates). Correlation values have been reported in the plot on a scale of-1 (blue) to 1 (red).

## Discussion

UV-C exposure induces a temporally coordinated molecular response involving cellular stress, adaptation and recovery. Previous studies in *C. elegans* have shown that Ca²⁺-dependent signalling, cytoskeletal remodelling, and membrane repair are essential during epidermal wound closure. Time-resolved transcriptional profiling has demonstrated distinct stages of response, repair, and remodelling[16, 28]. However, temporal changes within sEVs cargoes during these transitions remain uncharacterised. In the present study, we profiled miRNA and protein abundance in sEVs obtained from UV-C-exposed *C. elegans*. The distinct molecular profiles observed in control/damage/response/recovery samples provided a temporal framework for characterising molecular remodelling. We identified 57 differentially expressed miRNAs, comprising 51 upregulated and 6 downregulated miRNAs. Among these, we found that the distinct miRNAs identified in our analysis had also been reported earlier. Such as, cel-miR-253-3p has previously been reported in dietary-restriction-related molecular remodelling in *C. elegans*[33]. In addition, cel-miR-85 has been experimentally implicated in the recovery phase post-heat shock regulation via HSP-70[21]. The progressive increase was also observed for cel-miR-51-3p, as reported earlier for embryonic development and maintenance of pharyngeal attachment[34]. Together, these distinct patterns indicated that miRNA response was dynamically remodelled. Along with three miRNAs discussed above, several additional DE miRNAs identified in the present study have previously been associated with stress responses, longevity, development, immunity, reproduction and metabolic regulation in *C. elegans*. These include cel-miR-71, cel-miR-65/66, cel-miR-79, cel-miR-85 and cel-miR-229/228, which have been reported in processes including stress adaptation, proteostasis, developmental timing, innate immunity and longevity[19, 21, 22, 35]. Collectively, these functions provided a biological context for the broader miRNA functioning observed post UV-C exposure. A detailed summary of the reported functions for these DE miRNAs is provided in Supplementary Table S5. Several temporally regulated *C. elegans* miRNAs showed previously reported sequence relationships with human miRNAs, providing an evolutionary context for the observed temporal responses (Supplementary Table S5). This comparison complements the temporal profiling by placing selected miRNAs within a broader cross-species sequence context. However, sequence similarity alone does not establish functional orthology, conserved target regulation, or equivalent biological activity; these relationships therefore provide comparative context rather than evidence of conserved function. miRNAs affect protein expression and a variety of cellular processes by regulating target transcripts in a sequence-dependent manner. Therefore, during cellular stress and recovery, stage-specific changes in miRNA abundance may coincide with corresponding changes in the proteome landscape.

Proteomic data showed a complementary temporal pattern, with 451, 116, and 187 differentially expressed proteins identified during the damage, response, and recovery stages, respectively. During the damage phase, altered proteins were predominantly associated with cellular and oxidative stress responses, autophagy, macroautophagy, proteolysis and Ca²⁺-dependent signalling (Figure 7b). These results were in accordance with the established importance of Ca²⁺-dependent signalling and cytoskeletal responses during tissue injury in *C. elegans*[28]. As the response progressed, the proteomic profile shifted toward cellular organisation and structural remodelling, including actin filament organisation, CDC42-associated signalling and endocytosis (Figure 7d). Representative proteins included CDC-42, PFN-1, CAP-1, MAK-2 and DAF-41, linking this stage to cytoskeletal organisation and cellular remodelling.

During the response-associated stage, CDC-42 provides a relevant biological context because injury-induced CDC-42 clustering promotes recruitment of WSP-1 and actin polymerisation[36]. PFN-1, a profilin involved in actin dynamics, further supports the association of this stage with cytoskeletal organisation (Figure 7d). These findings are consistent with a transition from an early stress-associated state toward structural and cellular remodelling. However, reported biological processes based on proteomic profiling do not imply that the detected proteins originate from a specific tissue or directly mediate repair.

During the recovery-associated stage, the proteomic profile shifted toward multicellular growth, development, morphogenesis, differentiation and growth-factor-associated processes. The recovery network included KIN-29, SMA-4, ELT-2, LON-1, EIF-2β, TIN-9.2, TSP-14, NEKL-3, APH-2, MIG-14 and CDC-25.2. These proteins were reported to show changes in metabolic, developmental, and cellular organisation activities (Figure 7e). Rather than indicating a complete return to the control expression level, these changes suggest continued molecular remodelling at 16 h. Therefore, this time point is more appropriately described as a recovery-associated stage rather than a completely repaired state.

Integration of the miRNA and protein datasets using Sankey analysis further highlighted stage-specific inverse associations. During damage, these associations included stress-and signalling-related proteins such as SKN-1, CED-9, SVH-2, EGL-8, MEK-1 and MLK-1; during response, CDC-42, PFN-1, CAP-1, MAK-2 and DAF-41; and recovery sample included LON-1, EIF-2β, KIN-29, SMA-4, TIN-9.2, TSP-14, NEKL-3, APH-2, ELT-2, MIG-14, CDC-25.2 and CBD-1. SKN-1 is well established as a regulator of oxidative stress defence and detoxification in C. elegans, providing a biological context for its representation within the damage-associated network[37]. Similarly, the temporal involvement of cytoskeletal proteins during the response stage is consistent with established wound-response biology[36]. However, inverse analysis identifies candidate temporal associations rather than direct miRNA–protein targeting or functional regulation.

A limitation of the current study is that the sEVs were obtained from whole worms. Consequently, the identified miRNAs and proteins cannot be assigned to specific tissues or cell populations, and their abundance cannot be interpreted as evidence of tissue-specific repair. In addition, the present workflow characterises sEVs and does not establish that all detected molecules represent bona fide intra-vesicular cargo or their cellular origin. Thus, the present findings provide a temporal molecular resource, describing changes in sEVs composition (miRNA and protein) associated with progression from UV-C-induced damage through response and recovery. Future studies incorporating tissue-specific sEV characterisation, EV uptake analysis, and functional manipulation of candidate miRNAs and proteins will be conducted to determine their functional relevance.

## Conclusion

Overall, this study provides a temporal profile of miRNA and protein abundance in sEVs, post UV-C exposure in *C. elegans*. The data revealed progression from damage (stress/cellular homeostasis changes) to response (through cellular and structural remodelling) to recovery (toward growth, developmental, and morphogenesis-associated changes). The temporal behaviour of selected miRNAs and associated proteins provides candidate molecular signatures for damage-to-recovery transitions.

## Materials and methods

### UV-C-induced damage in *C. elegans*

For this experiment, the wild-type *C. elegans* N2 strain (Bristol) was used. Worms were cultured in high-growth media at 22℃ and supplemented with *Escherichia coli* OP50 strain as per WormBase[38]. Homogeneous culture was used for experiments conducted in this paper. For induced damage, two UV-C tubes were set at 30 cm above worm plates at 22℃. UV-C tube (λ=254 nm, L-GPH287T5L, ARK Lite, India) was used for damage induction (0.728 mW, measured by an optical Power Monitor, THORLABS).

To study the effect of UV-C on *C. elegans*, a touch-response assay was conducted using a heated platinum-nichrome wire. It was placed near head region of worms at intervals of 30 minutes from 0 to 4 hours. If any movement was observed, classified as live; if not observed, then dead. The final count of live and dead worms was used to calculate live percentage (n =100, in triplicate), providing a quantitative measure of the damage.

Following touch response assay, cuticle morphology was observed using FESEM and label-free holotomography. Worms without UV-C exposure (Control) and worms with 2.5 hours of exposure (Damaged) were collected from plates and washed with M9 buffer to remove residual medium and bacteria. Randomly, few worms were picked by worm picker from both sample group and processed for imaging. For FESEM, gold-sputtered worms were used (magnification: 50.13 KX, WD: 5.3 mm at 15 kV) (JSM 6610LV, JEOL, Japan). In holotomography, live worms in M9 buffer were transferred to imaging chamber and positioned within field for acquisition. The Tomocube HT-X1™ (Tomocube Inc., Daejeon, South Korea) system was used to acquire multiple holographic projections at different illumination angles. These projections were combined and computationally reconstructed to generate 3D refractive-index tomograms.

Live & dead cell imaging was performed using dual AO-EtBr staining for a period of continuous 0 to 4 hours of exposure, as well as a subsequent recovery period lasting from 1 hour to 16 hours after 2.5 hours of exposure. For this, 50 µL of AO (10 mg/ml) and 50 µL of EtBr (10 mg/ml) were mixed. AO-EtBr stain was added to microcentrifuge tube containing ≅ 100 worms and incubated on a gel rocker for 30 minutes at 22℃ to ensure gentle mixing. Stained worms were collected, washed, and placed under a fluorescence microscope for imaging (EVOS™ M5000 Imaging System, Thermo Fisher Scientific, United States). A positive control group was also included, prolonged period of incubation with H_2_O_2_ was used to induce cell damage. Images were acquired for AO in GFP channel (10X magnification, GFP, λ_ex_= 482 nm, λ_em_= 524 nm) and for EtBr in RFP channel (10X magnification, RFP, λ_ex_= 531 nm, λ_em_= 593 nm). We used corrected total cell fluorescence (CTCF) for each channel and then calculated the EtBr/AO ratio as mentioned:

The final metric we used for the graph was:

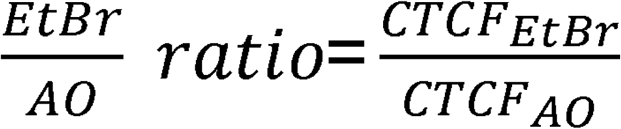

Where,

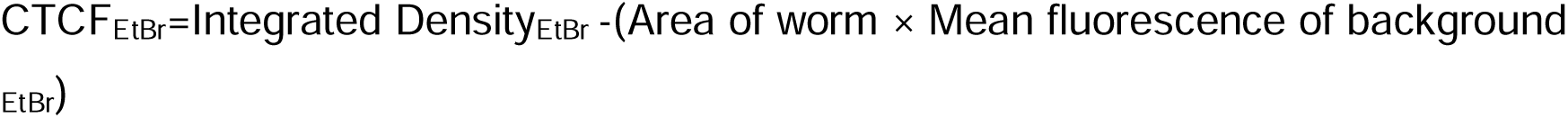

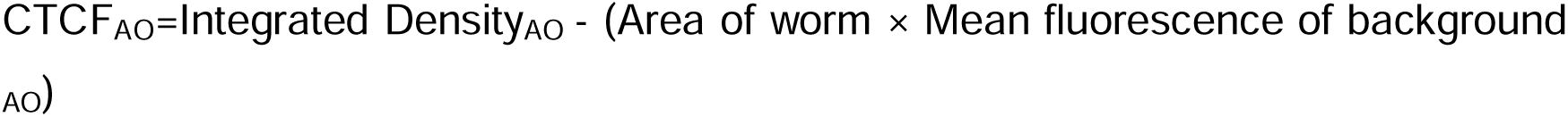

Following EtBr/AO based assessment of cellular damage, intracellular oxidative stress was evaluated by quantifying ROS using 2′,7′-dichlorodihydrofluorescein diacetate (H2DCFDA) (Cat no. D399, Invitrogen, Thermo Scientific, United States).H₂DCFDA (Non-Fluorescent), which undergoes deacetylation by cellular esterases and subsequent oxidation by reactive species to form highly fluorescent dichlorofluorescein (DCF). A 10 mM H2DCFDA stock solution was prepared in DMSO, and a 50 μM final working solution in M9 buffer was prepared. Worms were exposed to UV-C for overall ROS measurement by DCF. Collected worms at different time points, washed and incubated with 200 µL H2DCFDA (50 µm) for 20 min in the dark, then washed with M9 buffer to remove excess dye. Resulting green fluorescence images were acquired using the GFP channel (10X magnification, GFP, λ_ex_= 482 nm, λ_em_= 524 nm), keeping identical settings for all experimental groups. Positive control was also included using H_2_O_2_.

The fluorescence images were analysed using ImageJ/Fiji by selecting entire worms as regions of interest (ROI) and measuring area, Mean grey value, and Integrated Density (IntDen). ROS-associated green fluorescence intensity was quantified using the corrected total cell fluorescence (CTCF) formula used in the AO & EtBr experiment and plotted on a graph (Figure 2b & 2d).

CTCF = Integrated Density-(Area of worm × Mean fluorescence of background)

To further evaluate the extent of UV-C-induced cellular damage at genomic level, DNA isolation of worm groups was done using the organic liquid-liquid extraction technique with phenol: chloroform: isoamyl alcohol (PCI). For this, samples were collected from plates in M9 buffer and centrifuged at 2000 rpm for 2 mins at RT for bacterial washing. Worm lysis buffer was added to pellets in 3:1 v/v (volume/volume). Mixed by gently vortexing 10-20 times for every. Following, 40 µL of Proteinase K and 20 µL of RNase A were added, incubated at 65℃ for 60 mins and mixed every 10 mins during incubation. Followed by adding 500 µL PCI (25:24:1) and centrifuging at 12000 rpm for 10 mins at 4℃. Aqueous phase was collected and transferred into new vial with 3 M sodium acetate and pre-chilled absolute alcohol (centrifuged at 12000 rpm for 10 mins at 4℃). Pellet was washed with 70% ethanol and centrifuged at 12000 rpm for 10 mins at 4℃, discarded the supernatant and air-dried the pellet. 50 µL of Tris-EDTA buffer was added for elution. DNA isolated at four different time points was loaded onto a 0.8% agarose gel at equal concentration along with a 1 kb ladder and run for 2 hours at a constant 60 V, and visualized (Amersham Image Quant 800 Cytiva, Uppsala, Sweden).

For miRNA and protein analysis, 36 worm plates (n≅50k worms/plate) were assigned to: control, damage, response, and recovery (Figure 2f). From this, 27 plates were exposed to UV-C light as mentioned in the methods section. Samples were collected at defined time points following UV-C exposure. Nine plates each were harvested at four stages: untreated controls, damaged samples (after 2.5 hours of exposure), response samples (6 hours post-exposure) and recovery samples (16 hours post-exposure) (Figure 2f). Following UV-C exposure, plates were maintained under optimal growth conditions.

### Small extracellular vesicles (sEVs) isolation

Our previously published protocol[39], with minor modifications, was used to isolate sEVs from *C. elegans*. In brief, worm pellets were sonicated on ice using probe sonicator (Misonix S-4000 liquid processor, Farmingdale, NY, USA), with a converter attachment (Q Sonica CL5, Newtown, CT, USA) (Parameters: 80 amplitude, 30s ON, 30s OFF, 2 cycles of 10 mins). Sonicated samples were centrifuged at 10,000 RPM, 4 ℃, for 30 minutes. The supernatant was filtered (0.22 μm) and loaded onto Sepharose CL-6B column (Cat. no. CL6B200, Sigma-Aldrich, St. Louis, MO, USA). According to our previously published protocol in *C. elegans*[39], 10 mL aliquot was collected using ice-chilled M9 Buffer and concentrated with an ultra-centrifugal filter (10 kDa MWCO, Cat No. UFC901008, Amicon, Merck KGaA, Darmstadt, Germany) and stored at-80℃ until further downstream processing. (Supplementary Figure S1c)

## Characterisation of small Extracellular vesicles (sEVs)

### Nanoparticle tracking analysis (NTA) and Dynamic light scattering (DLS)

NTA was employed to measure concentration and size of freshly diluted 1:10000 sEV samples in M9 Buffer. Approx 1000 particles/FOV were acquired for analysis by NanoSight NS300 (Malvern Panalytical, Netherlands), equipped with 488 nm laser and acquired by sCMOS camera. Brownian motion was tracked via light scattering, and the Stokes-Einstein equation was used to obtain particle sizes from mean^2^ displacement. Similarly, hydrodynamic diameter and average size were determined using nanoPartica SZ-100V2 (HORIBA, Ltd, Japan) (parameters: refractive index of 1.335 for M9 Buffer, scattering angle=173°, @ 25℃).

### Western blotting

Isolated sEVs were lysed in 10X RIPA buffer (Cat No. 20-188, Merck Millipore, Darmstadt, Germany) and mixed with protease inhibitor (Cat No. BMP1001, Abbkine Scientific, Atlanta, GA, USA). Protein was quantified using Bradford reagent Cat. No. B6916, Sigma-Aldrich, St. Louis, MO, USA) with bovine serum albumin as standard. 50 μg protein was separated on an 8% polyacrylamide gel, and wet transfer was done onto a nitrocellulose membrane (100 mA for 1 hour). Followed by blocking with 5% skim milk and incubation with antibodies against exosomal EV markers CD63 (Cat No. 10628D, Thermo Fisher Scientific, Waltham, MA, USA) and flotillin-2 (Cat No. 610383, BD Biosciences, San Jose, CA, USA) with a dilution 1:1000 and 1:5000, respectively, for 16 h at 4°C. The membrane was washed 3X for 15 min with TBST buffer and incubated with anti-goat HRP-conjugated (Cat No. A21010, Abbkine Scientific, Atlanta, GA, USA) at a dilution of 1:10000 and then visualised using ECL substrate (Cat No. BMU102, Abbkine Scientific, Atlanta, GA, USA).

### Field Emission Scanning Electron Microscope (FESEM) and Transmission Electron Microscopy (TEM)

A 10 μL diluted sEVs sample was air-dried on 16 mm circular cover slip. Followed by gold sputtering for 8s after mounting on the stub for FESEM analysis (Magnification 50.45 KX, WD 8.8 mm at 15 KV) (GeminiSEM 500, Carl Zeiss, Germany)

For TEM analysis, sEVs sample was fixed with 4% PFA; a 10 μL sEVs volume was drop-cast onto a holey carbon-supported copper grid (300 mesh, Ted Pella, Inc., USA) and air-dried for 1 min. Followed by negative staining with 2% uranyl acetate. Images were obtained using HR-TEM (Parameters: 200 kV HV and 0.04 s exposure, JEM-F200, JEOL, Tokyo, Japan) equipped with Gatan OneView camera.

### Flow cytometry

Flow cytometry was used to examine the homogeneity of sEV population. sEV samples were loaded into the BD FACSAria™ III at a flow rate of 4, equipped with 505 nm laser and 230-450 V (BD Biosciences, San Jose, CA, USA). BD FACSDiva software was used to discriminate and gate these EVs based on size to generate dot scatter plot (x-axis= forward scatter, y-axis= side scatter).

### Fourier transform infrared spectroscopy (FTIR) and Raman spectroscopy

For bio-fingerprint profiling, sEV samples were placed on ATR crystal of Nicolet iS50 FTIR Spectrometer (Thermo Fisher Scientific, Waltham, MA, USA). Spectra were collected to measure % transmittance over 4000–400 cm⁻¹ with 64 scans in triplicate. Similarly, for Raman spectroscopy, concentrated sEV samples were drop-casted on glass coverslips and air-dried for 1 hour under sterile conditions. Peak acquisition was done for three different samples (glass coverslip, coverslip with air-dried M9 buffer, and air-dried sEVs in M9 buffer on glass coverslip). All spectra were collected from 100 to 1200 Raman shift (cm^-1^) from the centre as well as the edge of the dried drop (Parameters: λ_ex_=532 nm, Zeiss Epiplan 50x objective, NA 0.65, grating 600 g/mm, and 10 accumulations for 20 s of integration time) using WITec alpha300 R (WITec GmbH, Ulm, Germany). Baseline correction was made before generating final spectra.

### Isolation, library preparation and sequencing of small RNA from sEV fractions

Total exosome RNA & protein isolation kit (Cat. no. 4478545, Invitrogen, Thermo Fisher Scientific, Waltham, MA, USA) was used for RNA isolation, and small RNA was enriched according to the manufacturer’s instructions. Elutions were made in 30 μL of nuclease-free water (NFW), and quality assessment was done using an Agilent 2100 Bioanalyzer (Agilent Technologies, USA), and concentrations were quantified using a Qubit 4 Fluorometer (Invitrogen, Thermo Fisher Scientific, Waltham, MA, USA).

For library preparation, the QIAseq miRNA library kit (Cat no. 331502, QIAGEN, Hilden, Germany) was used. In brief, pre-adenylated adapters were ligated to the 3’ ends of all miRNAs by miRNA 3’ ligase, while 5’ adapters were subsequently ligated to the 5’ end. These miRNAs were converted to cDNA using a reverse transcription primer integrated with a Unique Molecular Index (UMI), assigning a UMI to every miRNA, and cleanup was done using magnetic beads. Library amplification was done using an aliquot from a single-well miRNA 96 index kit plate, along with premixed oligos used for PCR (unique i5 index was premixed with i7 index) (Supplementary figure S1l). Following amplification, library cleanup was performed, and a quality check was done using Bioanalyzer® (Cat. no. 5067-4626, Agilent Technologies, Santa Clara, CA, USA). Next, sample-wise (triplicate) libraries were sequenced on the Illumina NextSeq 2000 System (Illumina, Inc., San Diego, CA, USA) using single-end 72 bp reads with 10 bp dual indexing and a minimum of 10 million reads were generated for each sample.

### RNA-Seq data analysis

For miRNA analysis, base call files were generated by the sequencer, converted to raw FASTQ files and demultiplexed using bcl2fastq Conversion Software (v2.20)[40]. Initial quality assessment of raw reads was performed using FastQC (v0.11.9)[41]. To process the miRNA fragments, raw reads were trimmed of adapter sequences using cutadapt (v4.1)[42], followed by UMI extraction and duplication removal before alignment with the reference database miRBase (v22, released March 2018)[43] using Bowtie 1(v1.3.0)[44]. Mapped reads were quantified using feature counts and normalised to counts per million (CPM). This was followed by log_2_ transformation with a pseudo-count of 1 (log_2_(CPM + 1). This method[18, 45] preserves relative expression changes suitable for miRNA differential analysis while accounting for variations in sequencing depth among the 12 sEV samples (A1-A3, control; B1-B3, damaged; C1-C3, response; D1-D3, recovery). Fold-change calculations were made by averaging the group-wise mean log_2_CPM values across replicates (Supplementary Table S2). Differential expression of miRNAs was also analysed by t-test (Parameters: *padj* < 0.05 and |log_2_fold change|> 1.5, equivalent to ∼ 2.8-fold change). Mean log_2_CPM values were used for principal component analysis (PCA) and heatmap generation using R software (v4.5.1)[46]. For additional information from the small RNA sequencing, we also used the analysis tool miRge3.0[17], developed for miRNAs and other small-RNA species.

### miRNA target prediction

The query sequences of differentially expressed miRNAs were retrieved from miRBase[47] (https://www.mirbase.org/) using miRNA_db_id (e.g. MIMATXXXXXXX). The miRNA sequences were saved in FASTA format separately for upregulated and downregulated miRNAs of the respective groups. Annotated *C. elegans* (PRJNA13758; WS290) 3′UTR sequences were downloaded in FASTA format from WormBase ParaSite[48] Version WBPS19 (WS291) using the BioMart tool. The downloaded FASTA file was filtered for 3′ UTR isoforms (24,090 isoforms), removing sequences with length < 30 nt, and used for downstream target analysis. miRanda[49, 50] algorithm was used for miRNA target predictions. For miRNA seed complementarity to its target, we used an energy threshold of-15.0 kcal/mol and a 155.0 score threshold. All the predictions produced by miRanda algorithm were generated using Cytoscape[51] software platform and subsequently visualised and explored using NetworkAnalyst[52].

### Protein Profiling of small Extracellular Vesicles

The eluted sEV samples were used for protein isolation. Briefly, RIPA lysis buffer (10X, Cat no. 20188, Sigma-Aldrich,St. Louis, MO, USA) and protease inhibitor cocktail (100X, Cat no. BMP1001, Abbkine, Wuhan, China) were added to create 1x reaction setup and kept for 18-20 hours incubation at 4℃. Total of 50 μg protein was used, and volume was raised to 200 μL by adding ammonium bicarbonate buffer (ABC). The buffer suspension was vortexed before adding 10% DTT and kept in a water bath for 1 hour at 60℃ for disulfide bond reduction. Followed by the addition of 10% IAA for alkylation of free thiols and kept for 30 min at room temperature (dark conditions). For enzymatic digestion of proteins, trypsin was added and incubated for 16 hours at 37℃. After incubation, formic acid was added to stop reaction. Column cleanup was done with solution A, which consisted of 50:50 (v/v) mixture of acetonitrile and water and then solution B, consisting of a 5:95 (v/v) mixture of acetonitrile and water. A 200 μL sample was added to the washed column, which was attached to a collection tube for binding (3 times). A 100 μL of solution B was added and centrifuged at 6000 RPM for 15 min for washing steps. At last, 100 μL of elutions were made with solution C consisting of a 70:30 (v/v) mixture of acetonitrile and water in collection tube at 6000 RPM. Eluted samples were evaporated to dryness using centrifugal vacuum concentrator (Cat No. 7810010, CentriVap® Benchtop Vacuum Concentrator, Labconco Corporation, MO, USA), leaving a solid substance at the bottom of the microcentrifuge tube and reconstituted by adding 100 μL of 0.1% formic acid. The collected samples were transferred to glass vials and analysed by LC–MS/MS using a C18 Easy-Spray column (Thermo Fisher Scientific; 50 cm, 3.0 μm) coupled to Q Exactive Plus Hybrid Quadrupole-Orbitrap mass spectrometer (Thermo Fisher Scientific, San Jose, CA, USA). Peptides were separated using a 60-min LC gradient and analysed by positive-mode nano-electrospray ionisation with full MS acquisition over *m/z* 200–2000.

### Mass spectrometry data processing

Raw Orbitrap data were processed with SEQUEST HT algorithm within Proteome Discoverer (v2.3.0.523, Thermo Fisher Scientific, San Jose, CA). The *C. elegans* UniProt reference proteome (Proteome ID UP000001940, download/revision 2025-06-12) was used for further analysis (parameters: trypsin digestion, carbamidomethylation of cysteine as a fixed modification, oxidation of methionine and protein N-terminal acetylation as variable modifications). Precursor and fragment mass tolerances were set according to the Q Exactive acquisition settings. Peptide-spectrum matches and proteins were filtered (target–decoy strategy to achieve a false discovery rate of 1%) at both peptide and protein level. Label-free quantification was performed in Proteome Discoverer based on precursor ion intensities across all runs, and protein abundances were exported to a matrix for downstream analysis.

### Proteomic data analysis

Proteome datasets were analysed using RStudio v2025.09.2 (Posit Software, PBC, USA)[53] and the Perseus platform v2.0.11 (Max Planck Institute of Biochemistry, Germany)[54]. Abundance values of triplicate set were loaded onto the Perseus software’s generic matrix. Protein abundance data were transformed into log_2_fold intensities, and missing values were also imputed. Sample groups were defined using categorical annotations for control, damaged, response and recovery. Associated P-value, P-adjusted value, and log_2_fold change were calculated using Welch’s t-test and Benjamini–Hochberg (FDR 0.05). Pearson correlation matrix and volcano plots were generated using RStudio v2025.09.2 (Posit Software, PBC, USA)[53].DE of all proteins were analysed by filtering proteins with [*Padj* <0.05 and |log_2_FC| >1.5] after t-test matrix generation. DEPs with respect to the control were identified and selected for downstream analysis. Data visualisations, including heatmaps generated via the ComplexHeatmap package[55] and box and scatter plots generated via the ggplot2 package[56], were produced within RStudio v2025.09.2[53] (Posit Software, PBC, USA). VENNY 2.1[57] was used for analysing upregulated as well as downregulated protein sets separately among the damaged, response and Recovery phases. g: Profiler[58] was used to perform GO pathway enrichment analyses using g: GOSt functional profiling. Upregulated and downregulated proteins were used as ordered queries with respect to decreasing log_2_FC values of the proteins. P-values were adjusted to correct for multiple testing using the Benjamini–Hochberg method with FDR 0.05, and other parameters were set to default for identifying significantly enriched protein sets. The top 10 biological pathways (GO: BP) were used to compare among different groups. Downstream dynamics, including clustering and dimensionality reduction matrices generated via the Seurat package[59] and lineage trajectory and pseudotime calculations generated via the Monocle 2 package[60] within RStudio v2025.09.2 (Posit Software, PBC, USA; Posit Team, 2025)[53].

### Integrated miRNA-Proteome Sankey analysis

DE miRNAs (*Padj* < 0.05, |log₂FC| > 1.5) and proteins (*Padj* < 0.05, |log₂FC| > 1.5) from sEV samples of damaged, response and recovery worms were integrated through target prediction and pathway overlap analysis. High-confidence miRNA targets were identified by scanning *C. elegans* 3′UTR sequences (WormBase WS290, PRJNA13758) against DE miRNA seed sequences using miRanda[49, 50] (score > 140, E <-7 kcal/mol). The regulatory flow network was represented as a Sankey flow diagram generated via the ggalluvial package[61] within RStudio v2025.09.2[53] (Posit Software, PBC, USA). The method involves using three sequential layers of regulation: Phase-specific upregulated miRNAs → predicted target genes; downregulated proteins → parent genes; and shared biological pathways (KEGG/GO, etc.). This multi-omics antagonism approach (up-miRNA↑ vs down-protein↓, *viz. a viz*) uniquely captures post-transcriptional suppression of healing regulators, revealing temporal regulatory cascades absent from single-omics analysis.

### Pseudo-temporal protein Dynamics

We performed sample-wise pseudotime trajectory analysis of 582 proteins (adjusted p-value <0.05). A protein expression heatmap was generated and clustered into 5 hierarchical clusters (Cluster A to E) (Figure 8). A row-wise Z-score normalisation was applied to 582 proteins, where red indicates upregulation (positive Z-score) above the mean expression and blue indicates downregulation (negative Z-score) below average expression (Figure 8a). The grey line represents the individual protein trajectory, whereas the red line represents the mean trajectory of the total proteins in each cluster (Figure 8b-f).

### Cross-Species Sequence Similarity and Human Orthology Analysis

We identified these Cross-Species sequence similarities based on either ≥7/10 nt continuous 5′-end similarity or ≥70% full-length sequence similarity, excluding weaker 60–69.9% matches[32]. In g: Profiler[58] DEPs were used to perform orthology analysis for in humans using g: GOSt functional profiling.

### Stem-loop qRT PCR

This technique was used for orthogonal validation of the DE miRNAs. A total of 10 miRNAs (Figure 9A) were selected using filter of *Padj* < 0.05 and an absolute log2 value > 1.5 (∼ 2.8-fold change). cel-miR-64-5p and cel-miR-65-5p were selected as internal controls because they were expressed nearly uniformly across all replicates of different sample groups. The stem-loop primers specific to miRNAs were designed, adhering to established protocols[62] and procured (Supplementary Table S1). TaqMan™ MicroRNA Reverse Transcription Kit (Cat no. 4366596, Applied Biosystems, USA) was employed for the reverse transcription of all 10 selected miRNAs across all samples as per the manufacturer’s instructions. In short, 10 ng of isolated enriched RNA was used along with 7 µl of RT reaction mix and 3 µl of 5X stem-loop RT primer (5 µM) in a 15 µl reaction setup. Followed by cDNA synthesis using Veriti™ Thermal Cycler (Cat no. 4375305, Applied Biosystems, USA) (Temp cycles:30 min. at 16℃ (primer annealing), 30 min. at 42℃ (extension), and 5 min at 85℃ (enzyme inactivation)). cDNA was diluted 1:5 (v/v) to a final concentration of 10 ng/µL and used for SYBR green-based real-time PCR for each miRNA with a miRNA-specific forward primer and a universal reverse primer (Supplementary Table S1). Real-time PCR was performed using CFX Opus 96 Real-Time PCR System (Cat No. 12011319, BIO-RAD, USA) according to manufacturer’s instructions (Supplementary Table S3, S4).

Stem-loop primers were refolded[62] using Veriti™ Thermal Cycler (Cat no. 4375305, Applied Biosystems, USA). Precisely, 100 µL of oligonucleotides were mixed with molecular biology grade mineral oil (Cat no. M5904, Sigma-Aldrich, USA) to prevent evaporation. Placed in a sequential temperature program at 95℃ for 10 min,75℃, 68℃, 65℃, and 62℃ for 1 hour and then at 60℃ for several additional hours. At last, the refolded primer was transferred under oil to a new tube and stored at-20℃.

## Statistical analysis

All statistical analyses and data visualizations were performed using GraphPad Prism v10.1.2 (GraphPad Software, Inc., San Diego, CA), OriginPro 2024 (OriginLab Corporation, Northampton, MA, USA), and RStudio v2025.09.2 (Posit Software, PBC, USA; Posit Team, 2025) with appropriate packages. Multiple testing corrections were performed using the Benjamini–Hochberg method to control the false discovery rate (FDR < 0.05). For comparisons between two groups, an unpaired two-tailed Welch’s t-test was used, whereas one-way ANOVA followed by Dunnett’s or Tukey’s multiple-comparison test was considered for comparisons involving multiple groups. A p-value of < 0.05 was considered statistically significant.

## Data Availability

The sEVs-derived small RNA-seq data generated in this study have been deposited in the SRA database (BioProject: PRJNA1392877). For control, damage, response and recovery triplicate samples were deposited under the accession codes SRR36579191-SRR46579202. The mass spectrometry proteomics data have been deposited in the ProteomeXchange Consortium via the PRIDE17 partner repository with the dataset identifier PXD072254. Stem-loop qRT-PCR primers are provided in the Supplementary file.

## Acknowledgements

We thank Dr Meenakshi Sharma, ACBR, University of Delhi (DU), for providing N2 strain *C. elegans*; Dr Shourya Dutta Gupta, MSME, IIT Hyderabad, for Raman Spectroscopy access; IIT Delhi for NTA measurement; USIC, DU for DLS, TEM, FTIR and FESEM for sEVs characterisation. We thank Prof. Shibnath Mazumder, department of Zoology, University of Delhi (DU) for critically reading the manuscript draft and for their helpful suggestions. We thank ANRF for funding of the project, project number EEQ/2022/001002 and CSIR for the fellowship.

## Code Availability

No custom codes were developed in our study.

---

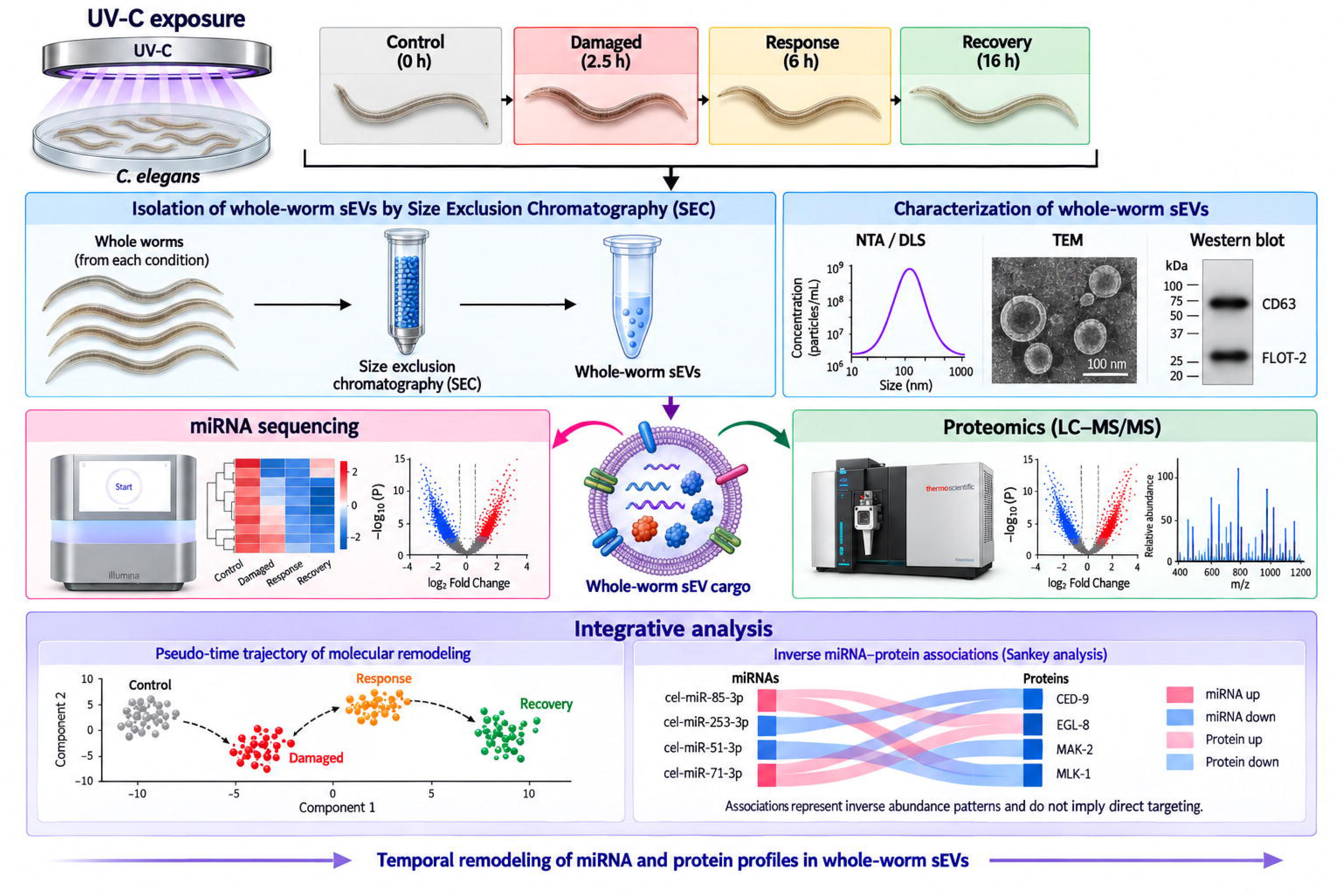

