## Supplementary file 1 for "Integrated miRNA-protein profiling of small extracellular vesicle cargo reveals dynamic cargo remodelling during tissue repair in UV-C-damaged *C. elegans*"

**This Supplementary file includes figures from S1a to S1l and Tables S1 to S5**

**S1a: Positive Control of dual AO and EtBr staining S1b: Positive Control of ROS activity**

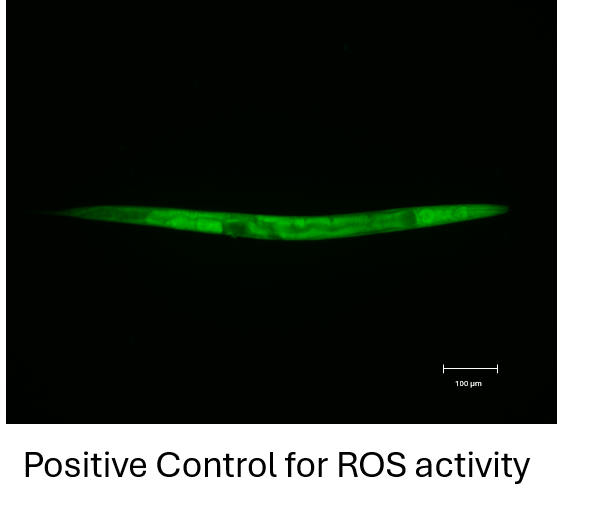

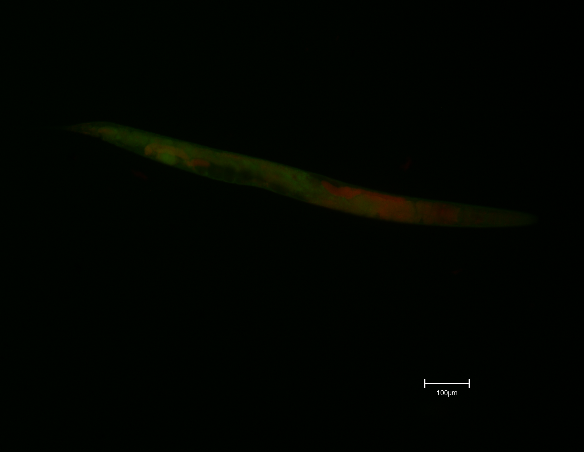

**S1c: CL-6B-based size-exclusion chromatography-Ultrafiltration method for sEV isolation method**

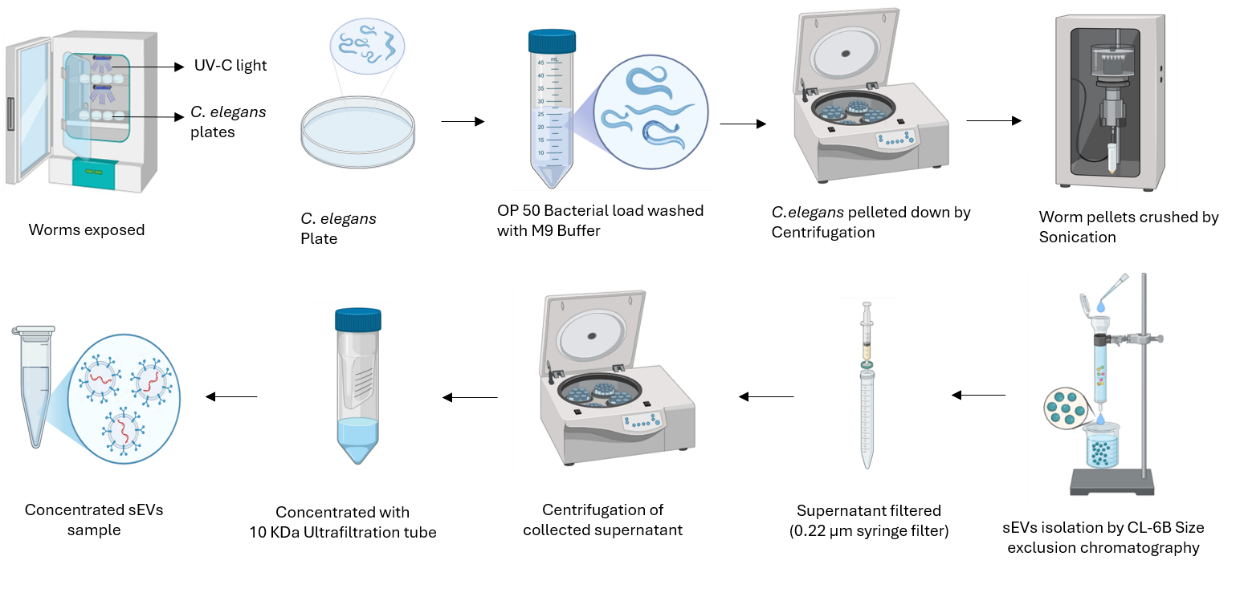

**S1d: Original Western Blot, uncropped images of immunoblotting for anti-CD63 and anti-flot2.**

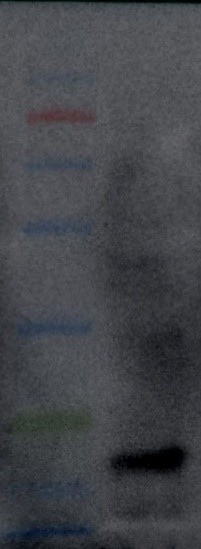

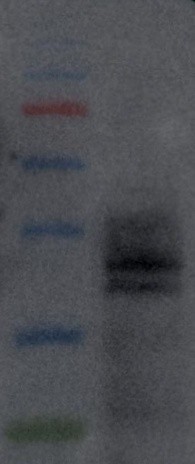

**Flotillin-2 Western blot (uncropped)	CD63 Western blot (uncropped)**

**S1e: Raw files for NTA analysis**

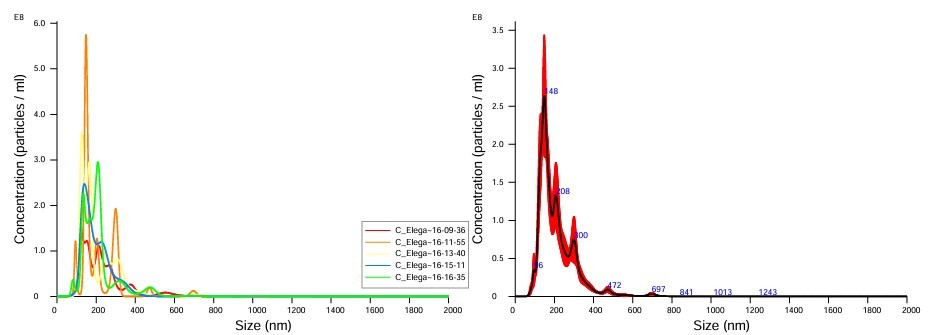

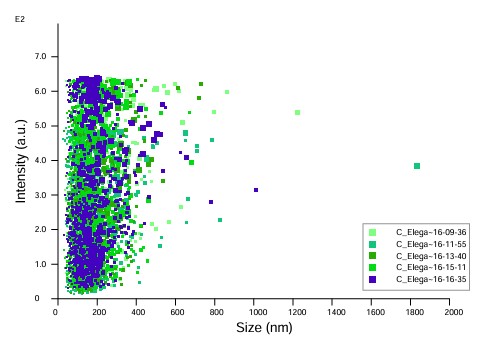

**S1f: Raman Spectra for sEVs characterisation**

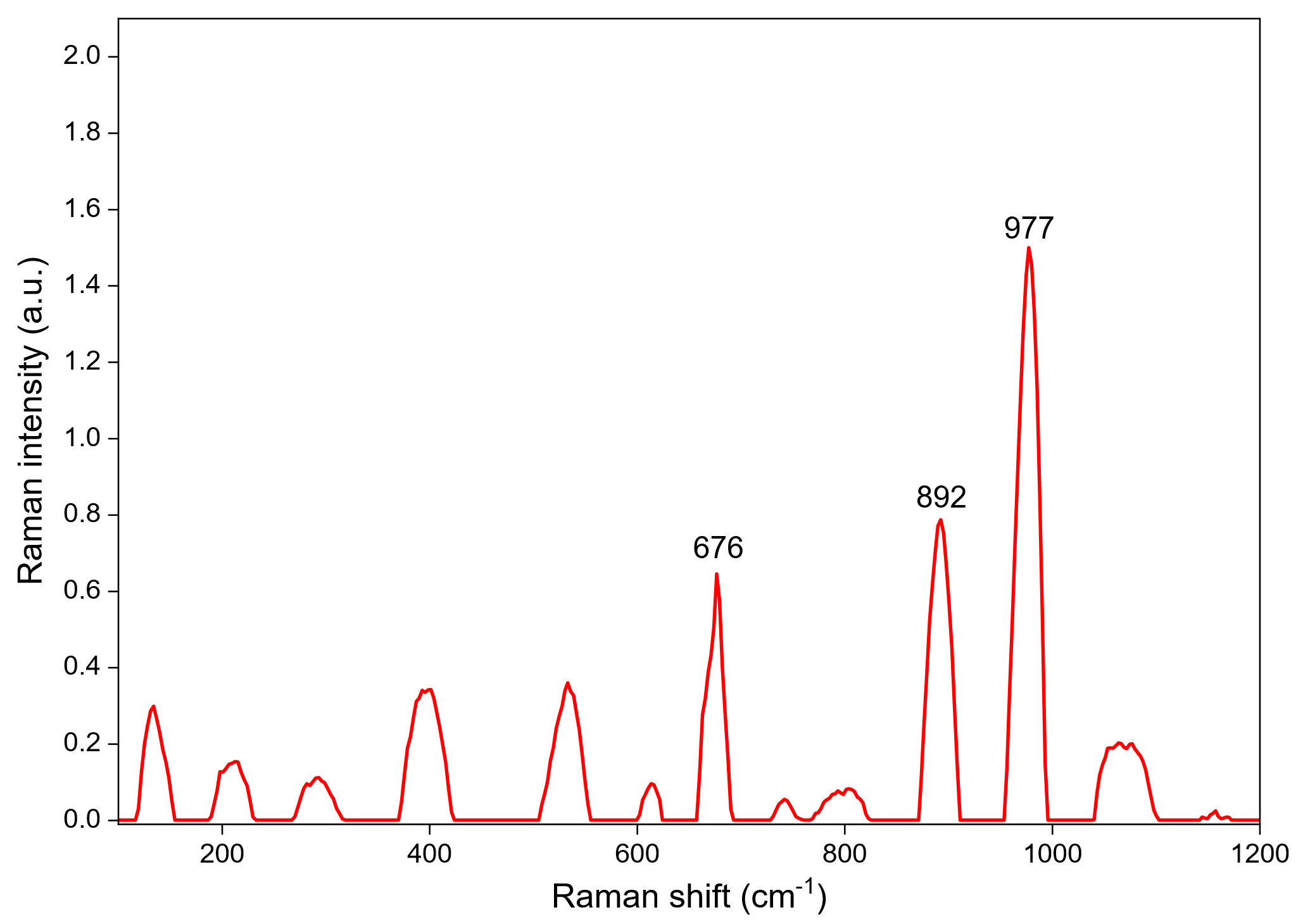

**676= Nucleic acid**

**892=lipids and proteins**

**977=Proteins**

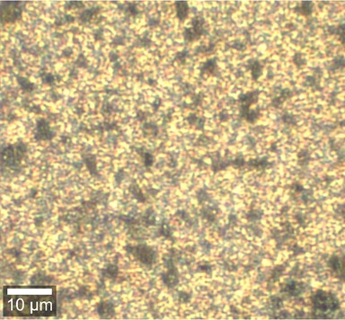

Optical image

Raman spectra showing vibrational signatures of molecular constituents of sEVs. The x-axis represents Raman shift (cm⁻¹; vibrational frequency), and the y-axis represents Raman intensity (a.u.).

**S1g: PCA plot for miRNA raw reads**

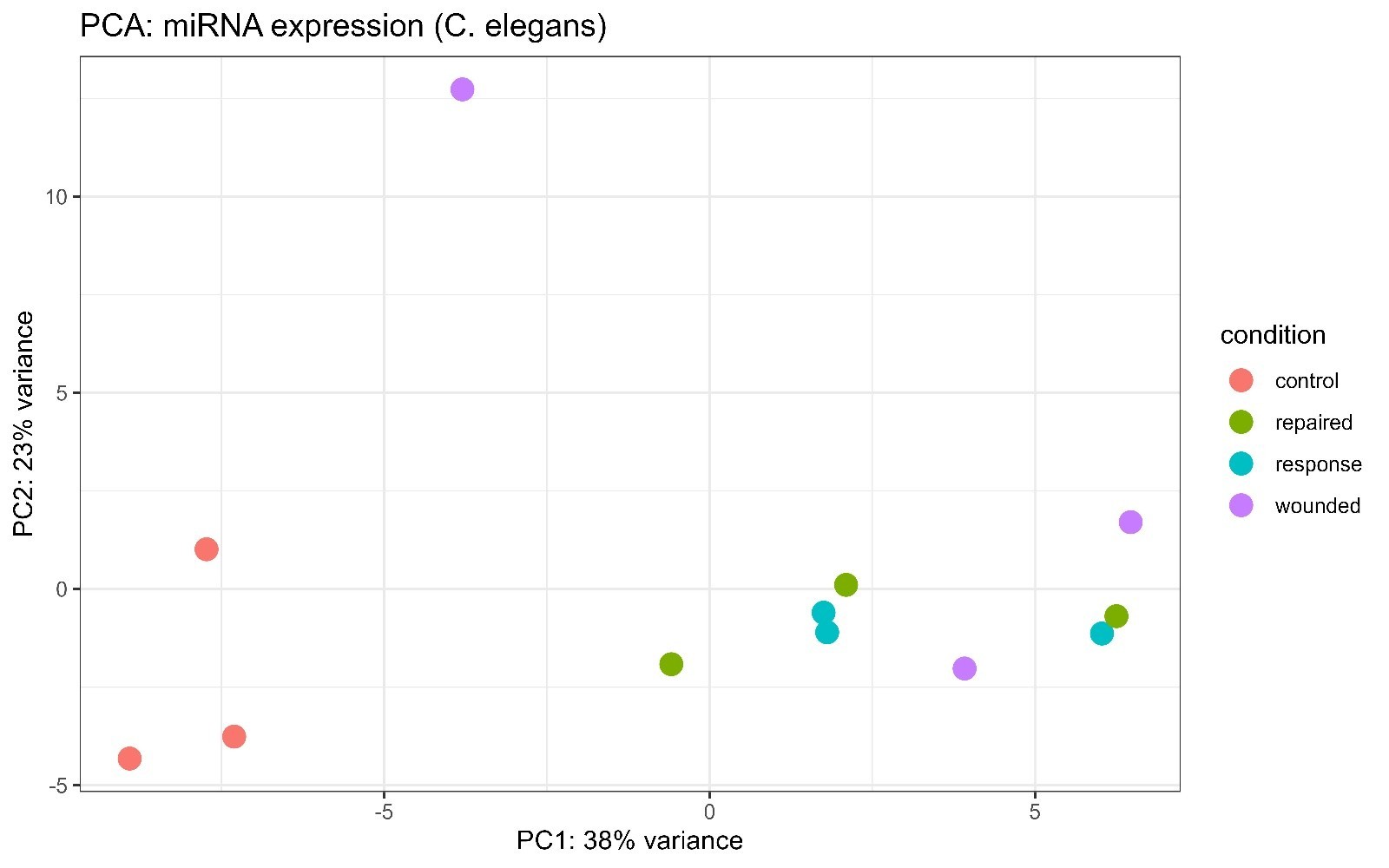

Control

Recovery

Response

Damaged

**S1h: Target network analysis of miRNAs and their respective protein target**.

**Damaged_down**

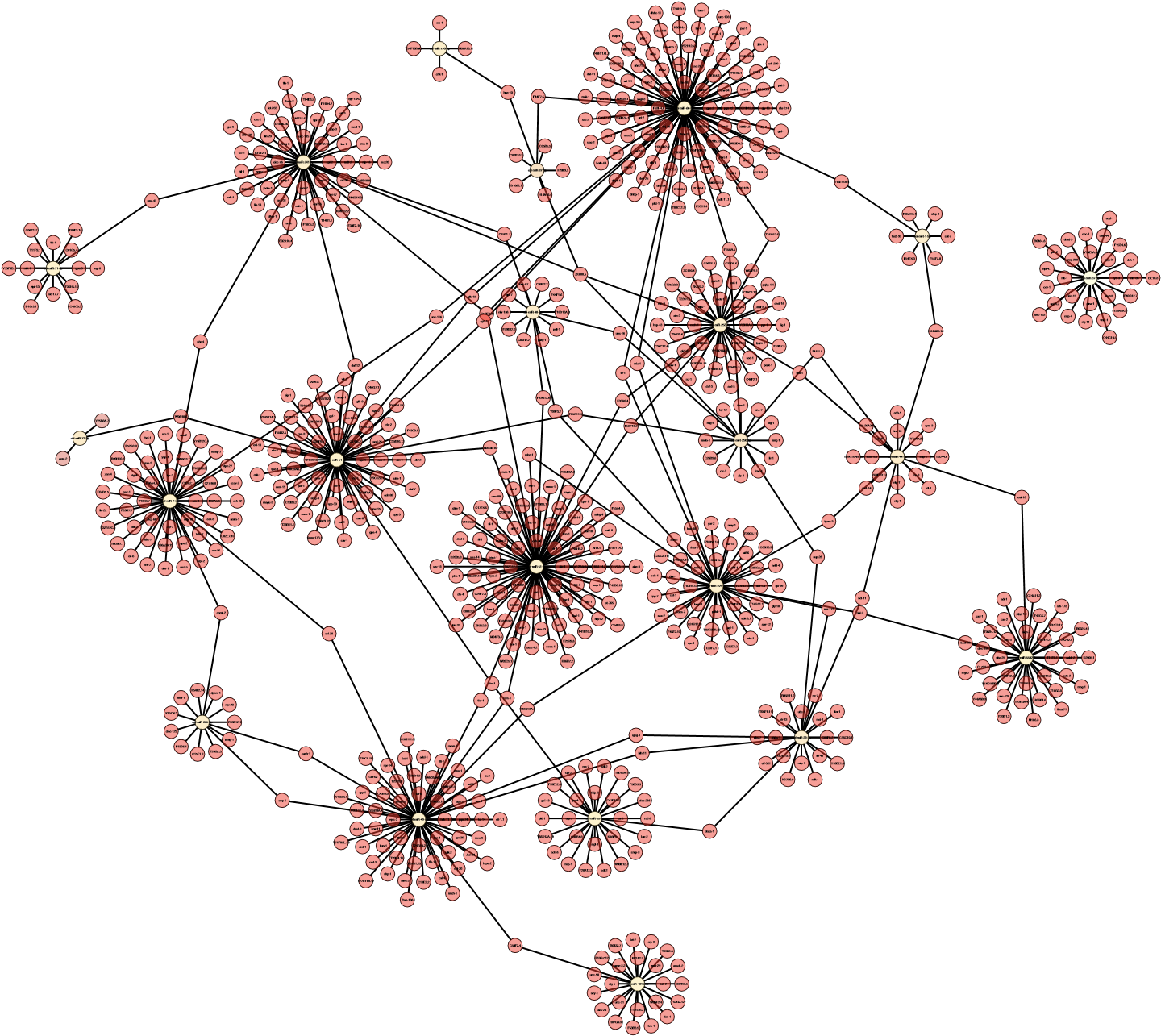

### **Damaged_up**

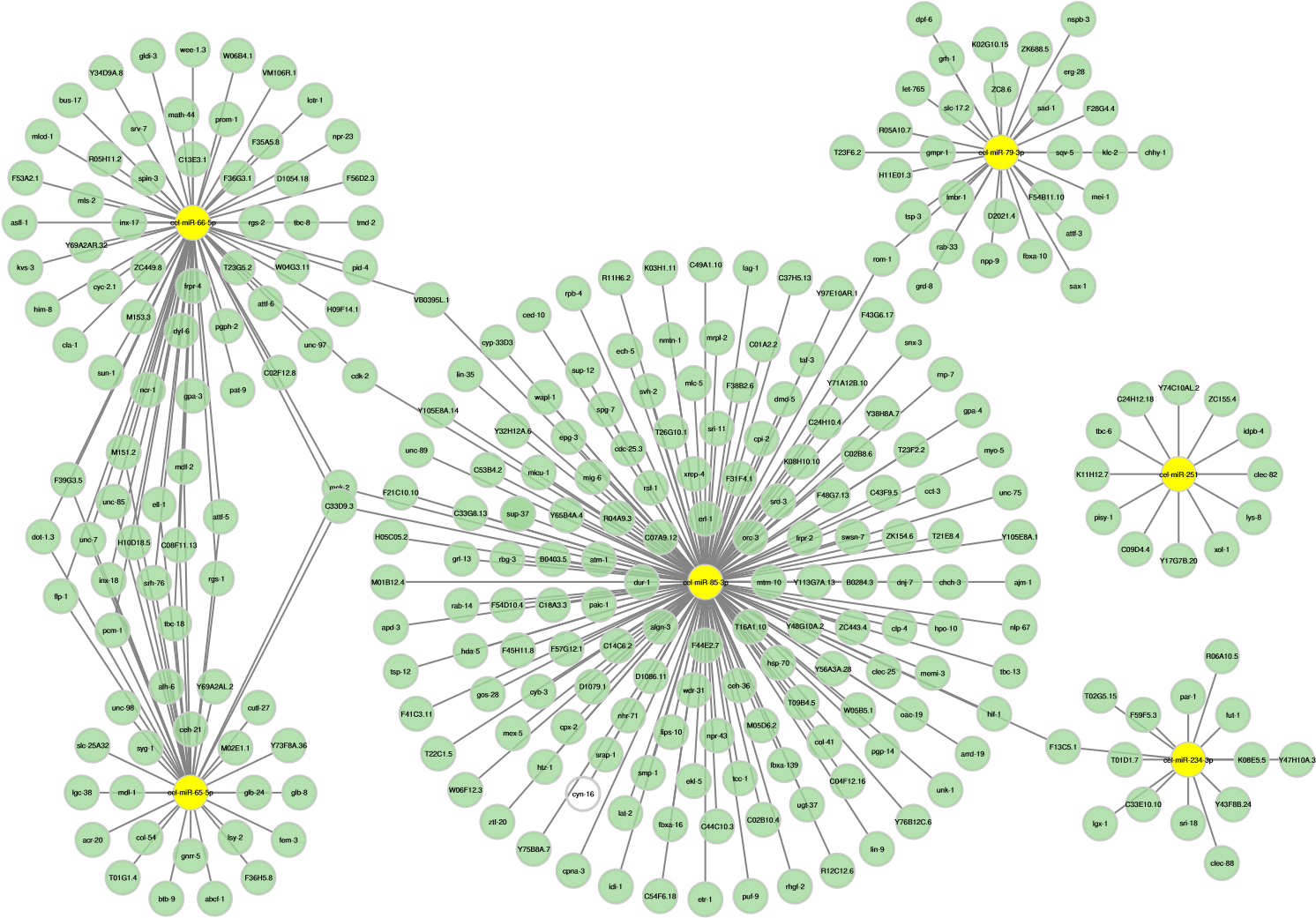

**Response_up**

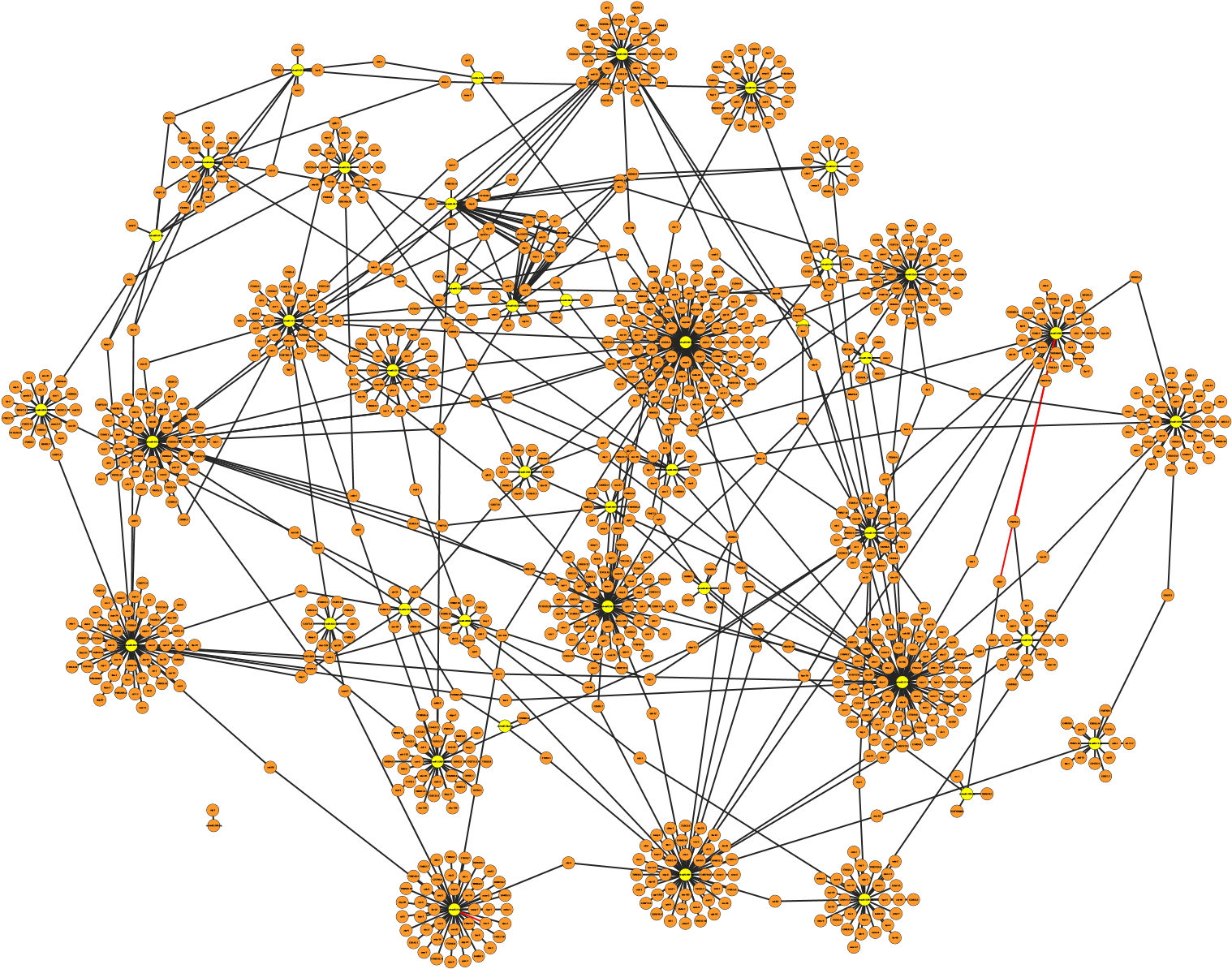

**Recovery_up**

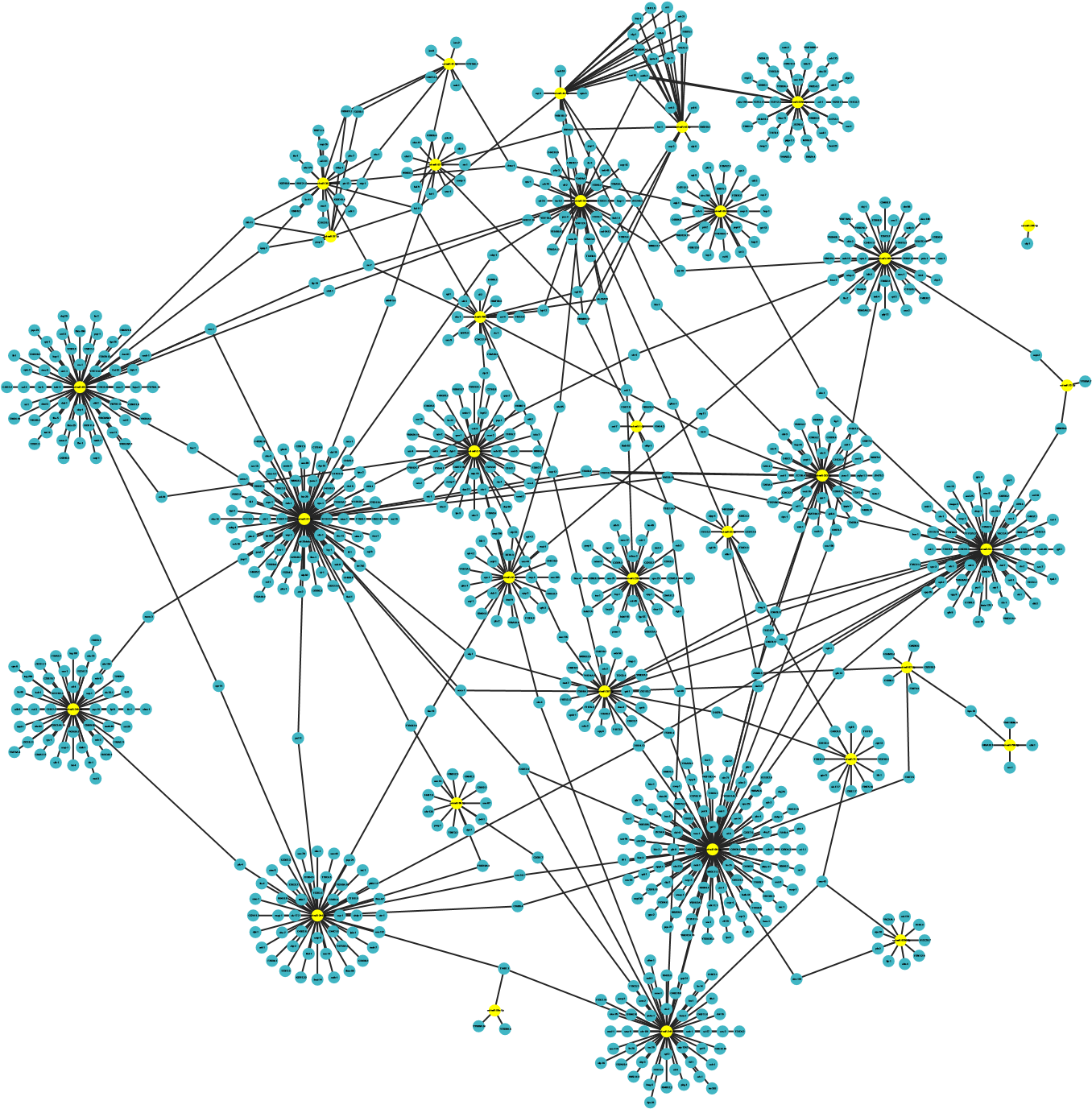

**S1i: STRING network analysis of DE miRNAs and their respective protein target**

**Damaged_up**

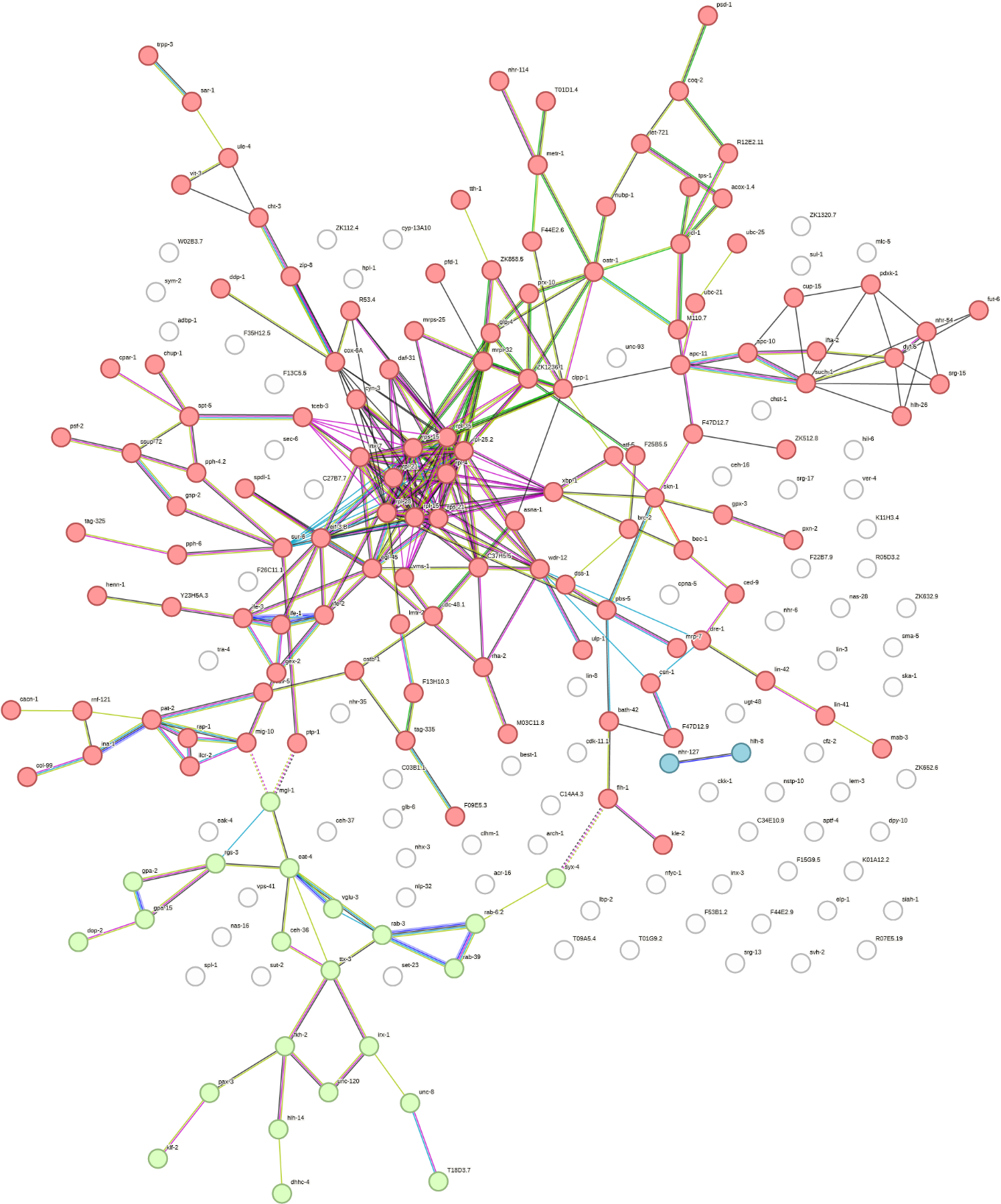

### **Damaged_down**

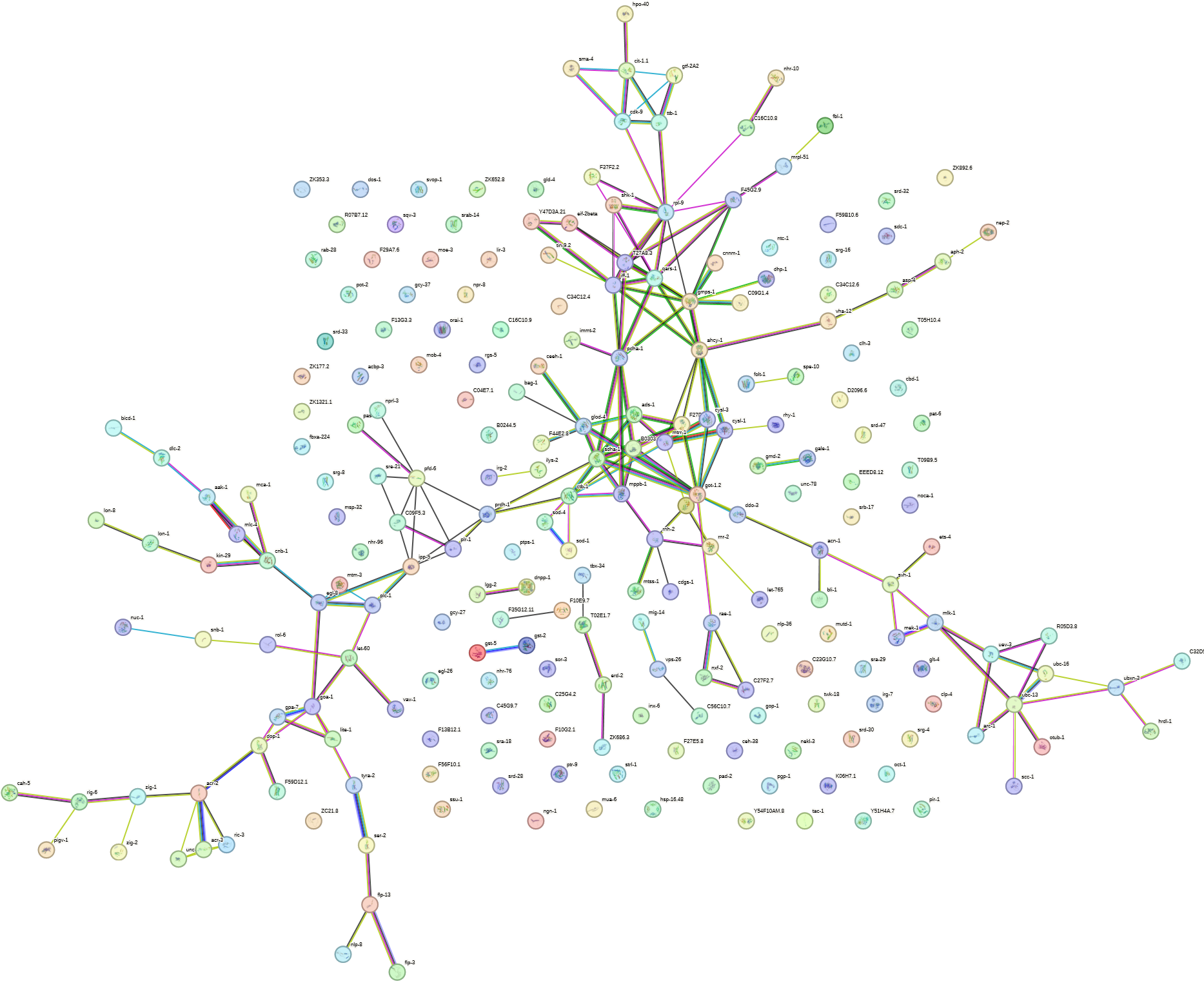

### **Response_up**

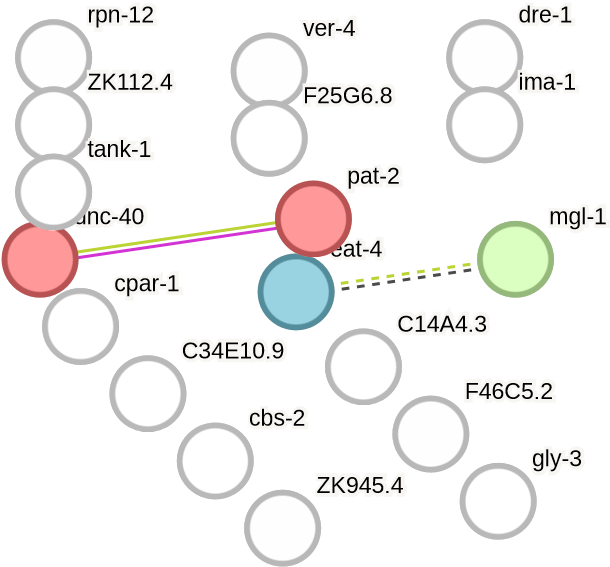

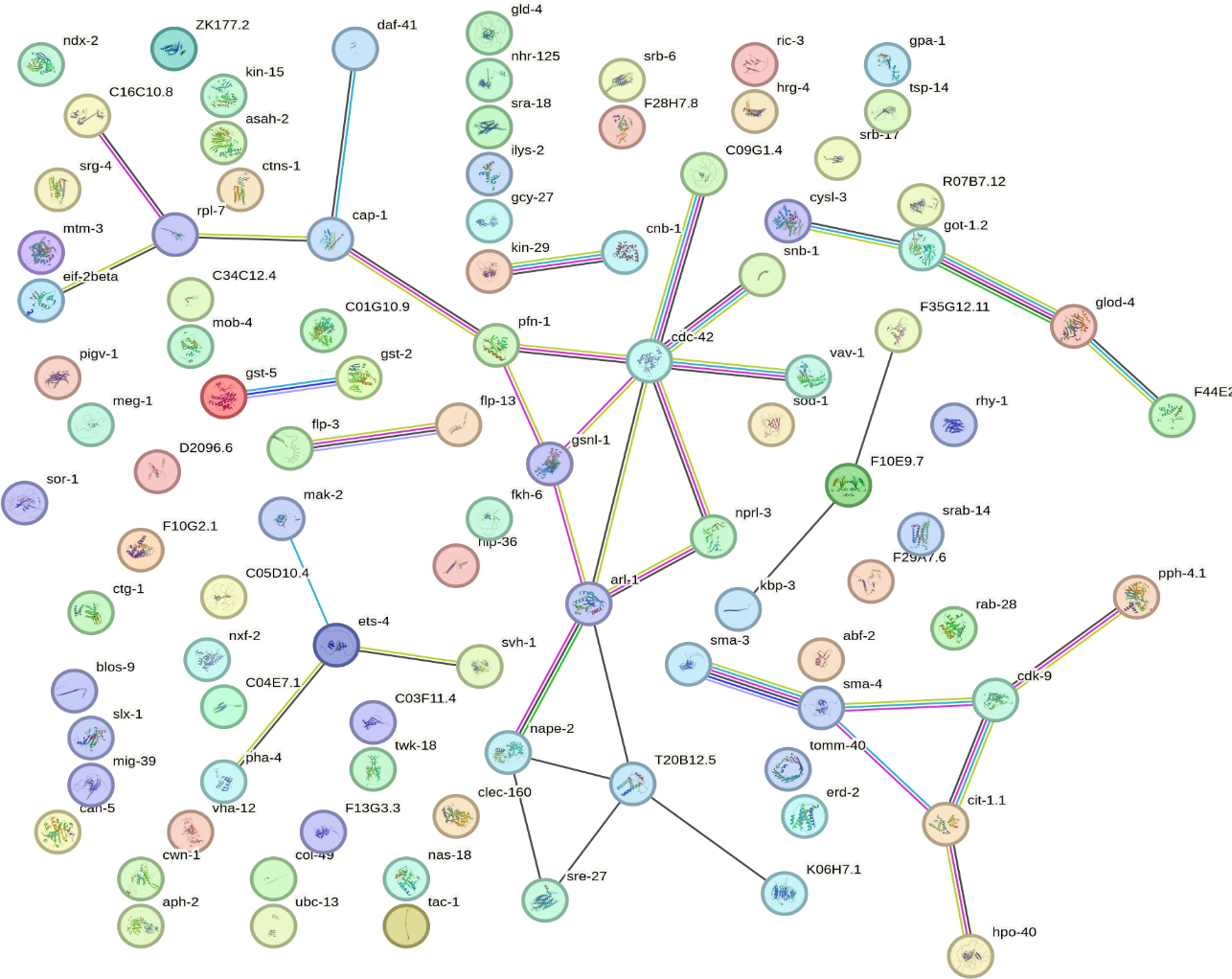

### **Response_down**

### **Recovery_up**

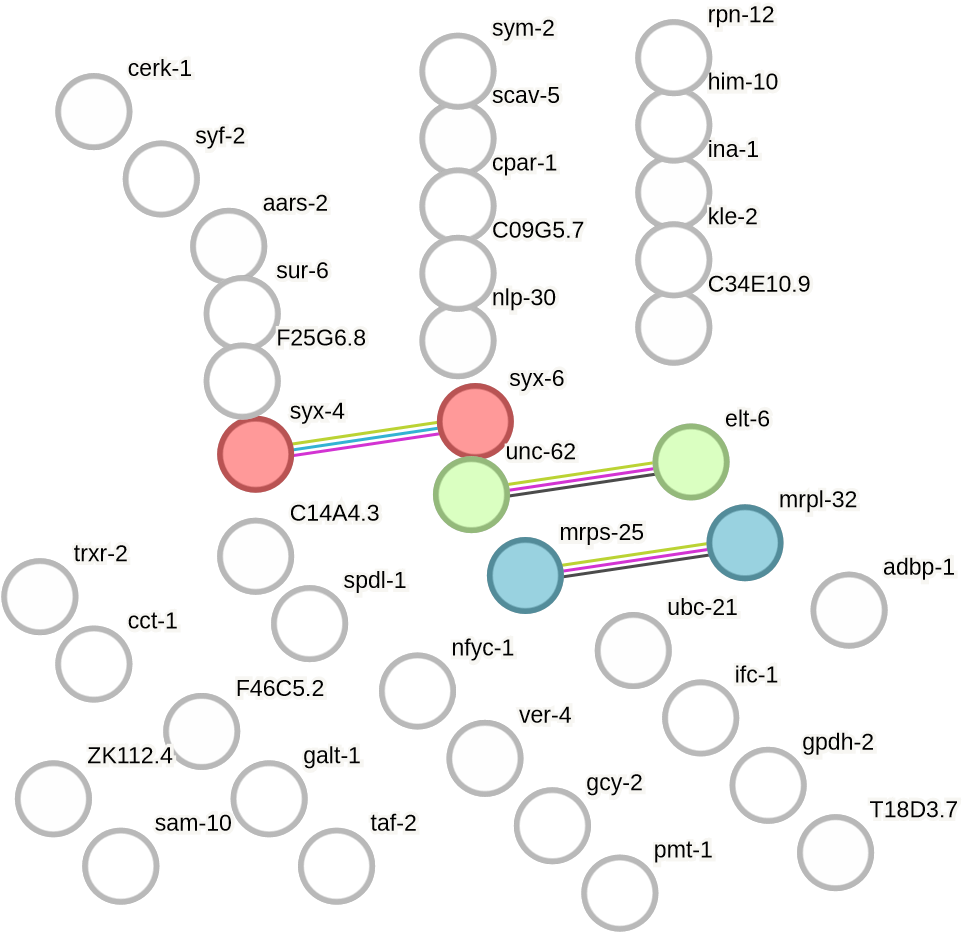

### **Recovery_down**

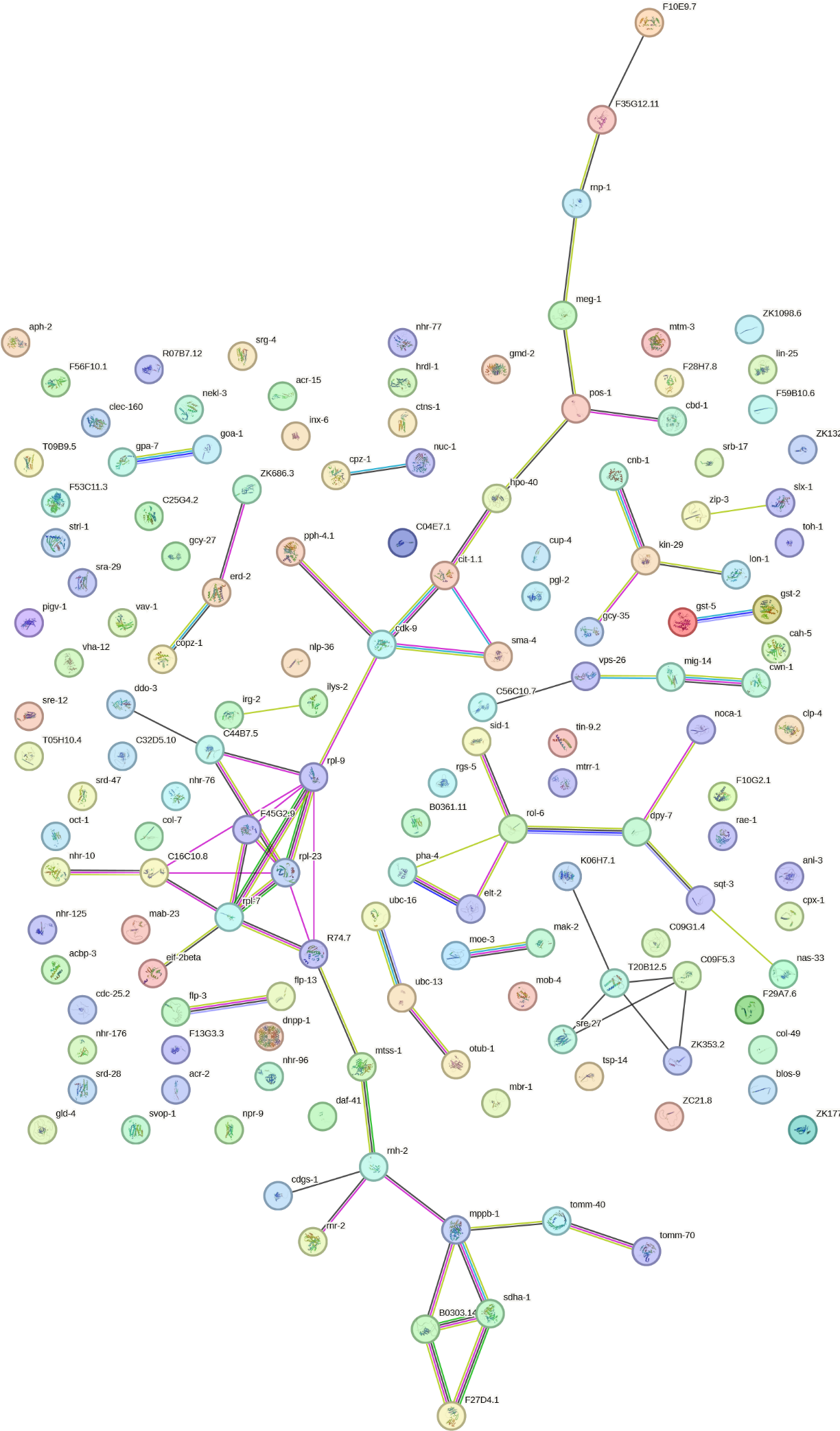

**S1j: Shanky diagram represents relationship between miRNA, target proteins and its associated biological processes**

### **Damaged_up**

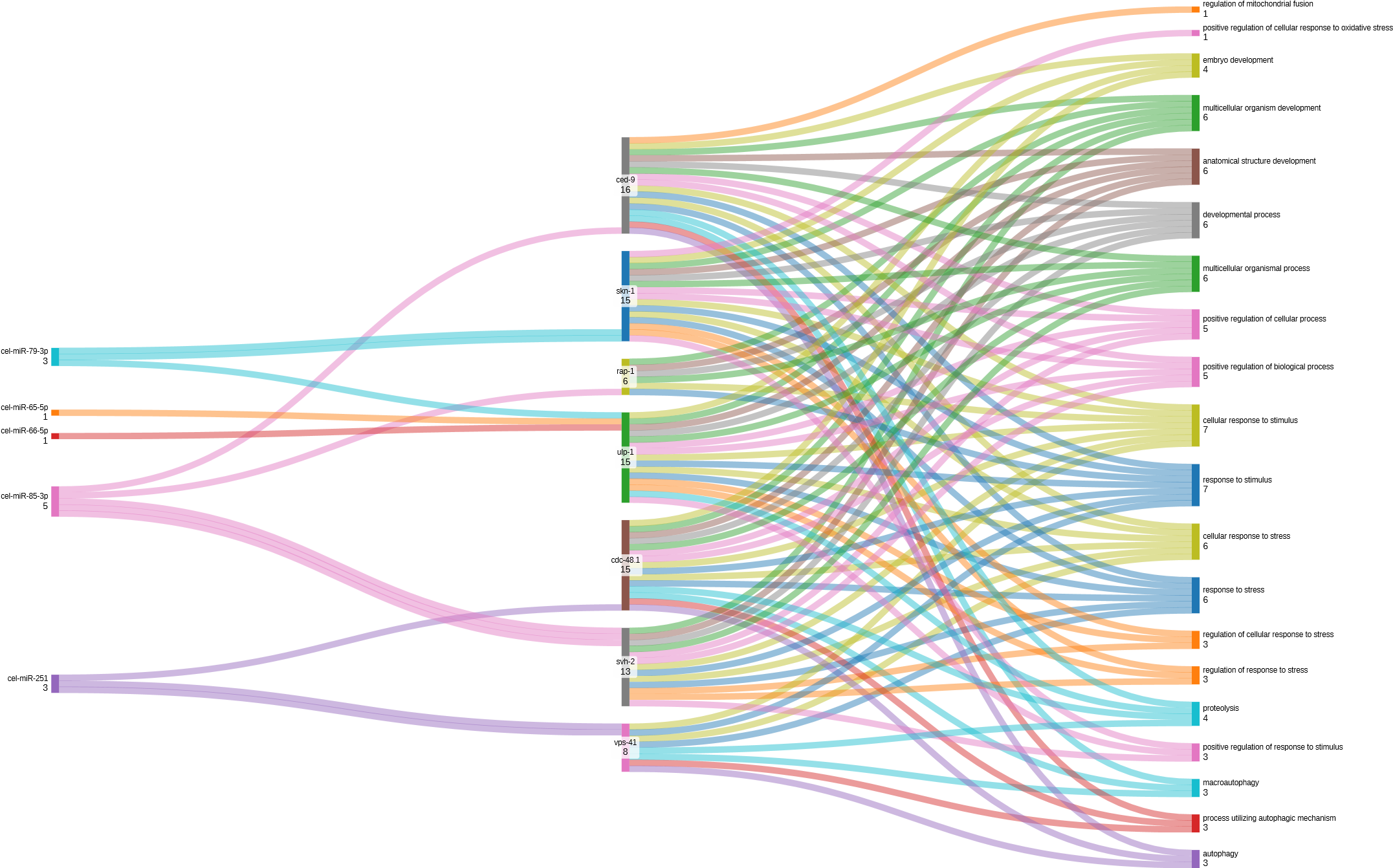

### **Damaged_down**

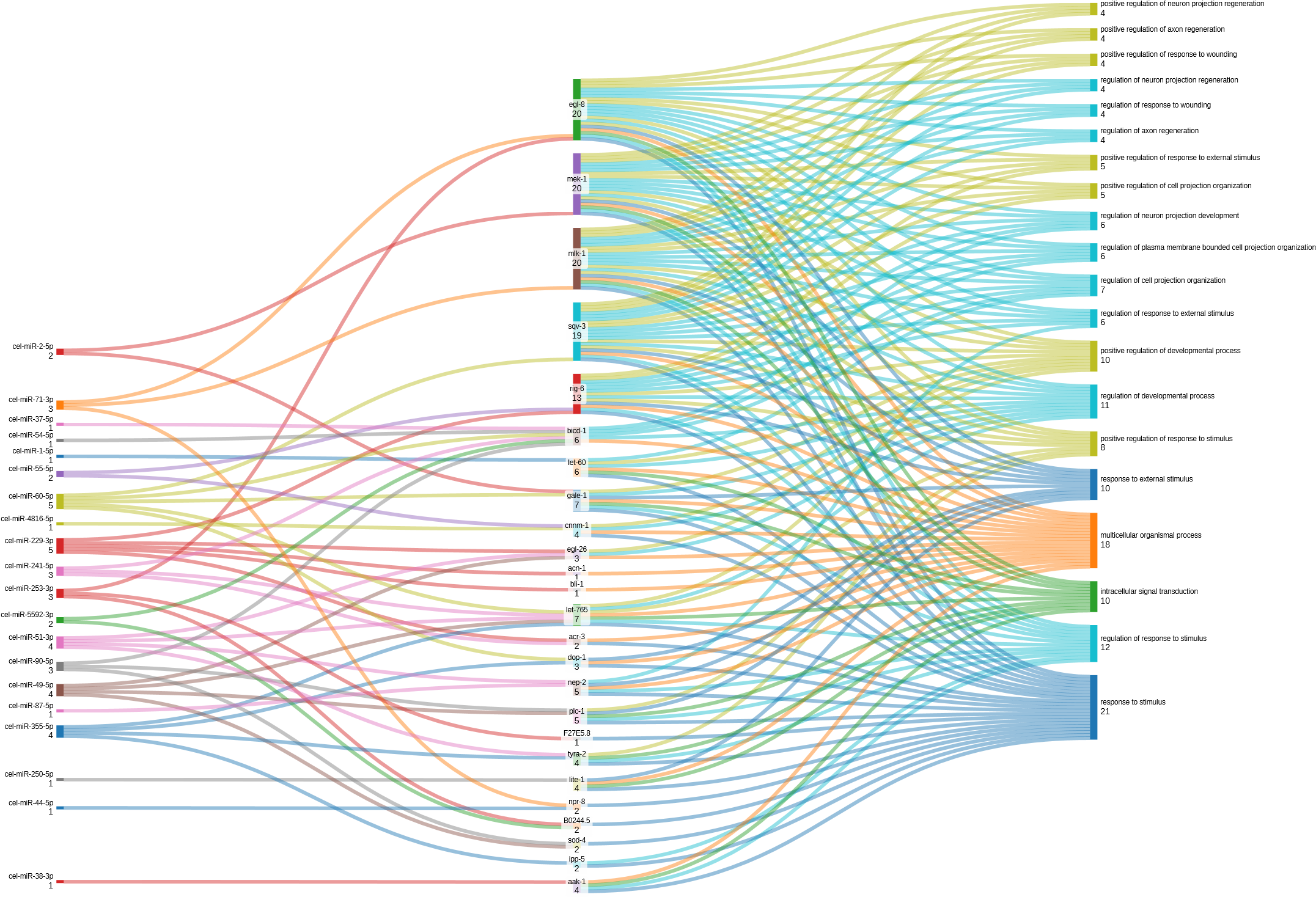

### **Response_down**

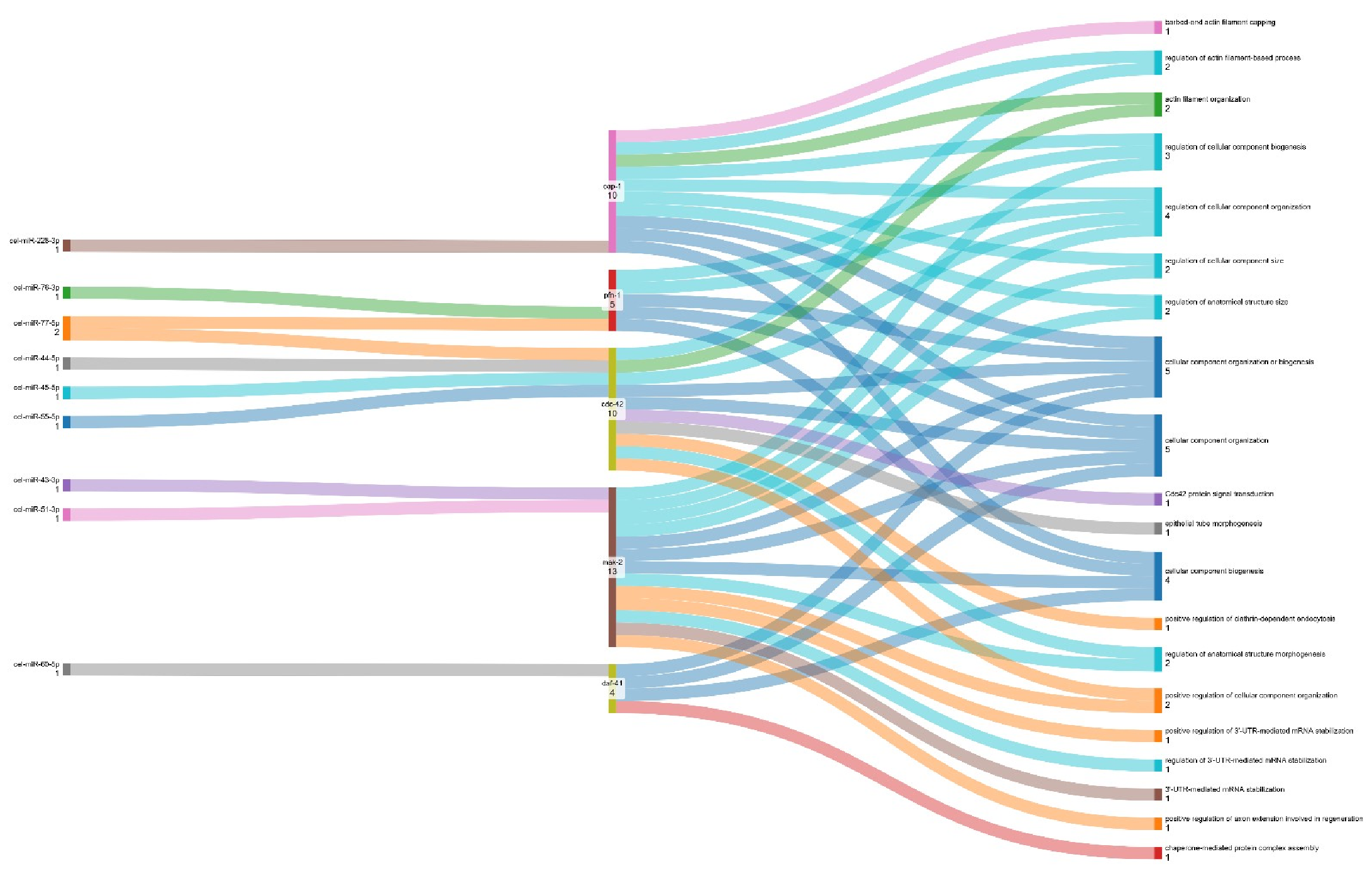

### **Recovery_down**

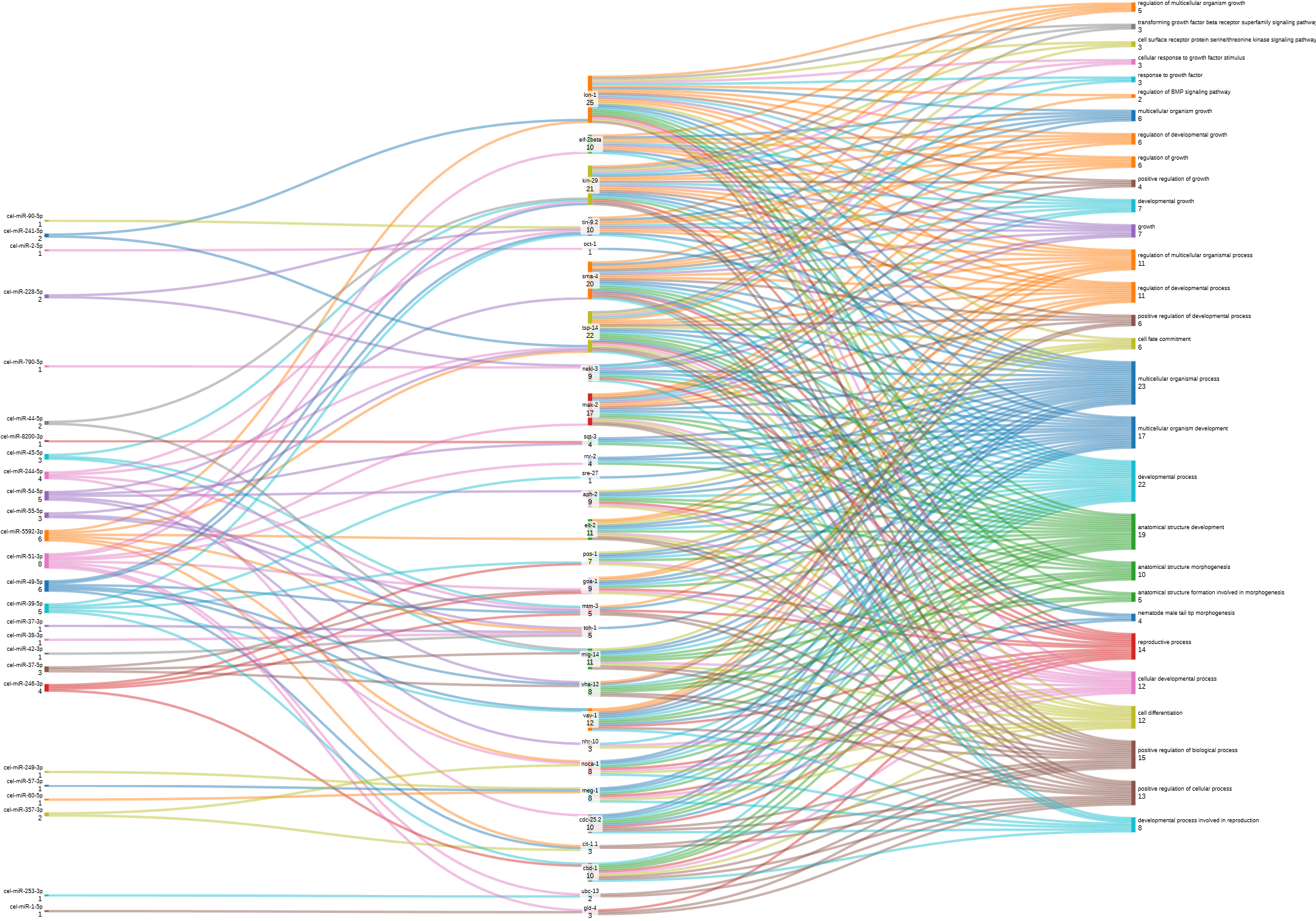

**S1k: Representative image of Melt curve output from Stem-loop RT-qPCR**

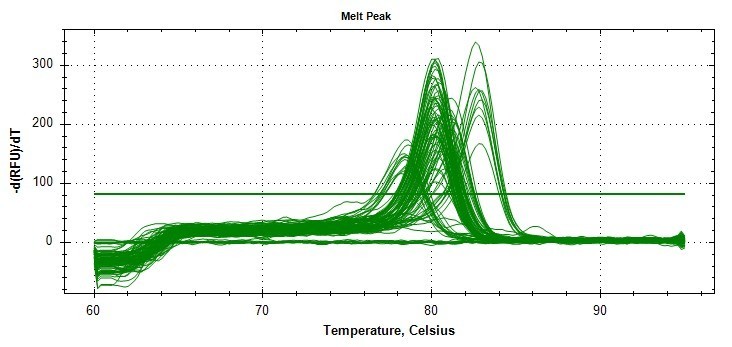

#### **S1l: Electrophoresis File Run Summary {High Sensitivity DNA Assay}**

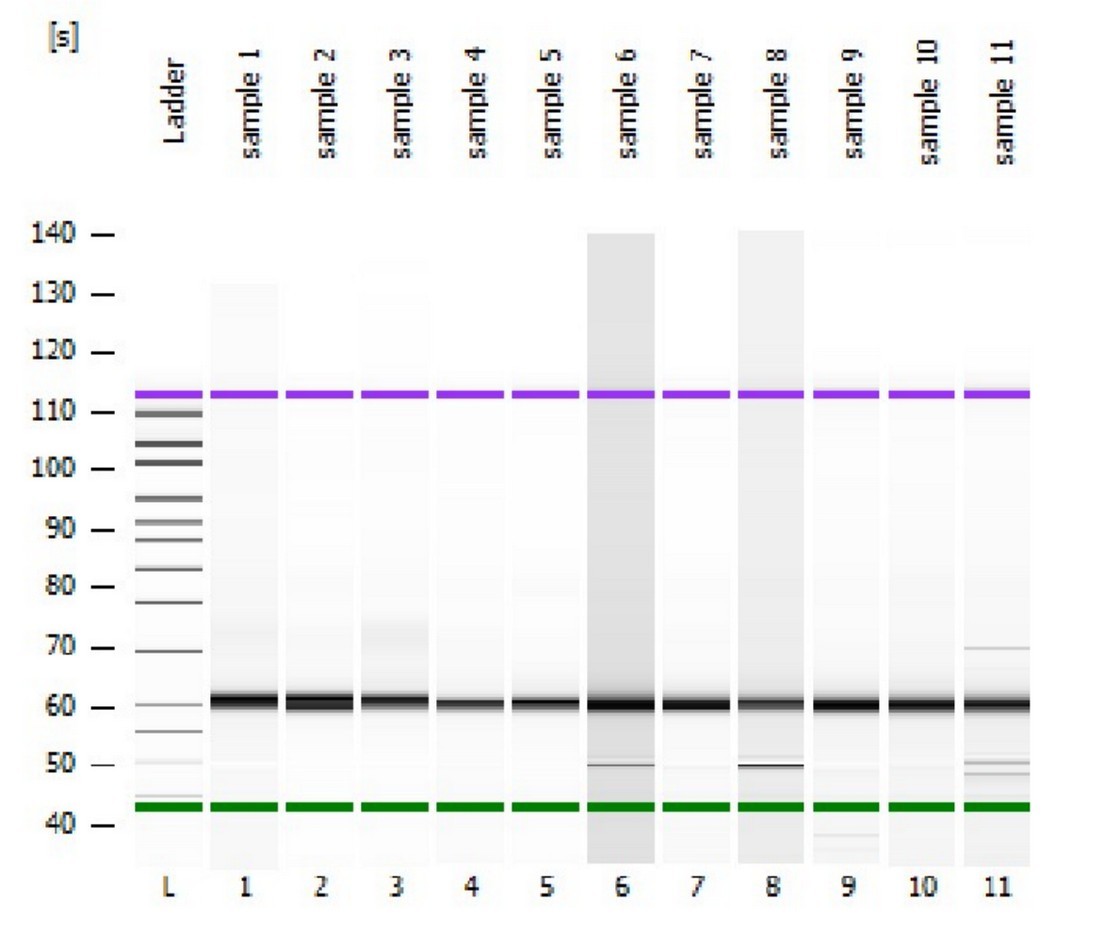

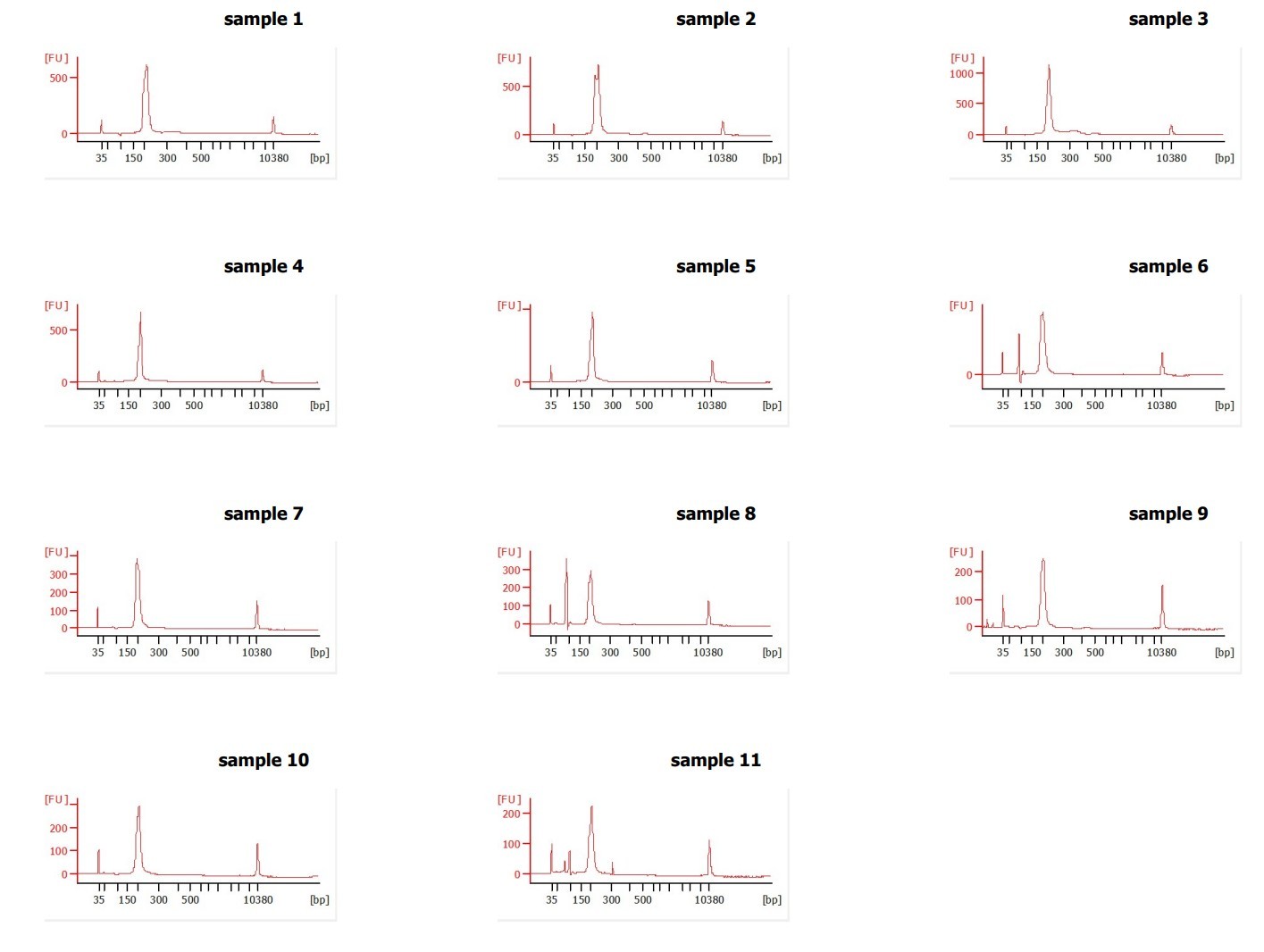

**Table S1. Stem-loop qRT-PCR Primers:**

| **miRNAs** | | **Forward Primer (5'-->3')** | **Universal Reverse Primer for all 10 miRNAs** |
| --- | --- | --- | --- |
| Cel-miR-253-3p | | TTAGTAGGCGTTGTGGGAAGGG | CCAGTGCAGGGTCCGAGGTA |
| Cel-miR-85-3p | | TACAAAGTATTTGAAAAGTCGTGC |  |
| Cel-miR-79-3p | | ATAAAGCTAGGTTACCAAAGCA |  |
| Cel-miR-90-5p | | CGGCTTTCAACGACGATATCAAC |  |
| Cel-miR-794-5p | | TGAGGTAATCATCGTTGTCACT |  |
| Cel-miR-249-3p | | TCACAGGACTTTTGAGCGTTGCC |  |
| Cel-miR-2-5p | | TATCAAAGCGGTGGTTGATGTG |  |
| Cel-miR-51-3p | | CATGGAAGCAGGTACAGGTGCA |  |
| Cel-miR-60-5p | | AACTGGAAGAGTGCCATAAAATC |  |
| Cel-miR-250-5p | | CCTTCAGTTGCCTCGTGATCCG |  |
| Internal control  Cel-miR-64-5p | | TATGACACTGAAGCGTTACCGAA |  |
| **miRNAs** | **Stem-loop RT Primer Sequences (5'---->3')** | | |
| Cel-miR-253-3p | GTCGTATCCAGTGCAGGGTCCGAGGTATTCGCACTGGATACGACCCCTTC | | |
| Cel-miR-85-3p | GTCGTATCCAGTGCAGGGTCCGAGGTATTCGCACTGGATACGACGCACGA | | |
| Cel-miR-79-3p | GTCGTATCCAGTGCAGGGTCCGAGGTATTCGCACTGGATACGACAGCTTT | | |
| Cel-miR-90-5p | GTCGTATCCAGTGCAGGGTCCGAGGTATTCGCACTGGATACGACGTTGAT | | |
| Cel-miR-794-5p | GTCGTATCCAGTGCAGGGTCCGAGGTATTCGCACTGGATACGACAGTGAC | | |
| Cel-miR-249-3p | GTCGTATCCAGTGCAGGGTCCGAGGTATTCGCACTGGATACGACGGCAAC | | |
| Cel-miR-2-5p | GTCGTATCCAGTGCAGGGTCCGAGGTATTCGCACTGGATACGACCACATC | | |
| Cel-miR-51-3p | GTCGTATCCAGTGCAGGGTCCGAGGTATTCGCACTGGATACGACTGCACC | | |
| Cel-miR-60-5p | GTCGTATCCAGTGCAGGGTCCGAGGTATTCGCACTGGATACGACGATTTT | | |
| Cel-miR-250-5p | GTCGTATCCAGTGCAGGGTCCGAGGTATTCGCACTGGATACGACCGGATC | | |
| Internal control  Cel-miR-64-5p | GTCGTATCCAGTGCAGGGTCCGAGGTATTCGCACTGGATACGACTTCGGT | | |

### **Table S2. List of DE miRNAs (Log2fold change Value)**

| **S. No.** | **miRNA** | **Damaged** | **Response** | **Recovery** |
| --- | --- | --- | --- | --- |
| 1.  | cel-miR-253-3p | 15.217566 | 14.797705 | 15.370711 |
| 2.  | cel-miR-2-5p | 9.320771 | 13.628083 | 13.629298 |
| 3.  | cel-miR-51-3p | 8.472307 | 11.805662 | 12.161905 |
| 4.  | cel-miR-60-5p | 8.096181 | 11.606612 | 11.960069 |
| 5.  | cel-miR-72-3p | 7.932023 | 11.322959 | 10.416706 |
| 6.  | cel-miR-5592-3p | 7.8124 | 8.061744 | 8.689552 |
| 7.  | cel-miR-250-5p | 7.49901 | 6.593667 |  |
| 8.  | cel-miR-790-3p | 7.564031 | 11.221378 | 7.482349 |
| 9.  | cel-miR-55-5p | 7.338282 | 10.079134 | 9.872383 |
| 10.  | cel-miR-71-3p | 7.196851 | 10.715938 | 7.215841 |
| 11.  | cel-miR-241-5p | 6.754145 | 9.984787 | 10.333924 |
| 12.  | cel-miR-44-5p | 6.75016 | 9.623891 | 9.713237 |
| 13.  | cel-miR-49-5p | 6.611946 | 9.711927 | 9.833163 |
| 14.  | cel-miR-90-5p | 6.405039 | 9.296516 | 5.341597 |
| 15.  | cel-miR-37-5p | 6.366608 | 8.701944 | 8.507258 |
| 16.  | cel-miR-1-5p | 6.324934 | 8.735154 | 8.978437 |
| 17.  | cel-miR-355-5p | 6.27949 | 5.552408 |  |
| 18.  | cel-miR-87-5p | 5.890487 | 7.389447 | 4.925573 |
| 19.  | cel-miR-4816-5p | 5.64474 | 4.938969 |  |
| 20.  | cel-miR-229-3p | 5.384355 |  |  |
| 21.  | cel-miR-251 | -5.154795 |  |  |
| 22.  | cel-miR-66-5p | -5.316488 |  |  |
| 23.  | cel-miR-65-5p | -5.674723 |  |  |
| 24.  | cel-miR-234-3p | -7.056085 |  |  |
| 25.  | cel-miR-79-3p | -7.111155 |  |  |
| 26.  | cel-miR-85-3p | -8.445603 |  |  |
| 27.  | cel-miR-45-5p |  | 10.83326 | 9.888896 |
| 28.  | cel-miR-64-3p |  | 9.994151 |  |
| 29.  | cel-miR-259-5p |  | 8.490693 |  |
| 30.  | cel-miR-57-3p |  | 8.162068 | 7.236382 |
| 31.  | cel-miR-793 |  | 7.582245 |  |
| 32.  | cel-miR-58a-5p | 7.45546 |  | 7.086723 |
| 33.  | cel-miR-42-3p |  | 7.300334 | 4.921033 |
| 34.  | cel-miR-228-5p |  | 6.954797 | 6.272276 |
| 35.  | cel-miR-37-3p |  | 6.625447 | 7.599706 |
| 36.  | cel-miR-48-3p | 6.46293 |  |  |
| 37.  | cel-miR-43-3p | 6.24747 |  |  |
| 38.  | cel-miR-76-3p |  | 6.161381 |  |
| 39.  | cel-miR-249-3p |  | 6.100951 | 6.894577 |
| 40.  | cel-miR-242 |  | 5.835626 |  |
| 41.  | cel-miR-228-3p |  | 5.755846 |  |
| 42.  | cel-miR-356b-3p |  | 5.591607 |  |
| 43.  | cel-miR-77-5p |  | 5.465642 |  |
| 44.  | cel-lin-4-3p |  | 5.459512 |  |
| 45.  | cel-miR-229-5p | 5.32619 |  |  |
| 46.  | cel-miR-74-5p |  | 5.183204 |  |
| 47.  | cel-miR-240-3p |  | 5.139177 | 5.796168 |
| 48.  | cel-miR-786-3p |  | 5.026322 |  |
| 49.  | cel-miR-39-5p |  |  | 8.042953 |
| 50.  | cel-miR-357-3p |  |  | 6.773676 |
| 51.  | cel-miR-8200-3p |  |  | 6.719894 |
| 52.  | cel-miR-244-5p |  |  | 6.040286 |
| 53.  | cel-miR-790-5p |  |  | 5.928921 |
| 54.  | cel-miR-63-5p |  |  | 5.427385 |
| 55.  | cel-miR-246-3p |  |  | 5.307335 |
| 56.  | cel-miR-38-3p | 13.663014 | 12.704585 | 12.843285 |
| 57.  | cel-miR-54-5p | 11.703844 | 10.002276 | 9.749616 |

**Table S3.** **Real-time PCR was performed using the following standard conditions**

| **No. of Cycles** | **Time** | **Temperature** |
| --- | --- | --- |
| 1 | 2 min. | 50℃ |
| 1 | 10 min. | 95℃ |
| 40X | 15 sec. | 95℃ |
|  | 1 min. | 60℃ |
| Melt Curve | 0.2℃ increment per second | 60℃ to 95℃ |

**Table S4**. **Stem-loop qRT-PCR reaction set-up**

| **qRT-PCR Reaction mixture** | **Sample** | **No Template Control (NTC)** |
| --- | --- | --- |
| PowerUp SYBR green master mix (2X) | 5 ul | 5 ul |
| *Specific forward primer (1 uM) | 0.5 ul | 0.5 ul |
| Universal Reverse primer (1 uM) | 0.5 ul | 0.5 ul |
| *C-DNA template  (1:5 dilution of 10ng/ul) | 1 ul | 0 ul |
| NFW | 3 ul | 4 ul |
| Total | 10 ul | 10 ul |

* Represents specific forward primers for miRNAs and specific cDNA.

**Table S5: Cross-species sequence relationships and biological context of DE miRNAs in *C. elegans* and humans**

| **S. No.** | ***C. elegans* miRNA** | **Human sequence-related miRNA(s)** | **Similarity criterion** | **Reported human biological function/relevance** | **Reported *C. elegans* biological association** | **References for *C.elegans*** |
| --- | --- | --- | --- | --- | --- | --- |
| 1 | cel-miR-253-3p | — | — | — | Dietary-restriction-associated metabolic remodelling | [1] |
| 2 | cel-miR-38-3p | — | — | — | Dietary-restriction-associated metabolic remodelling | [1] |
| 3 | cel-miR-54-5p | hsa-miR-99a; hsa-miR-99b; hsa-miR-100 | 5′ ≥7/10 nt continuous similarity | Cellular differentiation, metabolism, proliferation and disease-associated regulation | Member of the miR-51 family; developmental association | [2] |
| 4 | cel-miR-2-5p | hsa-miR-499-3p | 5′ ≥7/10 nt continuous similarity | Muscle/cardiac and cellular regulatory contexts | miR-2 family sequence group | [3] |
| 5 | cel-miR-51-3p | hsa-miR-99a; hsa-miR-99b; hsa-miR-100 | 5′ ≥7/10 nt continuous similarity + ≥70% full-length similarity | Cellular differentiation, metabolism, proliferation and disease-associated regulation | Embryonic development and pharyngeal attachment | [2] |
| 6 | cel-miR-60-5p | — | — | — | Stress/oxidative-response context | [4] |
| 7 | cel-miR-72-3p | hsa-miR-31 | 5′ ≥7/10 nt continuous similarity + ≥70% full-length similarity | Cell proliferation, differentiation, migration and disease-associated regulation | Fertility, survival, lifespan and mitochondrial phenotypes | [5] |
| 8 | cel-miR-5592-3p | — | — | — | — | — |
| 9 | cel-miR-250-5p | hsa-miR-27a; hsa-miR-27b; hsa-miR-128; hsa-miR-499-3p; hsa-miR-768-3p | 5′ ≥7/10 nt continuous similarity | Metabolic, developmental and cellular regulatory contexts | miR-2-related sequence family | [6] |
| 10 | cel-miR-790-3p | — | — | — | miR-63/64/65/66/228/229 sequence family | — |
| 11 | cel-miR-55-5p | hsa-miR-99a; hsa-miR-99b; hsa-miR-100 | 5′ ≥7/10 nt continuous similarity | Cellular differentiation, metabolism, proliferation and disease-associated regulation | miR-51 family | [2] |
| 12 | cel-miR-71-3p | — | — | — | Longevity and stress resistance | [7] |
| 13 | cel-miR-241-5p | hsa-let-7a; hsa-let-7b; hsa-let-7c; hsa-let-7d; hsa-let-7e; hsa-let-7f; hsa-let-7g; hsa-let-7i; hsa-miR-98 | 5′ ≥7/10 nt continuous similarity | Developmental timing, differentiation, metabolism and disease-associated regulation | Developmental timing | [8] |
| 14 | cel-miR-44-5p | hsa-miR-134; hsa-miR-708* | 5′ ≥7/10 nt continuous similarity | Neural, developmental and cellular regulation | Germline and sperm specification | [9] |
| 15 | cel-miR-49-5p | hsa-miR-21*; hsa-miR-29a; hsa-miR-29b; hsa-miR-29c; hsa-miR-593* | 5′ ≥7/10 nt continuous similarity | Stress, immune, fibrotic and disease-associated regulation | Stress-associated miRNA context | [10] |
| 16 | cel-miR-90-5p | hsa-miR-190; hsa-miR-190b | 5′ ≥7/10 nt continuous similarity | Cellular stress, neuronal and developmental regulation | — | — |
| 17 | cel-miR-37-5p | — | — | — | — | — |
| 18 | cel-miR-1-5p | hsa-miR-1; hsa-miR-122; hsa-miR-206 | 5′ ≥7/10 nt continuous similarity + ≥70% full-length similarity | Muscle, metabolic and tissue-specific regulatory functions | Muscle and neuromuscular regulation | [11] |
| 19 | cel-miR-355-5p | — | — | — | Innate immunity; DAF-2/p38 MAPK-associated regulation | [12] |
| 20 | cel-miR-87-5p | — | — | — | — | — |
| 21 | cel-miR-4816-5p | — | — | — | — |  |
| 22 | cel-miR-229-3p | hsa-miR-96; hsa-miR-183; hsa-miR-200a; hsa-miR-514 | 5′ ≥7/10 nt continuous similarity | Sensory, epithelial and developmental regulation | Longevity/stress-associated miR-229 family | [13] |
| 23 | cel-miR-251 | hsa-miR-26a; hsa-miR-26b | 5′ ≥7/10 nt continuous similarity | Immune, metabolic and cellular regulation | Innate immunity during fungal infection | [14] |
| 24 | cel-miR-66-5p | hsa-miR-96; hsa-miR-183; hsa-miR-200a; hsa-miR-514 | 5′ ≥7/10 nt continuous similarity | Sensory, epithelial and developmental regulation | Heat-stress response | [10] |
| 25 | cel-miR-65-5p | hsa-miR-96; hsa-miR-183; hsa-miR-200a; hsa-miR-514 | 5′ ≥7/10 nt continuous similarity | Sensory, epithelial and developmental regulation | Heat-stress response | [10] |
| 26 | cel-miR-234-3p | hsa-miR-126*; hsa-miR-137 | 5′ ≥7/10 nt continuous similarity + ≥70% full-length similarity | Vascular, neural and cellular regulatory functions | Neuropeptide release at the neuromuscular junction | [15] |
| 27 | cel-miR-79-3p | hsa-miR-7; hsa-miR-9*; hsa-miR-320; hsa-miR-340; hsa-miR-548a-3p | 5′ ≥7/10 nt continuous similarity + ≥70% full-length similarity | Neural, epithelial and developmental regulatory contexts | Epidermal regulation and neuronal migration | [16] |
| 28 | cel-miR-85-3p | — | — | — | Heat-shock recovery and HSP-70 regulation | [17] |
| 29 | cel-miR-45-5p | hsa-miR-134; hsa-miR-708* | 5′ ≥7/10 nt continuous similarity | Neural, developmental and cellular regulation | Germline and sperm specification | [9] |
| 30 | cel-miR-64-3p | hsa-miR-96; hsa-miR-183; hsa-miR-200a; hsa-miR-514 | 5′ ≥7/10 nt continuous similarity | Sensory, epithelial and developmental regulation | Heat/stress-associated miRNA family | [10] |
| 31 | cel-miR-259-5p | hsa-miR-216a; hsa-miR-216b | 5′ ≥7/10 nt continuous similarity | Metabolic and cellular regulation | — | — |
| 32 | cel-miR-57-3p | hsa-miR-10a; hsa-miR-10b; hsa-miR-146b-3p; hsa-miR-99a; hsa-miR-100 | 5′ ≥7/10 nt continuous similarity + ≥70% full-length similarity | Developmental, immune, stress and metabolic regulatory contexts | Lifespan/fertility/mitochondrial association in recent work | [5] |
| 33 | cel-miR-793 | hsa-let-7a; hsa-let-7b; hsa-let-7c; hsa-let-7e; hsa-let-7f; hsa-let-7g; hsa-let-7i; hsa-miR-98; hsa-miR-202 | 5′ ≥7/10 nt continuous similarity + ≥70% full-length similarity | Human biological relevance varies among the listed sequence-related miRNAs; no single function is assigned here. | let-7-related developmental context | [8] |
| 34 | cel-miR-58a-5p | hsa-miR-450b-3p | 5′ ≥7/10 nt continuous similarity | Cellular differentiation and developmental regulatory contexts | Stress/proteostasis-associated context | [1] |
| 35 | cel-miR-42-3p | — | — | — | — | — |
| 36 | cel-miR-228-5p | hsa-miR-96; hsa-miR-183; hsa-miR-200a; hsa-miR-514 | 5′ ≥7/10 nt continuous similarity | Sensory, epithelial and developmental regulation | Longevity/stress-associated miRNA family | [13] |
| 37 | cel-miR-37-3p | — | — | — | — | — |
| 38 | cel-miR-48-3p | hsa-let-7a; hsa-let-7b; hsa-let-7c; hsa-let-7d; hsa-let-7e; hsa-let-7f; hsa-let-7g; hsa-let-7i; hsa-miR-98 | 5′ ≥7/10 nt continuous similarity | Developmental timing, differentiation, metabolism and disease-associated regulation | Developmental timing | [8] |
| 39 | cel-miR-43-3p | hsa-miR-27a; hsa-miR-27b; hsa-miR-128; hsa-miR-499-3p; hsa-miR-768-3p | 5′ ≥7/10 nt continuous similarity | Metabolic, developmental and cellular regulatory contexts | miR-2-related sequence family | — |
| 40 | cel-miR-76-3p | — | — | — | — |  |
| 41 | cel-miR-249-3p | — | — | — | Fertility, survival, lifespan and mitochondrial association | [5] |
| 42 | cel-miR-242 | — | — | — | — | — |
| 43 | cel-miR-228-3p | hsa-miR-96; hsa-miR-183; hsa-miR-200a; hsa-miR-514 | 5′ ≥7/10 nt continuous similarity | Sensory, epithelial and developmental regulation | Longevity/stress-associated miRNA family | [13] |
| 44 | cel-miR-356b-3p | — | — | — | — | — |
| 45 | cel-miR-77-5p | — | — | — | Fertility, survival, lifespan and mitochondrial association | [5] |
| 46 | cel-lin-4-3p | hsa-miR-125a-5p; hsa-miR-125b; hsa-miR-331-3p | 5′ ≥7/10 nt continuous similarity + ≥70% full-length similarity | Development, differentiation, immune and disease-associated regulation | Developmental timing through LIN-14/LIN-28 regulation | [3] |
| 47 | cel-miR-229-5p | hsa-miR-96; hsa-miR-183; hsa-miR-200a; hsa-miR-514 | 5′ ≥7/10 nt continuous similarity | Sensory, epithelial and developmental regulation | Longevity/stress-associated miR-229 family | [13] |
| 48 | cel-miR-74-5p | hsa-miR-31; hsa-miR-513b; hsa-miR-873 | 5′ ≥7/10 nt continuous similarity | Human biological relevance varies among the listed sequence-related miRNAs; no single function is assigned here. | miR-72/73/74 sequence family | [1] |
| 49 | cel-miR-240-3p | hsa-miR-193a-3p; hsa-miR-193b | 5′ ≥7/10 nt continuous similarity + ≥70% full-length similarity | Metabolic, stress and cellular regulatory functions | Metabolic/defecation-associated context | [6] |
| 50 | cel-miR-786-3p | hsa-miR-18a*; hsa-miR-18b*; hsa-miR-365 | 5′ ≥7/10 nt continuous similarity | Metabolic, developmental and cellular stress-associated regulation | Intestinal Ca2+-wave initiation and defecation rhythm | [18] |
| 51 | cel-miR-39-5p | — | — | — | — | — |
| 52 | cel-miR-357-3p | hsa-miR-302a; hsa-miR-302b; hsa-miR-302c; hsa-miR-302d | 5′ ≥7/10 nt continuous similarity | Pluripotency, stem-cell and developmental regulation | — | — |
| 53 | cel-miR-8200-3p | — | — | — | — | — |
| 54 | cel-miR-244-5p | hsa-miR-9 | 5′ ≥7/10 nt continuous similarity | Neural development and differentiation | — | — |
| 55 | cel-miR-790-5p | hsa-miR-96; hsa-miR-183; hsa-miR-200a; hsa-miR-514 | 5′ ≥7/10 nt continuous similarity | Sensory, epithelial and developmental regulation | miR-63/64/65/66/228/229 sequence family | [19] |
| 56 | cel-miR-63-5p | hsa-miR-96; hsa-miR-183; hsa-miR-200a; hsa-miR-514 | 5′ ≥7/10 nt continuous similarity | Sensory, epithelial and developmental regulation | miR-63/64/65/66 family; stress/longevity context | [1] |
| 57 | cel-miR-246-3p | — | — | — | Lifespan/stress-associated phenotype | [13] |

The above human sequence-related miRNAs are reported only where the *C. elegans* precursor/family appears in [20], Table 4.

Ibáñez-Ventoso et al. classified miRNAs as sequence-related using either

(i) at least 7 continuous identical nucleotides within the first 10 nucleotides of the mature miRNA, with no gaps or mismatches except G·U pairing, or
(ii) ≥70% identity across the mature miRNA sequence.
(iii) MiRNAs showing 60–69.9% full-length identity were reported separately as weaker relationships and are not included here as primary sequence-related matches.
