## Supplementary file 2 for "Integrated miRNA-protein profiling of small extracellular vesicle cargo reveals dynamic cargo remodelling during tissue repair in UV-C-damaged *C. elegans*"

**miRge3.0: Comprehensive analysis of small RNA sequencing Data.**

SmallRNA distribution | Read Length | isomiR results | Abundant miRNAs | UMI distribution | Novel miRNAs

Cumulative isomiR variant type distribution of the samples

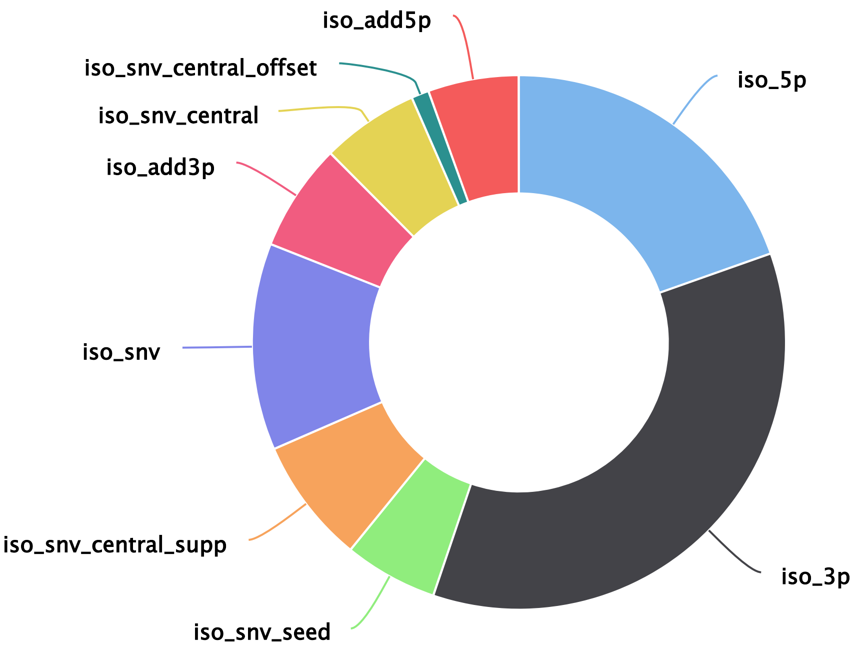


Read distribution of isomiRs for the top 20 abundant miRNAs: miR_A1_DU_R1

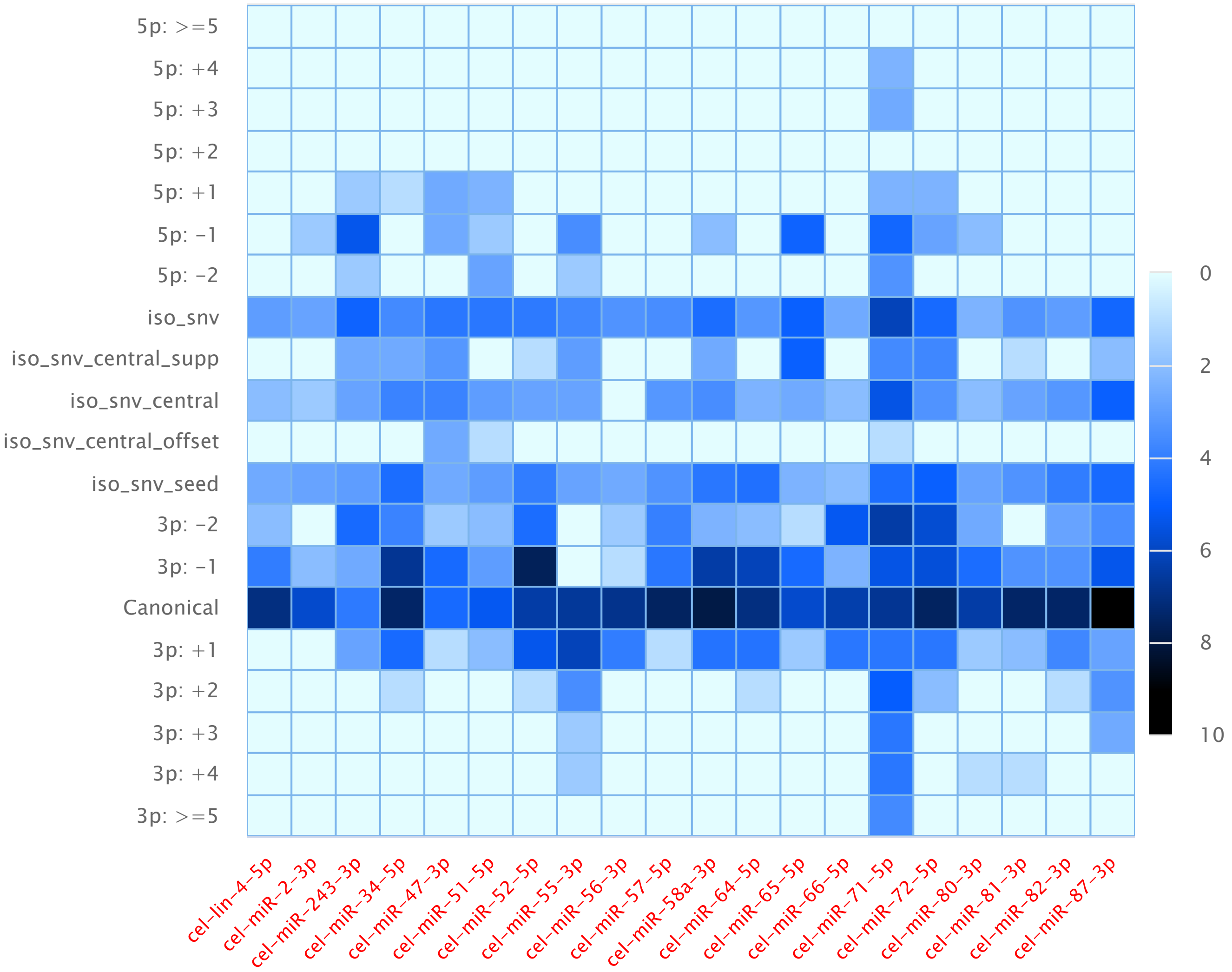

Read distribution of isomiRs for the top 20 abundant miRNAs: miR_A2_DU_R1

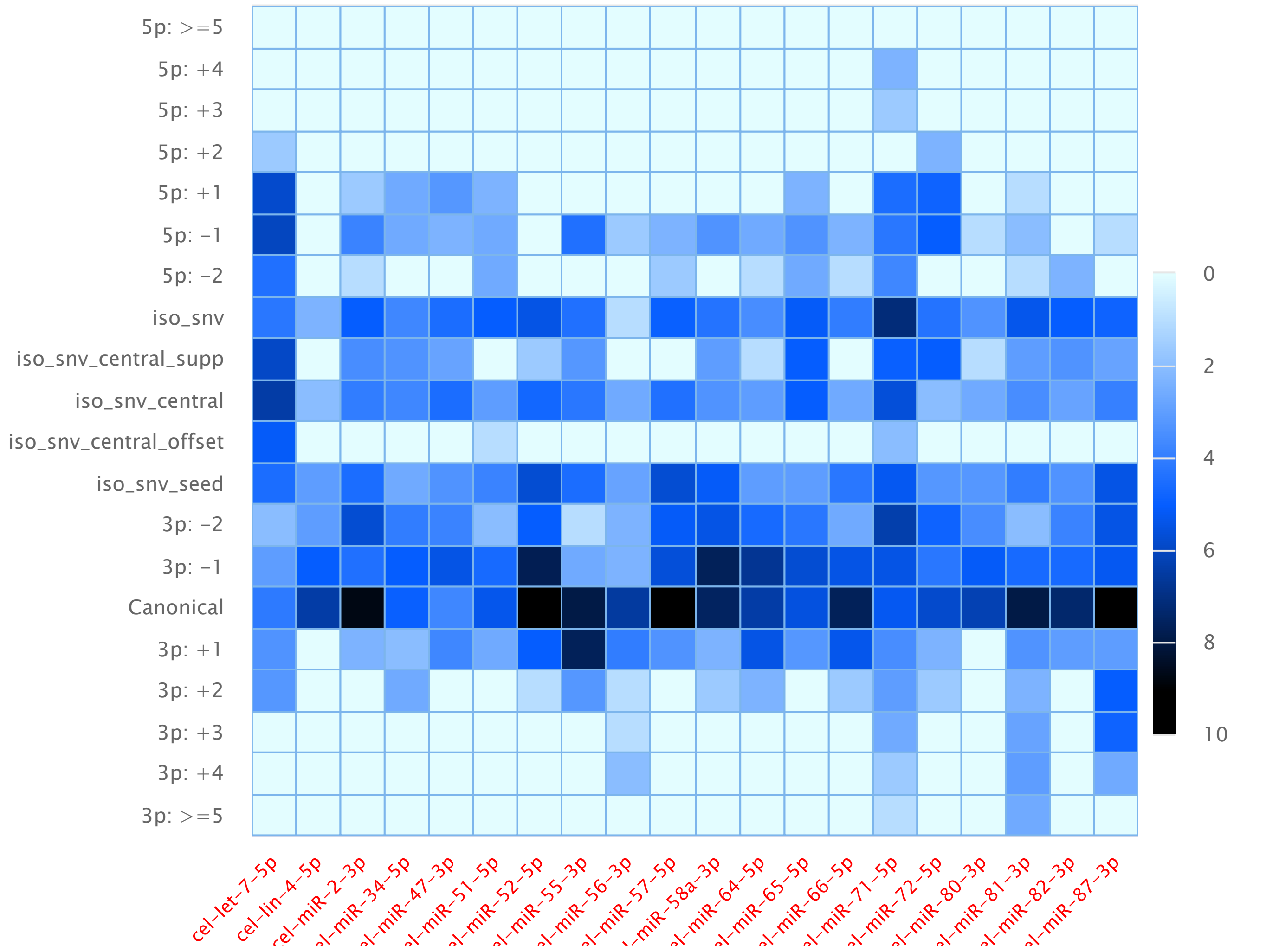


Read distribution of isomiRs for the top 20 abundant miRNAs: miR_A3_DU_R1

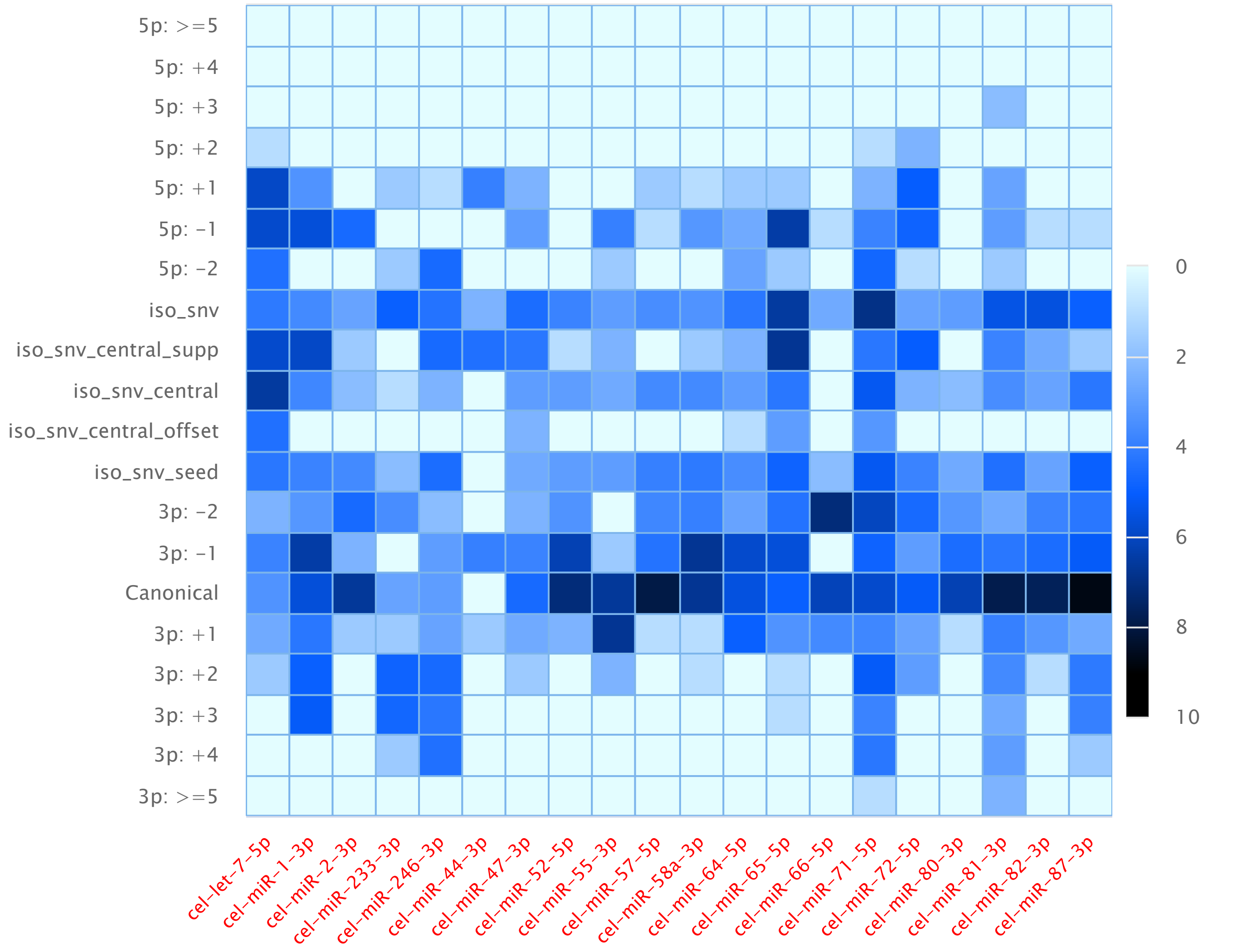

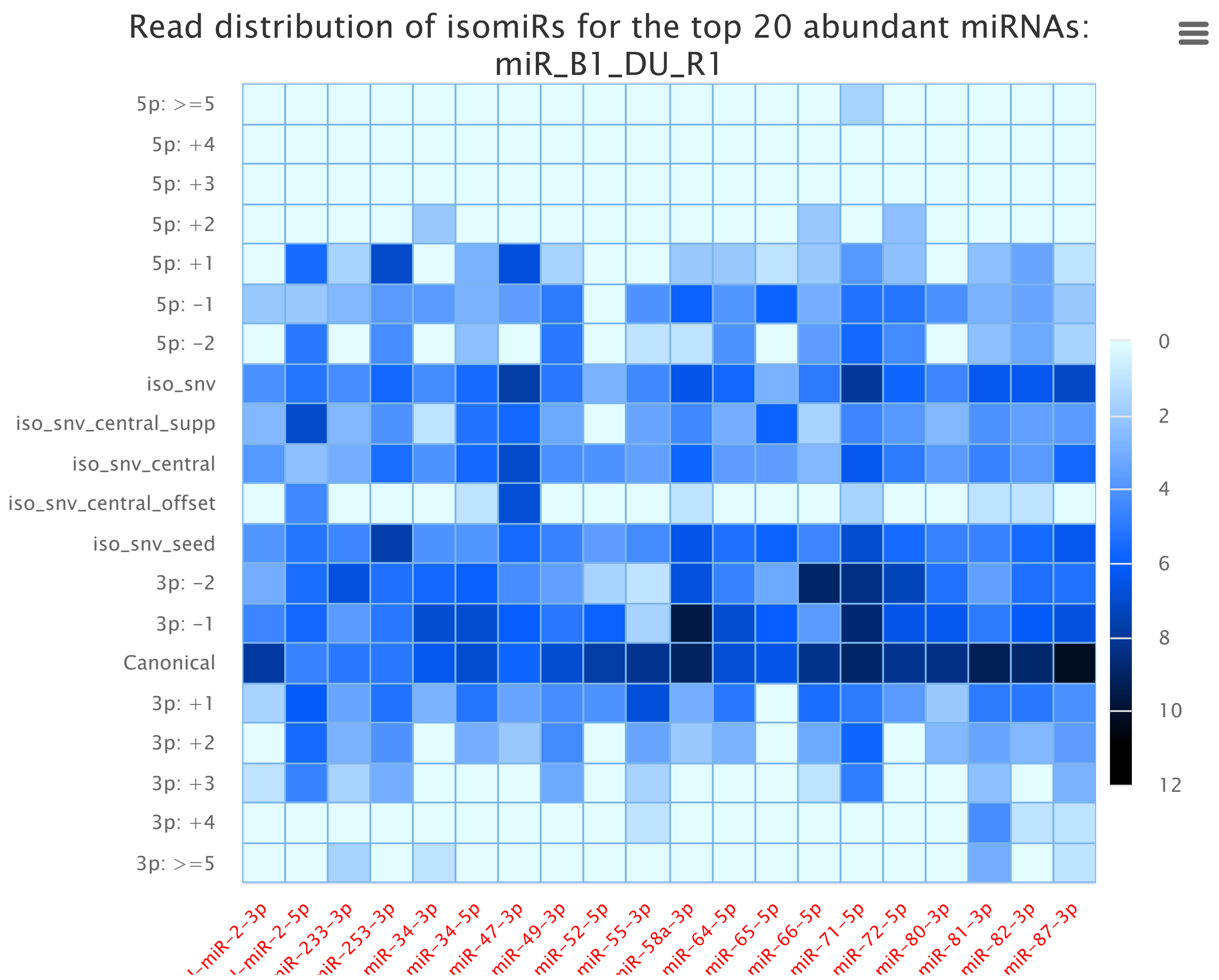
Read distribution of isomiRs for the top 20 abundant miRNAs: miR_B1_DU_R1
5p: >=5
5p: +4
5p: +3
5p: +2
5p: +1
5p: −1
5p: −2
iso_snv
iso_snv_central_supp
iso_snv_central
iso_snv_central_offset
iso_snv_seed
3p: −2
3p: −1
Canonical
3p: +1
3p: +2
3p: +3
3p: +4
3p: >=5
0
2
4
6
8
10
12
miR−2−3p
miR−2−5p
iR−233−3p
iR−253−3p
miR−34−3p
miR−34−5p
miR−47−3p
miR−49−3p
miR−52−5p
miR−55−3p
iR−58a−3p
miR−64−5p
miR−65−5p
miR−66−5p
miR−71−5p
miR−72−5p
miR−80−3p
miR−81−3p
miR−82−3p
miR−87−3p

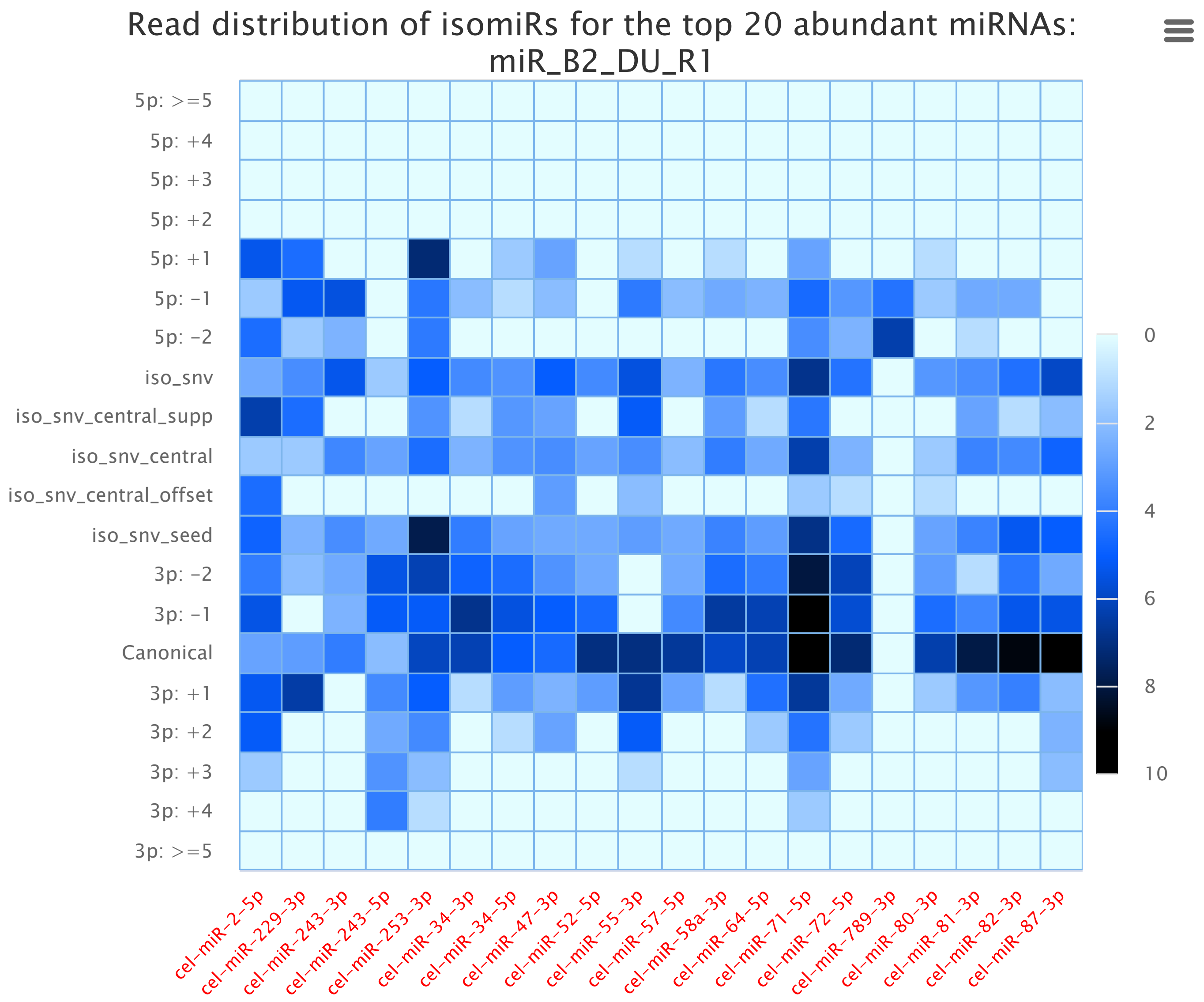
Read distribution of isomiRs for the top 20 abundant miRNAs: miR_B2_DU_R1
5p: >=5
5p: +4
5p: +3
5p: +2
5p: +1
5p: −1
5p: −2
iso_snv
iso_snv_central_supp
iso_snv_central
iso_snv_central_offset
iso_snv_seed
3p: −2
3p: −1
Canonical
3p: +1
3p: +2
3p: +3
3p: +4
3p: >=5
0
2
4
6
8
10
cel−miR−2−5p
cel−miR−229−3p
cel−miR−243−3p
cel−miR−243−5p
cel−miR−253−3p
cel−miR−34−3p
cel−miR−34−5p
cel−miR−47−3p
cel−miR−52−5p
cel−miR−55−3p
cel−miR−57−5p
cel−miR−58a−3p
cel−miR−64−5p
cel−miR−71−5p
cel−miR−72−5p
cel−miR−789−3p
cel−miR−80−3p
cel−miR−81−3p
cel−miR−82−3p
cel−miR−87−3p

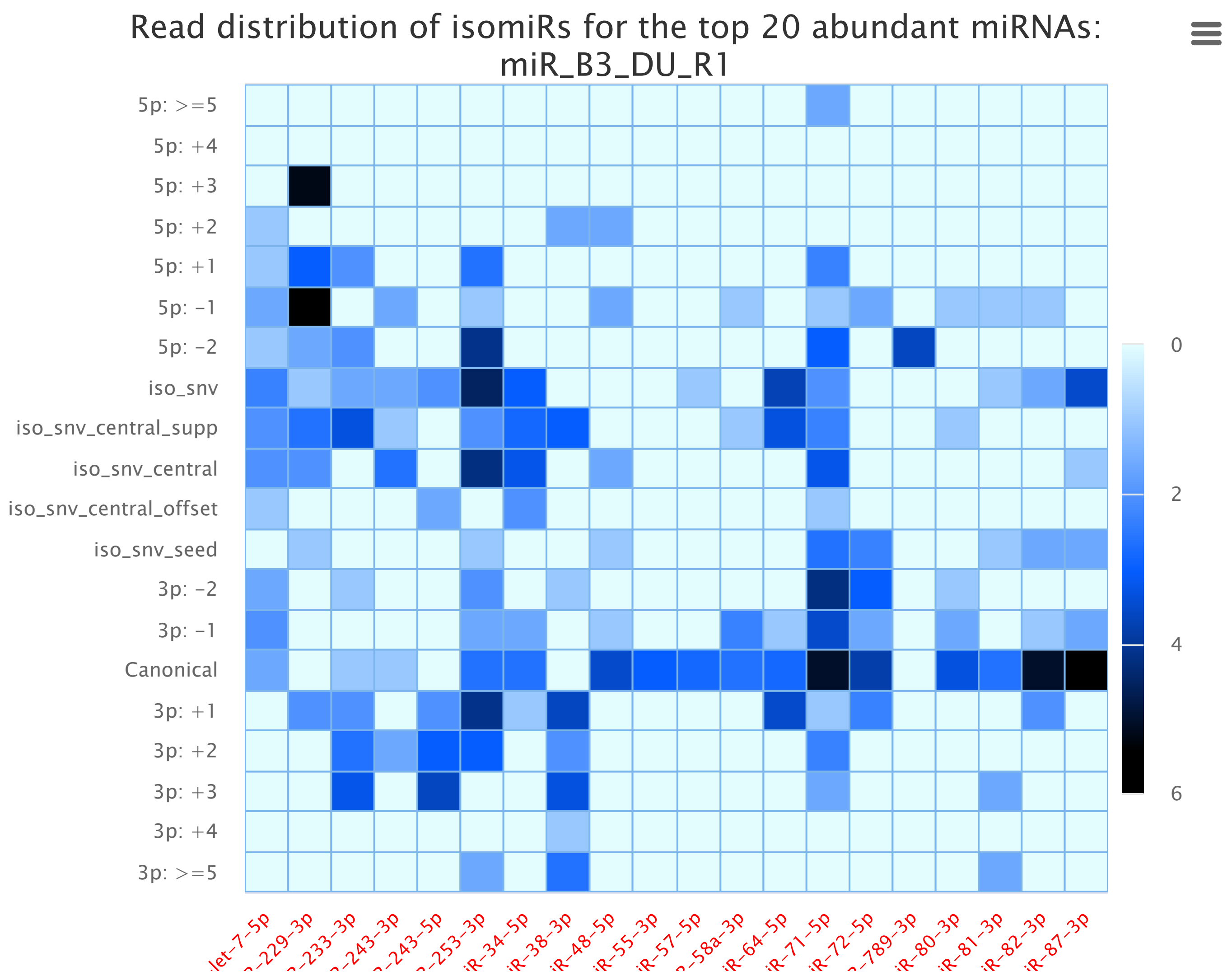
Read distribution of isomiRs for the top 20 abundant miRNAs: miR_B3_DU_R1
5p: >=5
5p: +4
5p: +3
5p: +2
5p: +1
5p: -1
5p: -2
iso_snv
iso_snv_central_supp
iso_snv_central
iso_snv_central_offset
iso_snv_seed
3p: -2
3p: -1
Canonical
3p: +1
3p: +2
3p: +3
3p: +4
3p: >=5
0
2
4
6

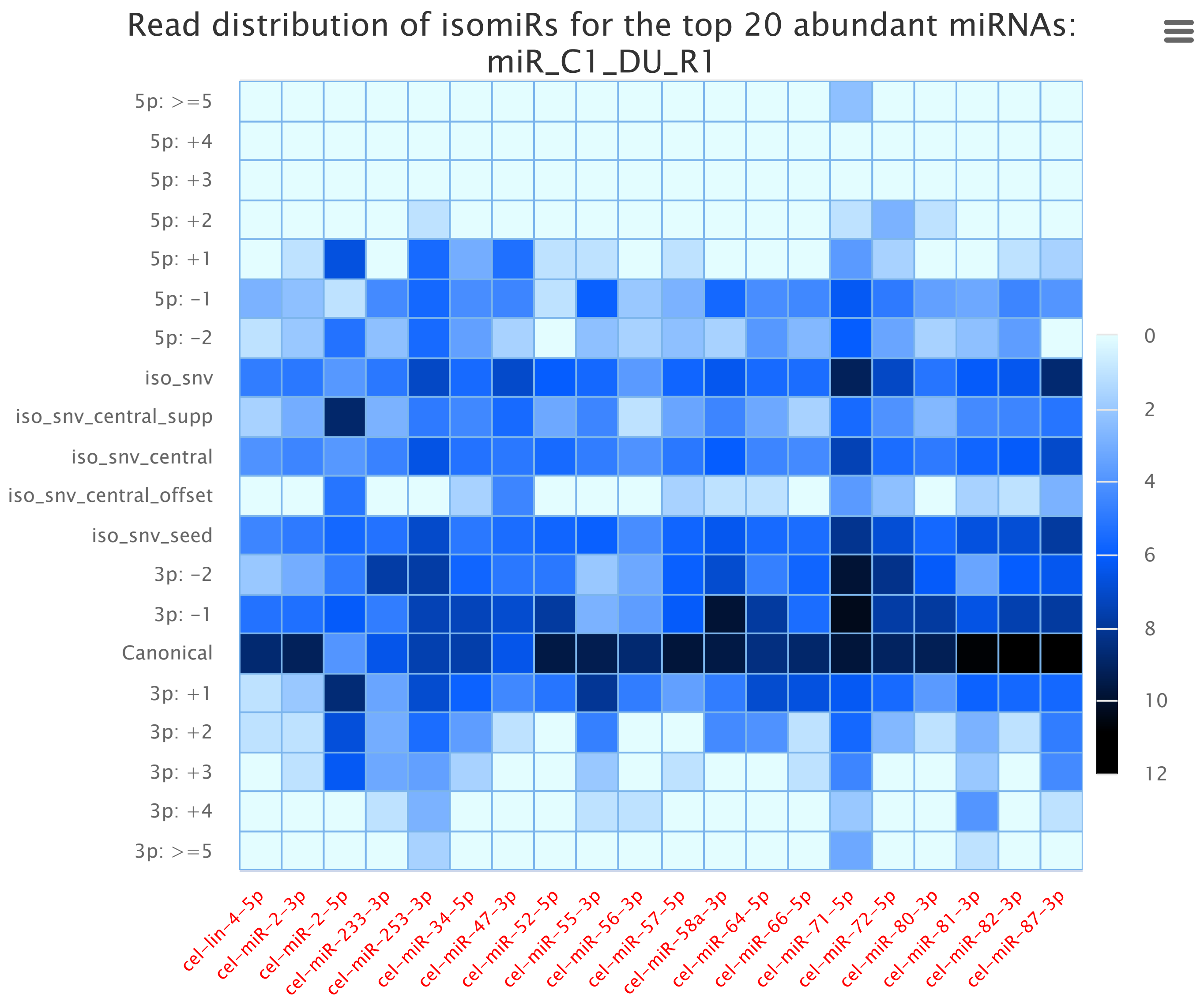
Read distribution of isomiRs for the top 20 abundant miRNAs: miR_C1_DU_R1
5p: >=5
5p: +4
5p: +3
5p: +2
5p: +1
5p: -1
5p: -2
iso_snv
iso_snv_central_supp
iso_snv_central
iso_snv_central_offset
iso_snv_seed
3p: -2
3p: -1
Canonical
3p: +1
3p: +2
3p: +3
3p: +4
3p: >=5
0
2
4
6
8
10
12
cel-lin-4-5p
cel-miR-2-3p
cel-miR-2-5p
cel-miR-233-3p
cel-miR-253-3p
cel-miR-34-5p
cel-miR-47-3p
cel-miR-52-5p
cel-miR-55-3p
cel-miR-56-3p
cel-miR-57-5p
cel-miR-58a-3p
cel-miR-64-5p
cel-miR-66-5p
cel-miR-71-5p
cel-miR-72-5p
cel-miR-80-3p
cel-miR-81-3p
cel-miR-82-3p
cel-miR-87-3p

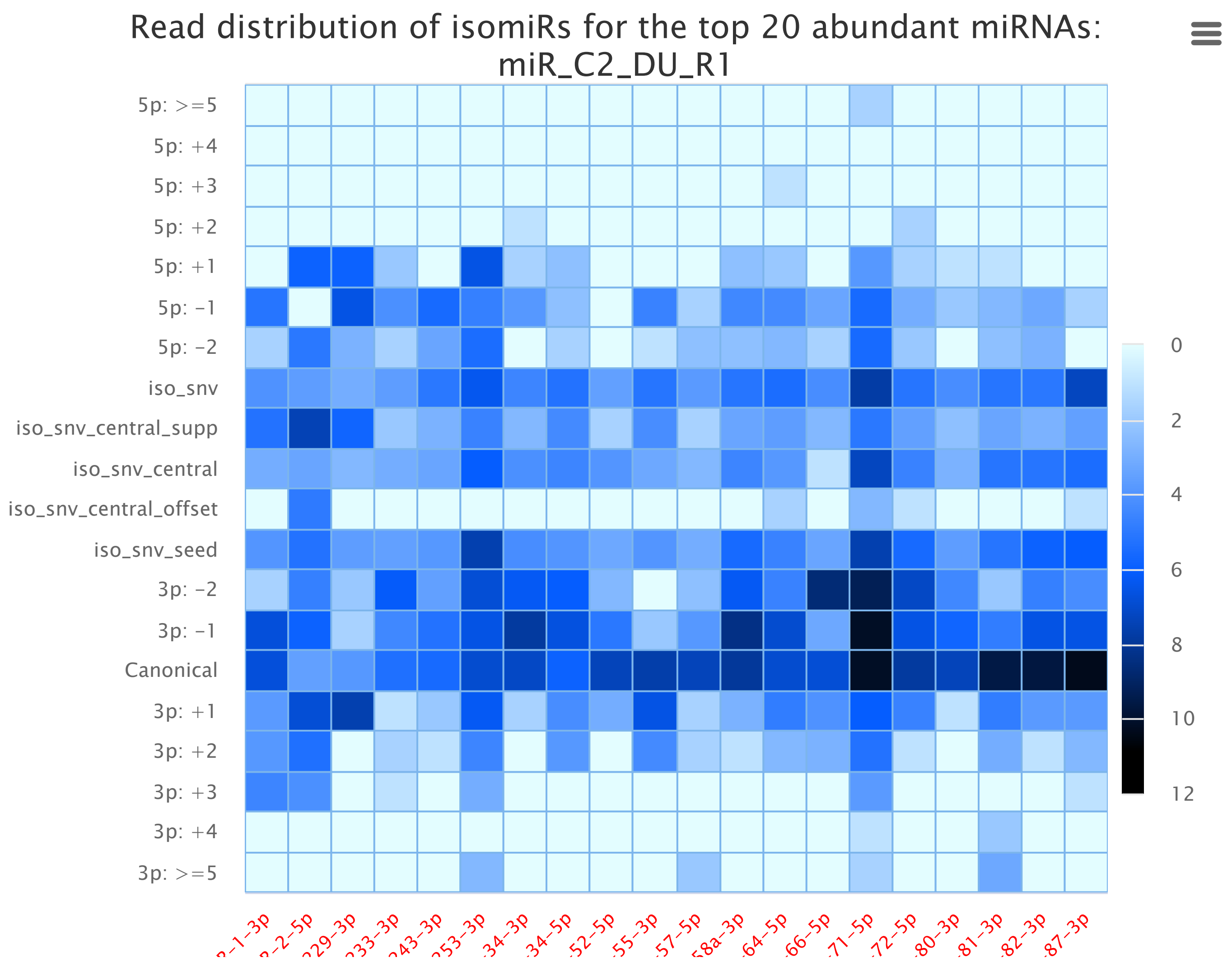
Read distribution of isomiRs for the top 20 abundant miRNAs:
miR_C2_DU_R1
5p: >=5
5p: +4
5p: +3
5p: +2
5p: +1
5p: -1
5p: -2
iso_snv
iso_snv_central_supp
iso_snv_central
iso_snv_central_offset
iso_snv_seed
3p: -2
3p: -1
Canonical
3p: +1
3p: +2
3p: +3
3p: +4
3p: >=5
0
2
4
6
8
10
12
-1-3p
-2-5p
29-3p
33-3p
43-3p
53-3p
34-3p
34-5p
52-5p
55-3p
57-5p
8a-3p
64-5p
66-5p
71-5p
72-5p
80-3p
81-3p
82-3p
87-3p

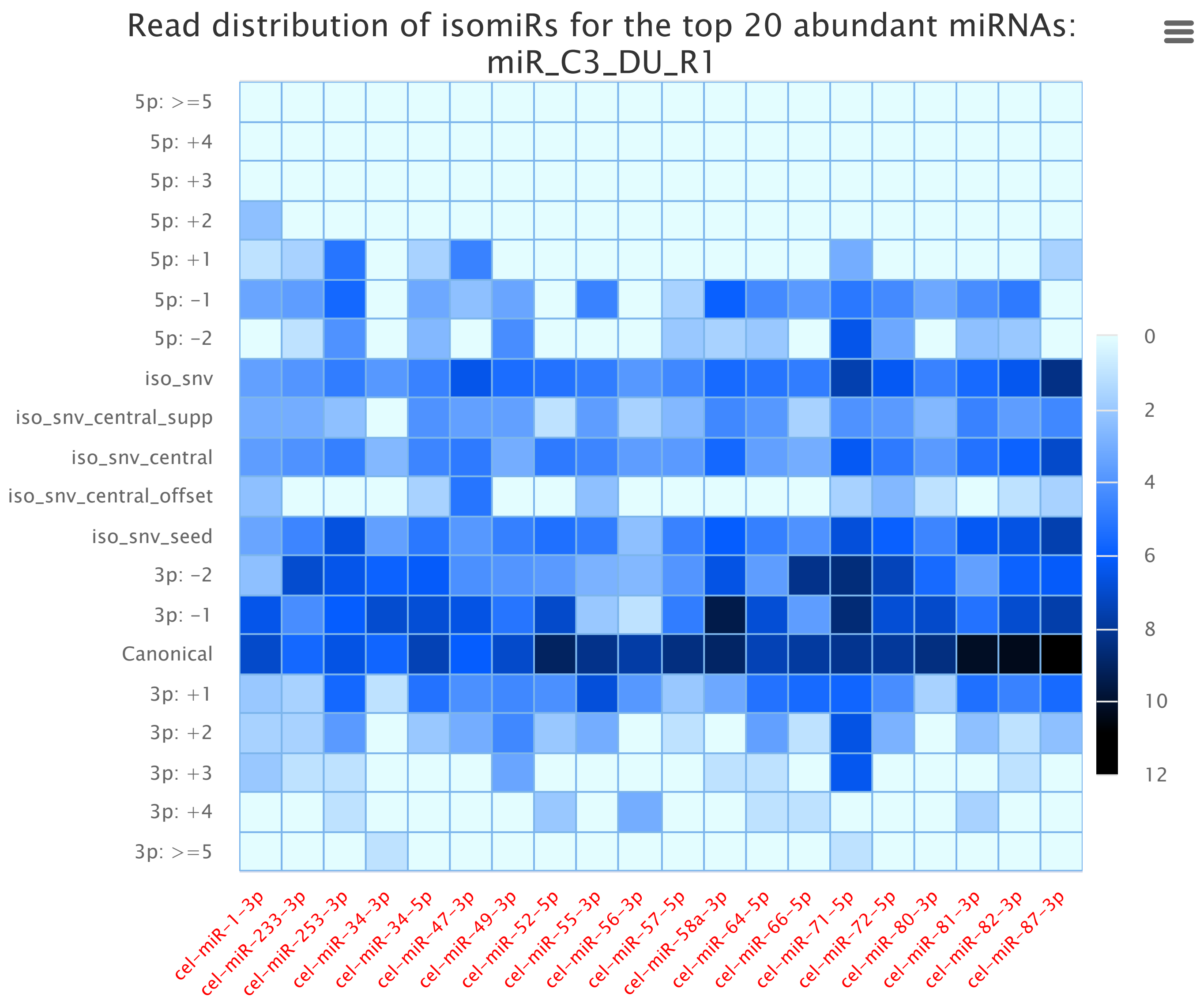
Read distribution of isomiRs for the top 20 abundant miRNAs:
miR_C3_DU_R1
5p: >=5
5p: +4
5p: +3
5p: +2
5p: +1
5p: -1
5p: -2
iso_snv
iso_snv_central_supp
iso_snv_central
iso_snv_central_offset
iso_snv_seed
3p: -2
3p: -1
Canonical
3p: +1
3p: +2
3p: +3
3p: +4
3p: >=5
0
2
4
6
8
10
12
cel-miR-1-3p
cel-miR-233-3p
cel-miR-253-3p
cel-miR-34-3p
cel-miR-34-5p
cel-miR-47-3p
cel-miR-49-3p
cel-miR-52-5p
cel-miR-55-3p
cel-miR-56-3p
cel-miR-57-5p
cel-miR-58a-3p
cel-miR-64-5p
cel-miR-66-5p
cel-miR-71-5p
cel-miR-72-5p
cel-miR-80-3p
cel-miR-81-3p
cel-miR-82-3p
cel-miR-87-3p

Read distribution of isomiRs for the top 20 abundant miRNAs: miR_D1_DU_R1

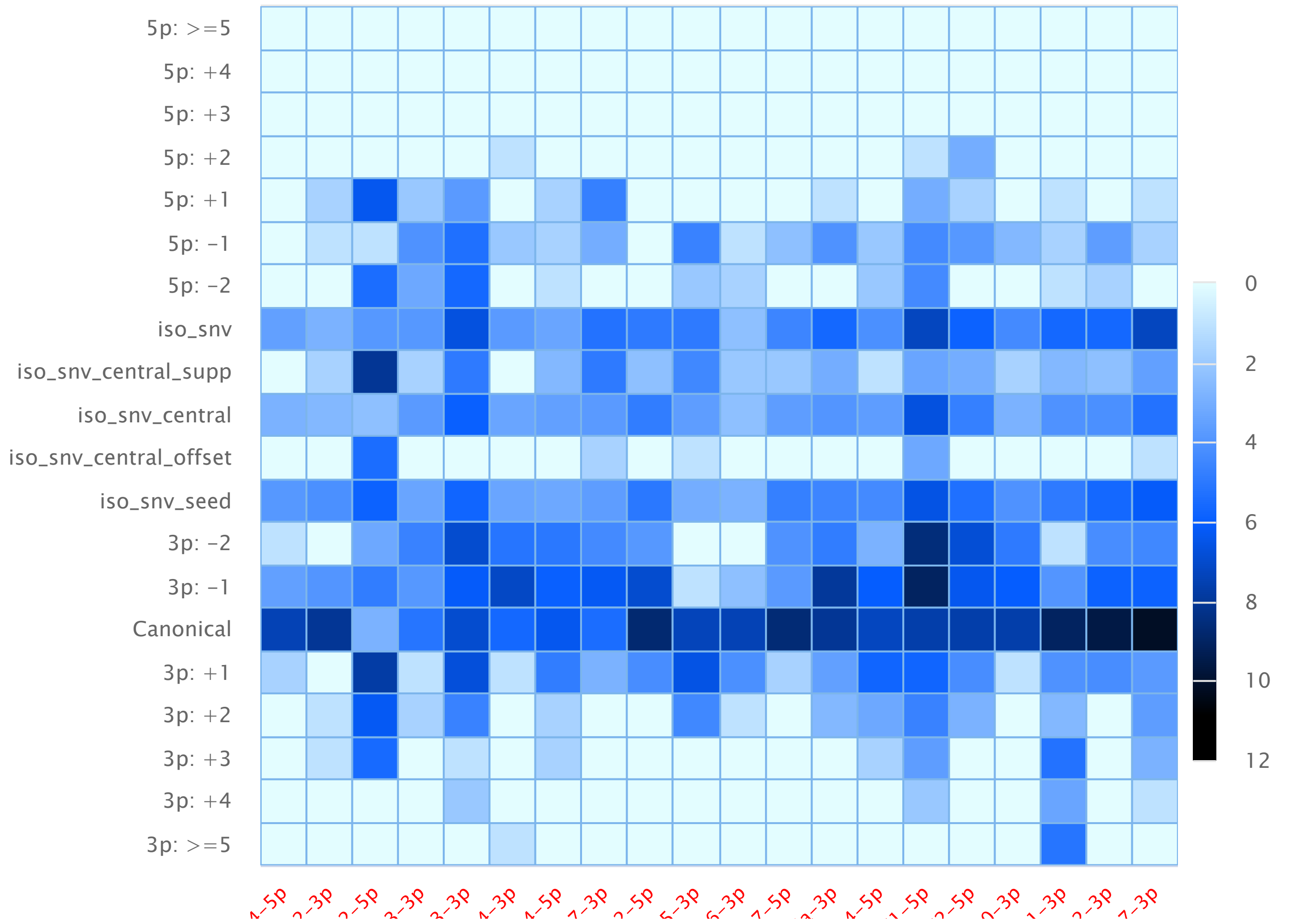


Read distribution of isomiRs for the top 20 abundant miRNAs: miR_D2_DU_R1

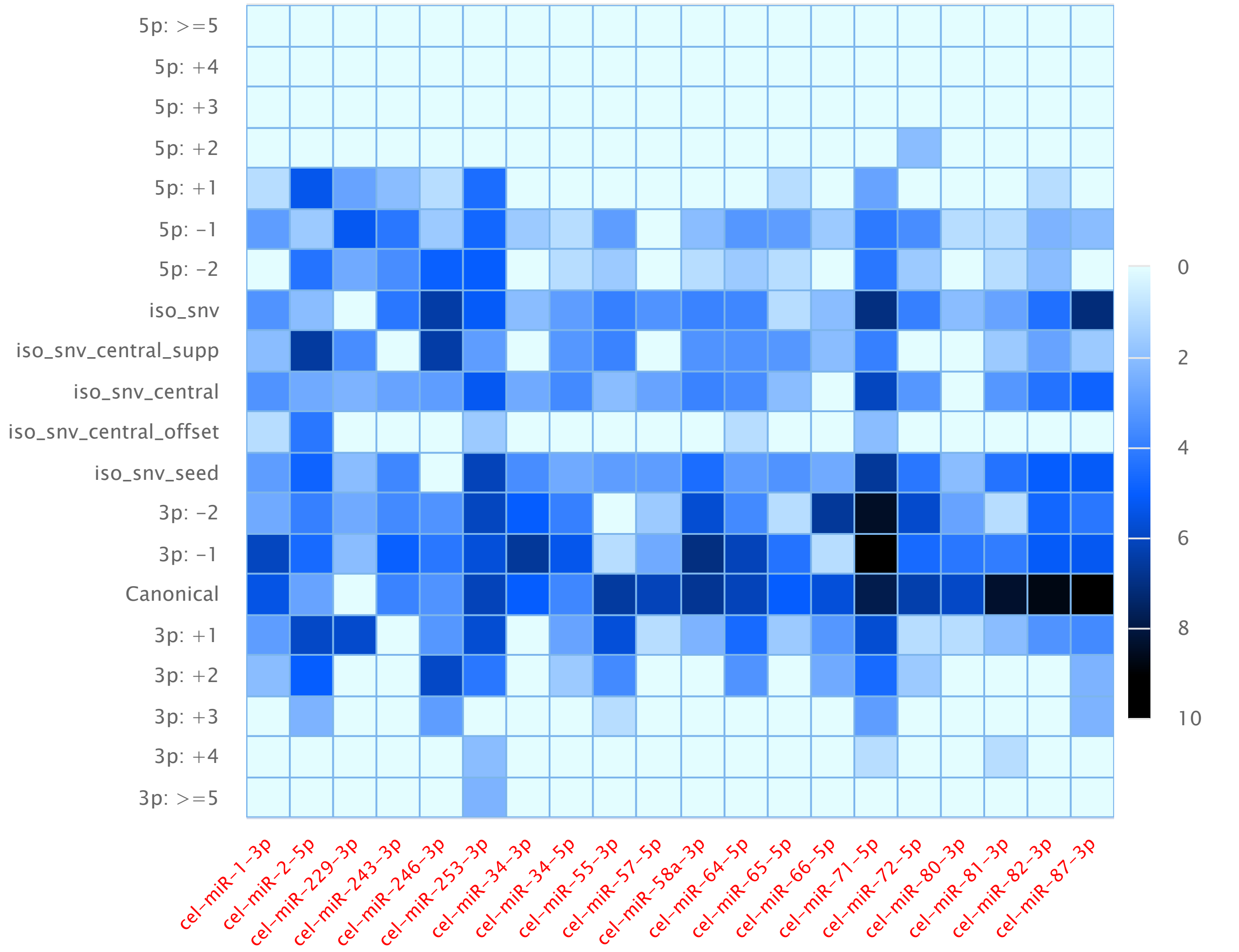

Read distribution of isomiRs for the top 20 abundant miRNAs: miR_D3_DU_R1

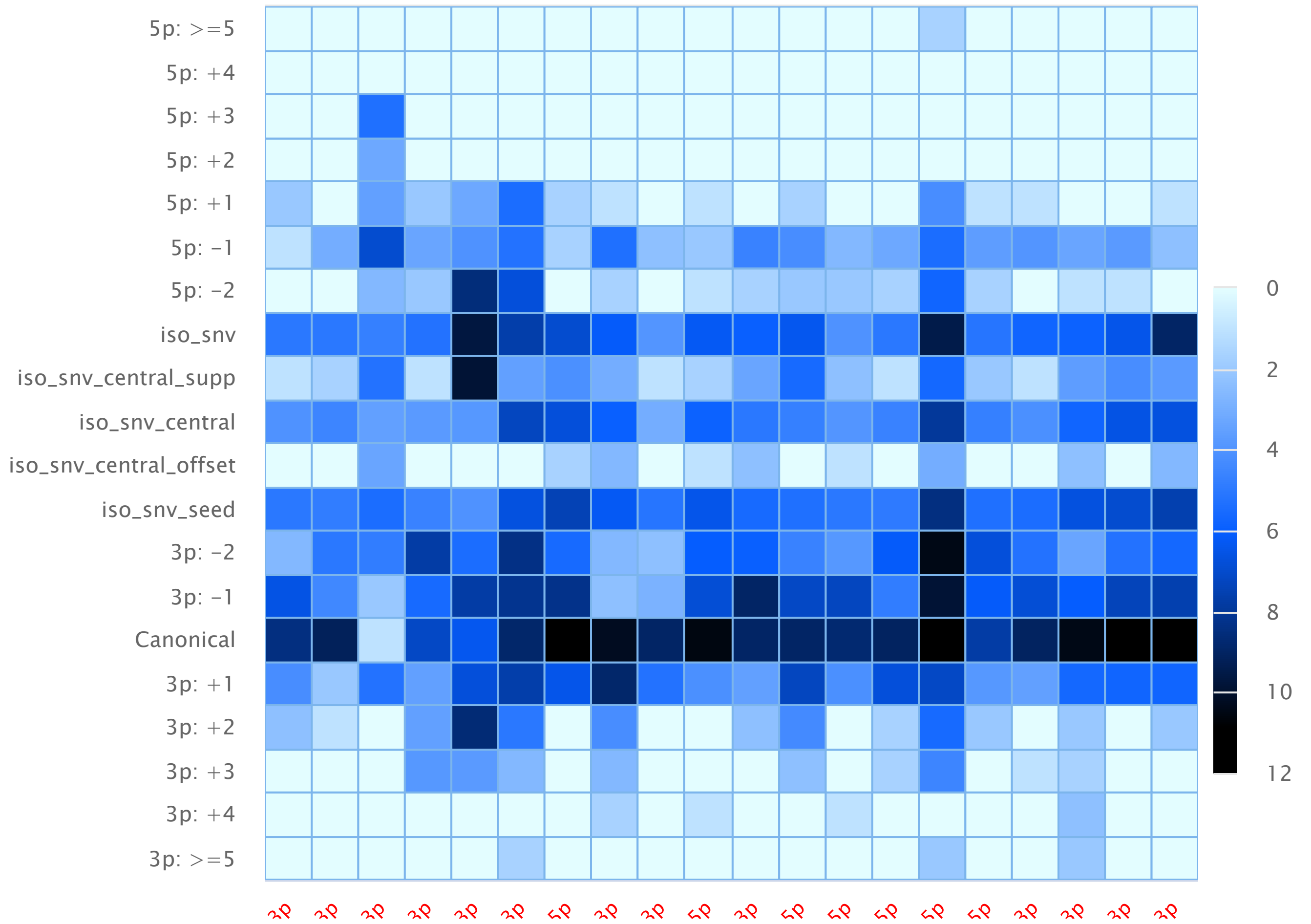
