## Supplementary file 3 for "Integrated miRNA-protein profiling of small extracellular vesicle cargo reveals dynamic cargo remodelling during tissue repair in UV-C-damaged *C. elegans*"

### miRge3.0: Comprehensive analysis of small RNA sequencing Data.

SmallRNA distribution | Read Length | isomiR results | Abundant miRNAs | UMI distribution | Novel miRNAs

#### miR_A1_DU_R1

Read Per Million (RPM) values of 40 most abundant miRNAs

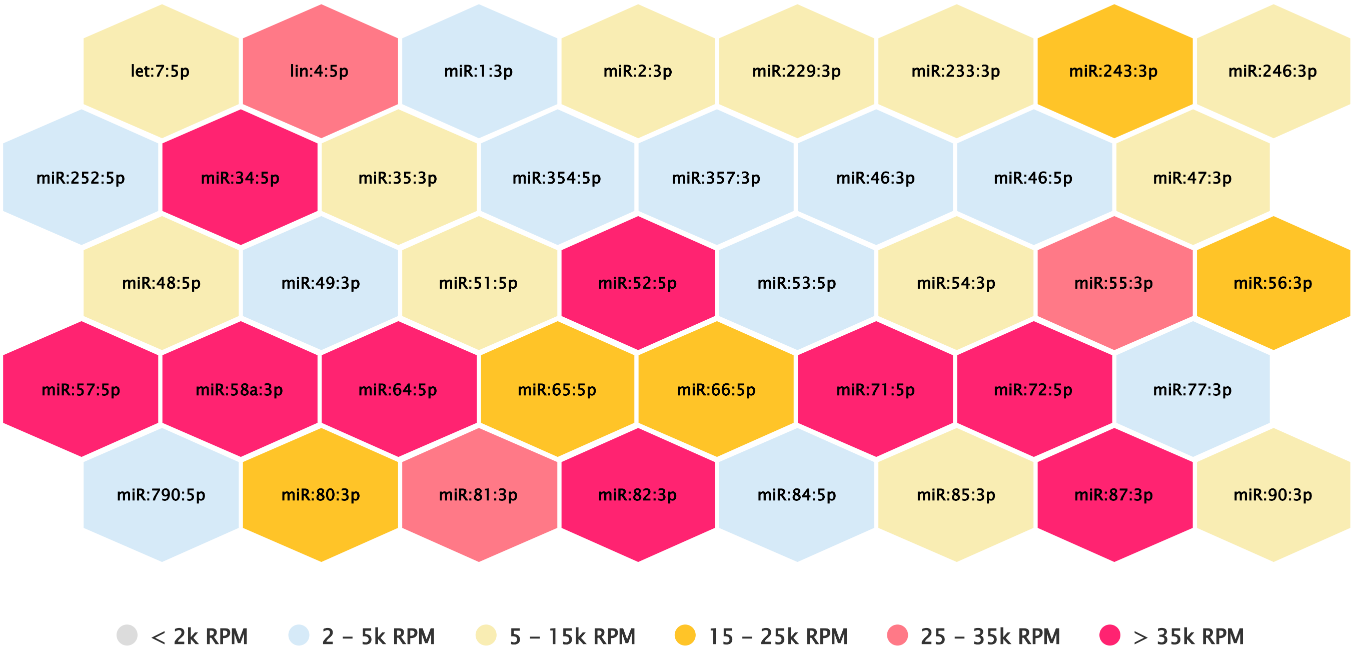


#### miR_A2_DU_R1

Read Per Million (RPM) values of 40 most abundant miRNAs

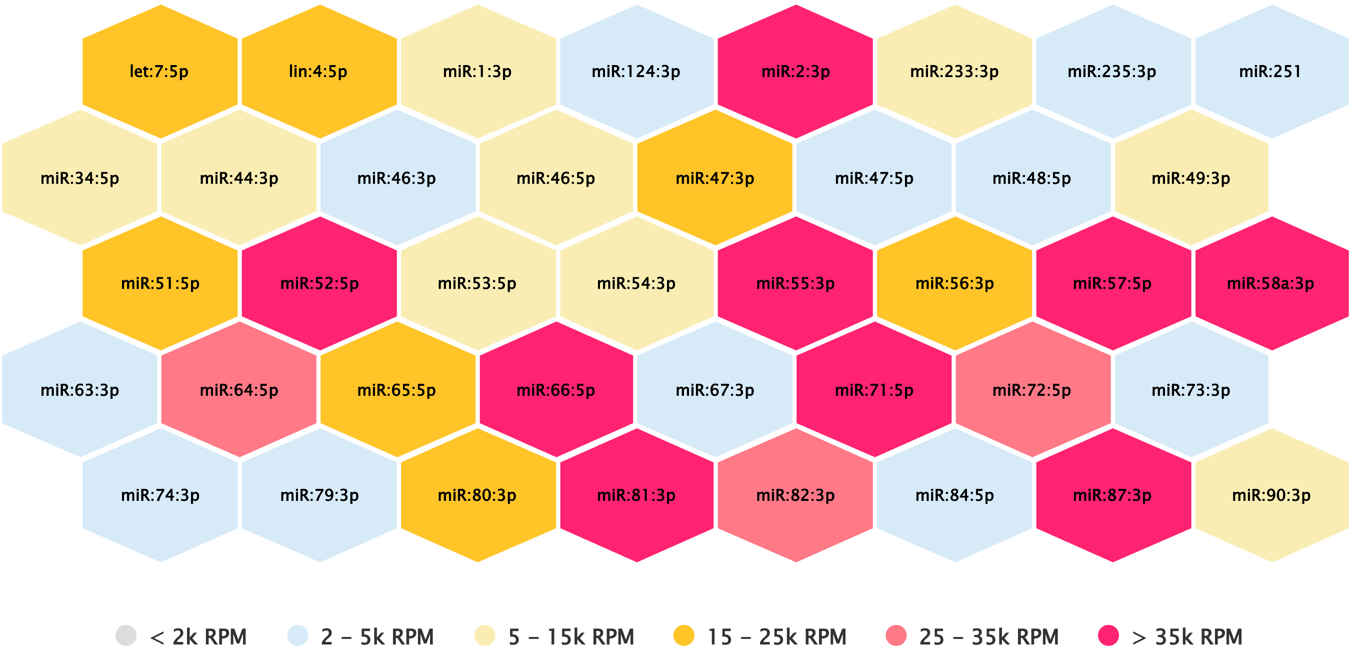

#### miR_A3_DU_R1

Read Per Million (RPM) values of 40 most abundant miRNAs

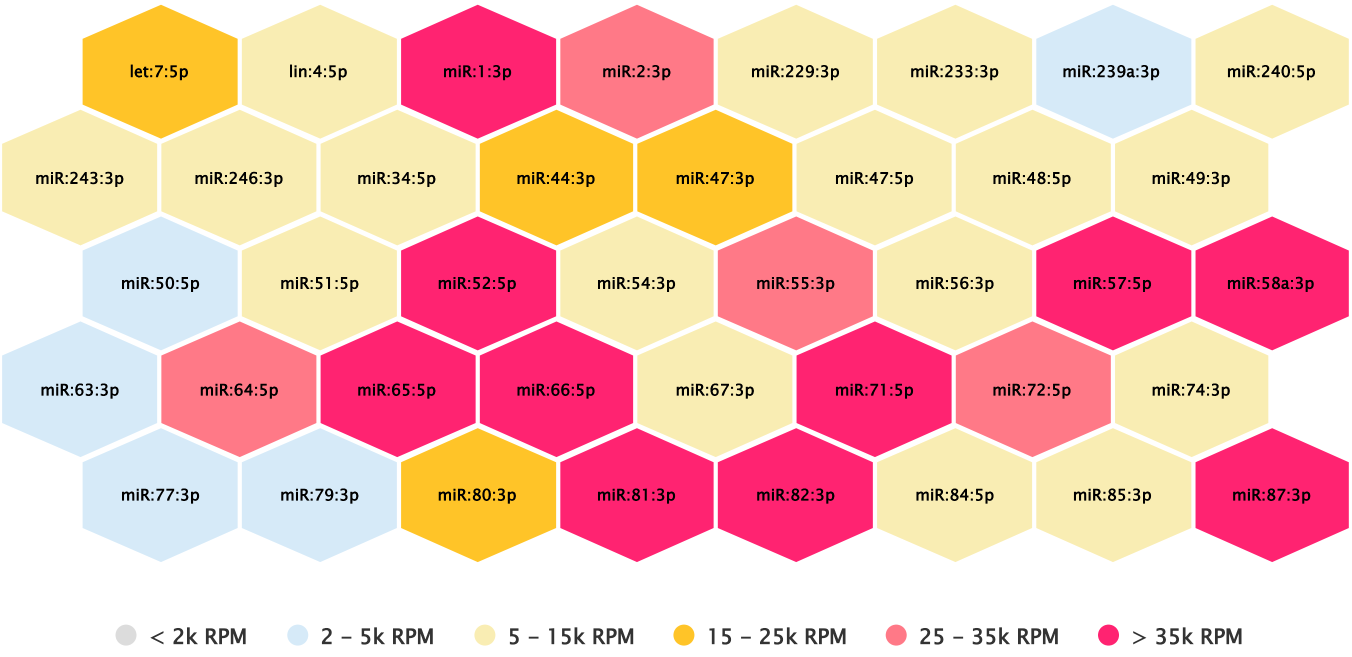


#### miR_B1_DU_R1

Read Per Million (RPM) values of 40 most abundant miRNAs

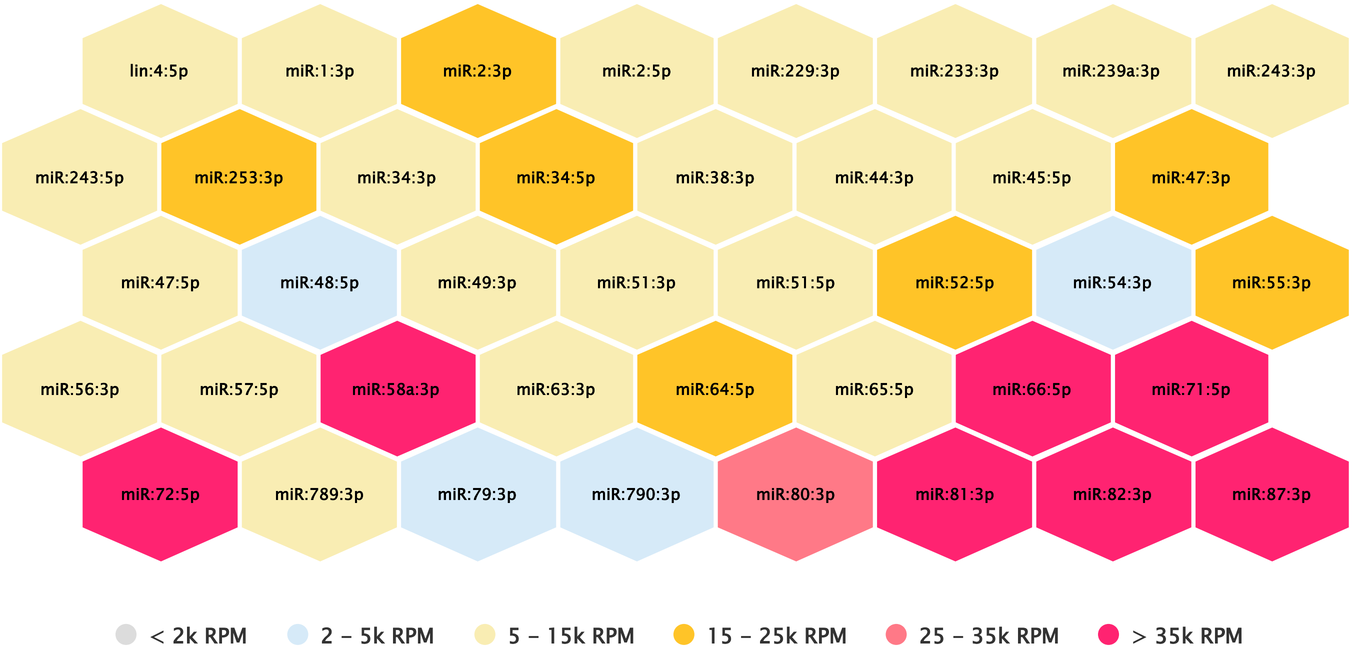

#### miR_B2_DU_R1

Read Per Million (RPM) values of 40 most abundant miRNAs

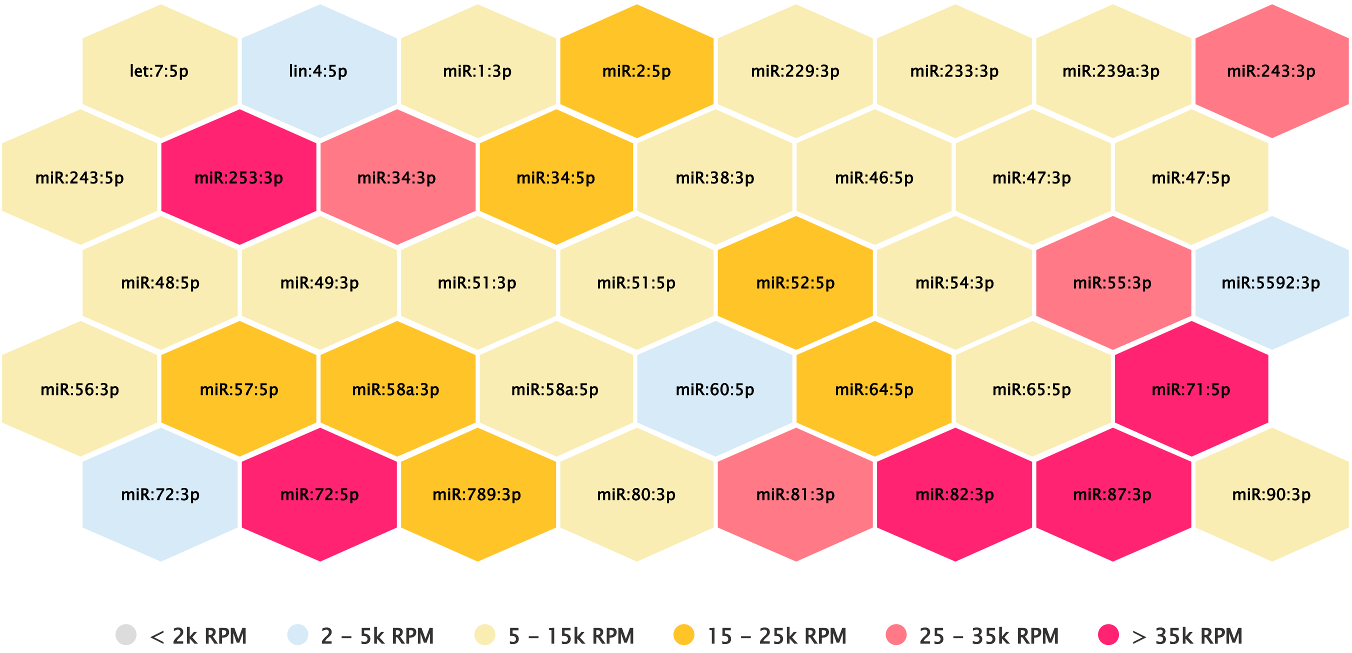


#### miR_B3_DU_R1

Read Per Million (RPM) values of 40 most abundant miRNAs

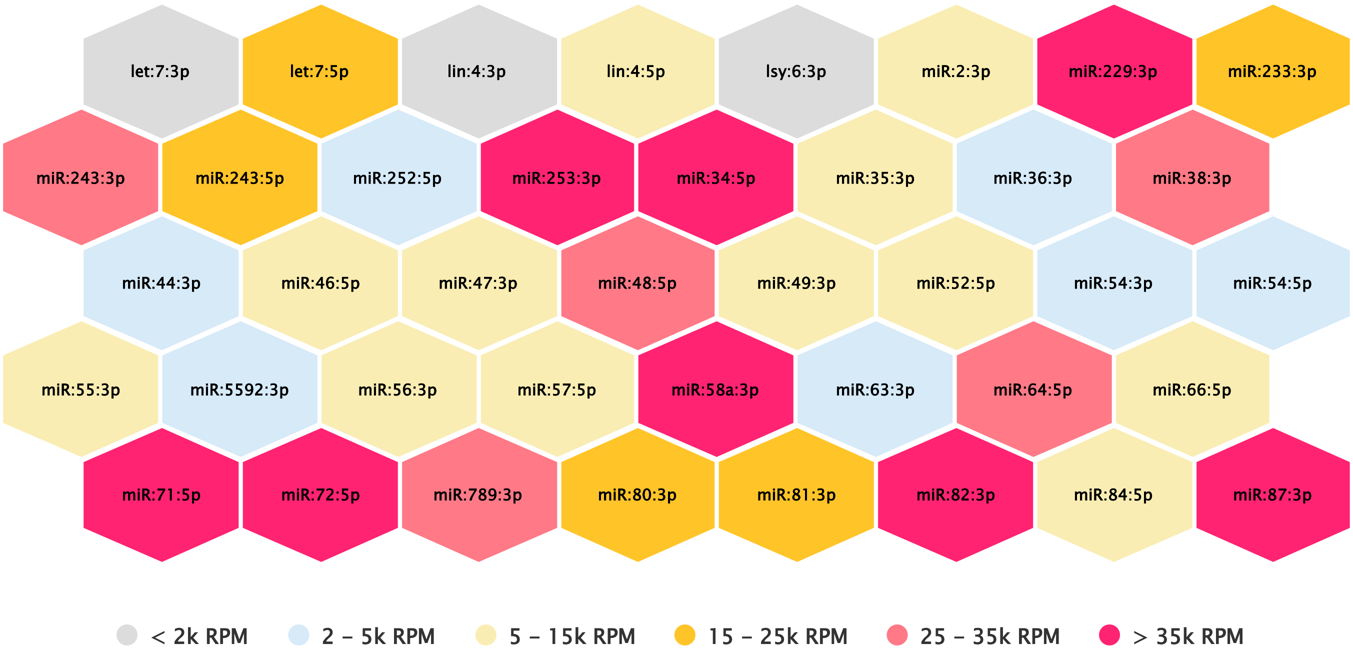

#### miR_C1_DU_R1

Read Per Million (RPM) values of 40 most abundant miRNAs

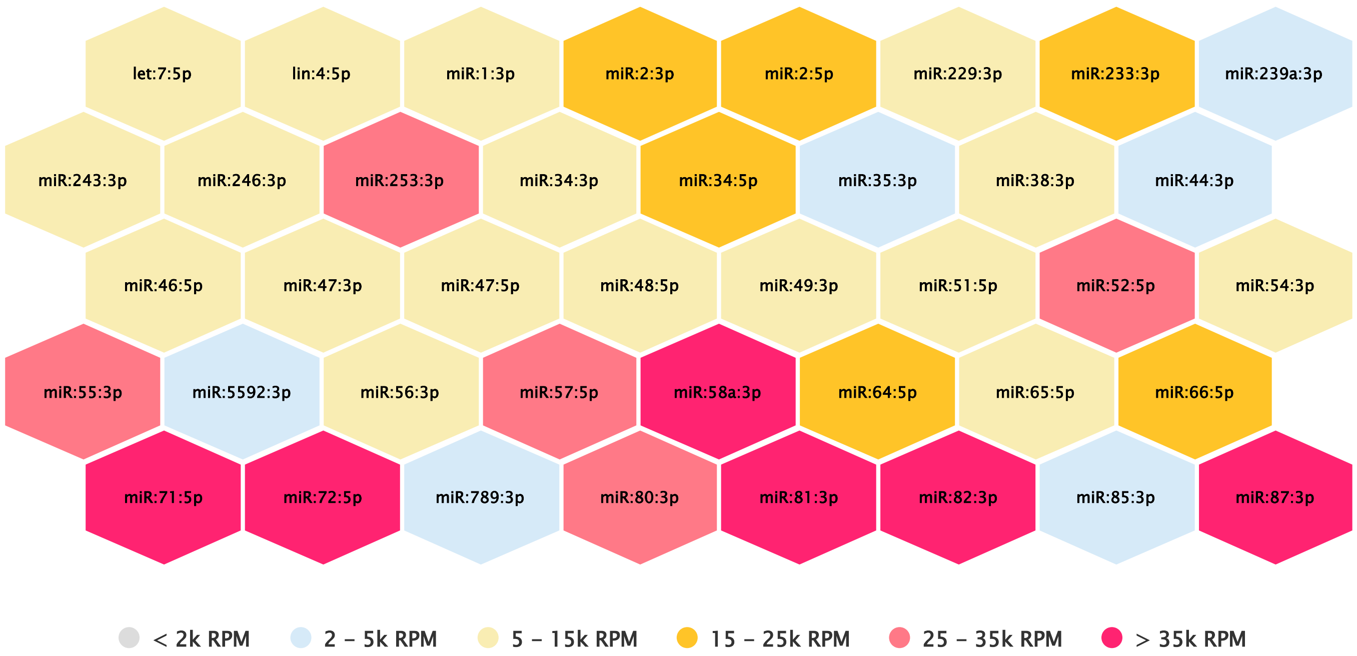


#### miR_C2_DU_R1

Read Per Million (RPM) values of 40 most abundant miRNAs

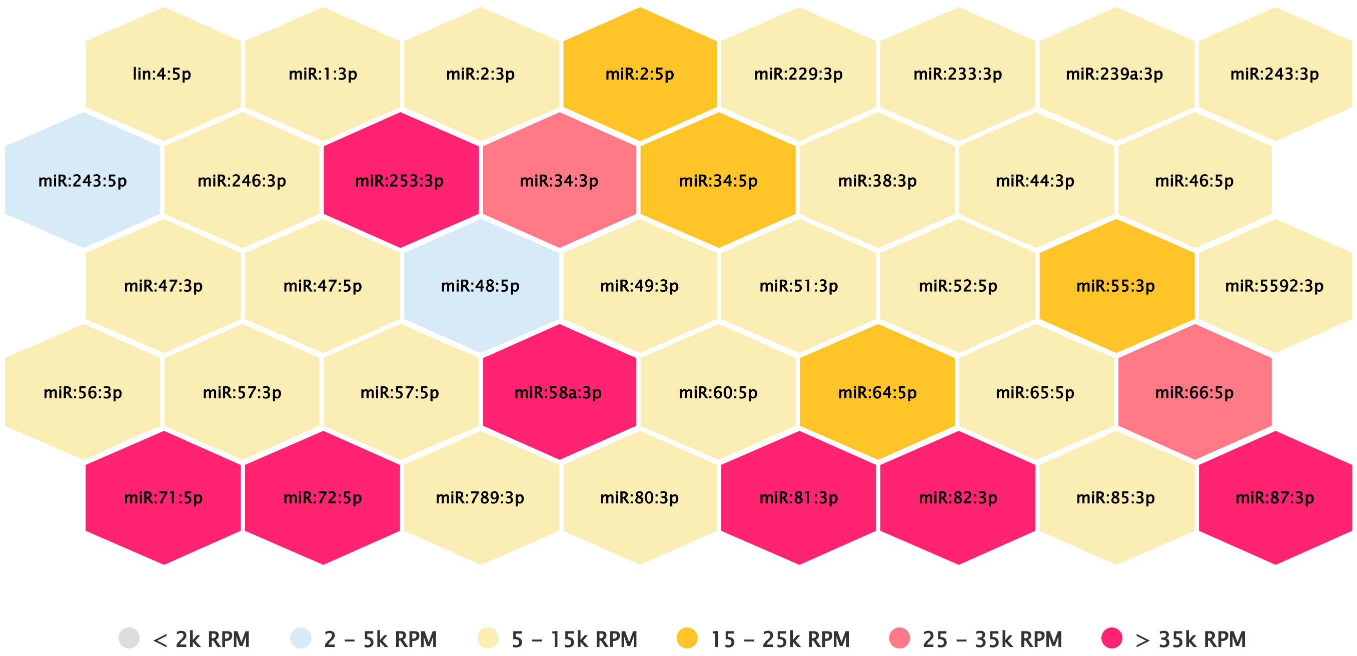

#### miR_C3_DU_R1

Read Per Million (RPM) values of 40 most abundant miRNAs

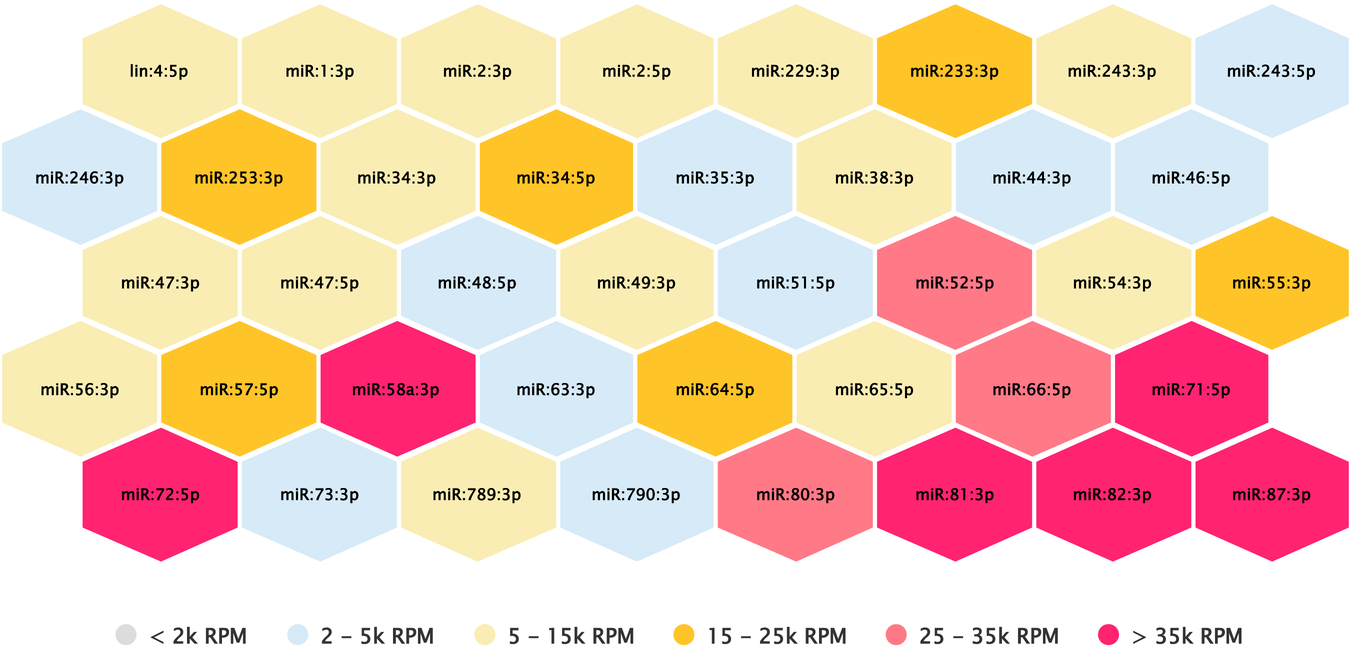


---

#### miR_D1_DU_R1

Read Per Million (RPM) values of 40 most abundant miRNAs

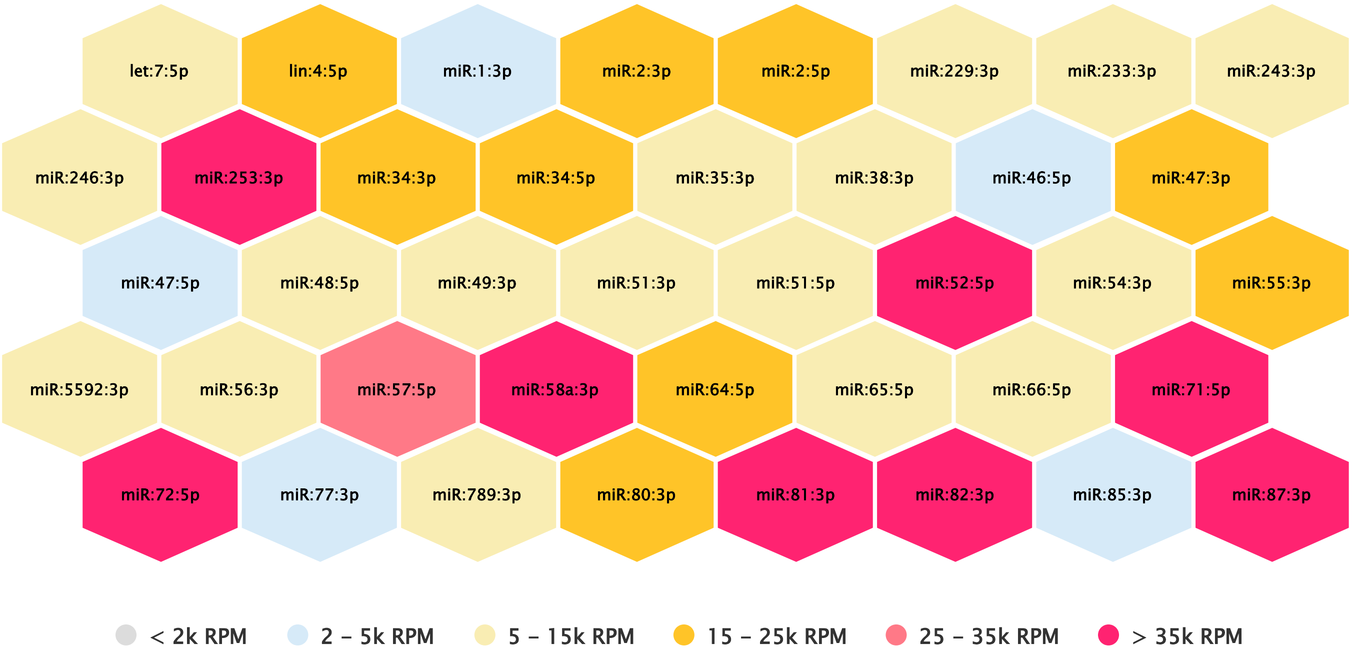


---

#### miR_D2_DU_R1

Read Per Million (RPM) values of 40 most abundant miRNAs

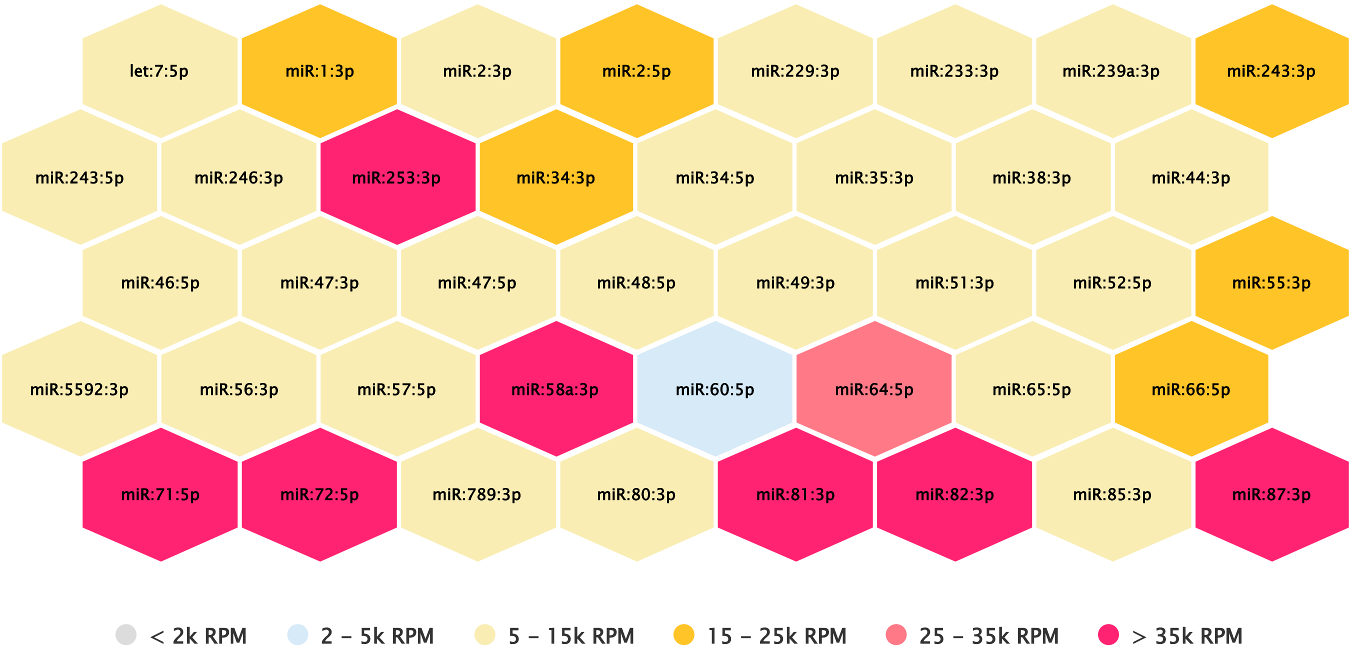


#### miR_D3_DU_R1

Read Per Million (RPM) values of 40 most abundant miRNAs

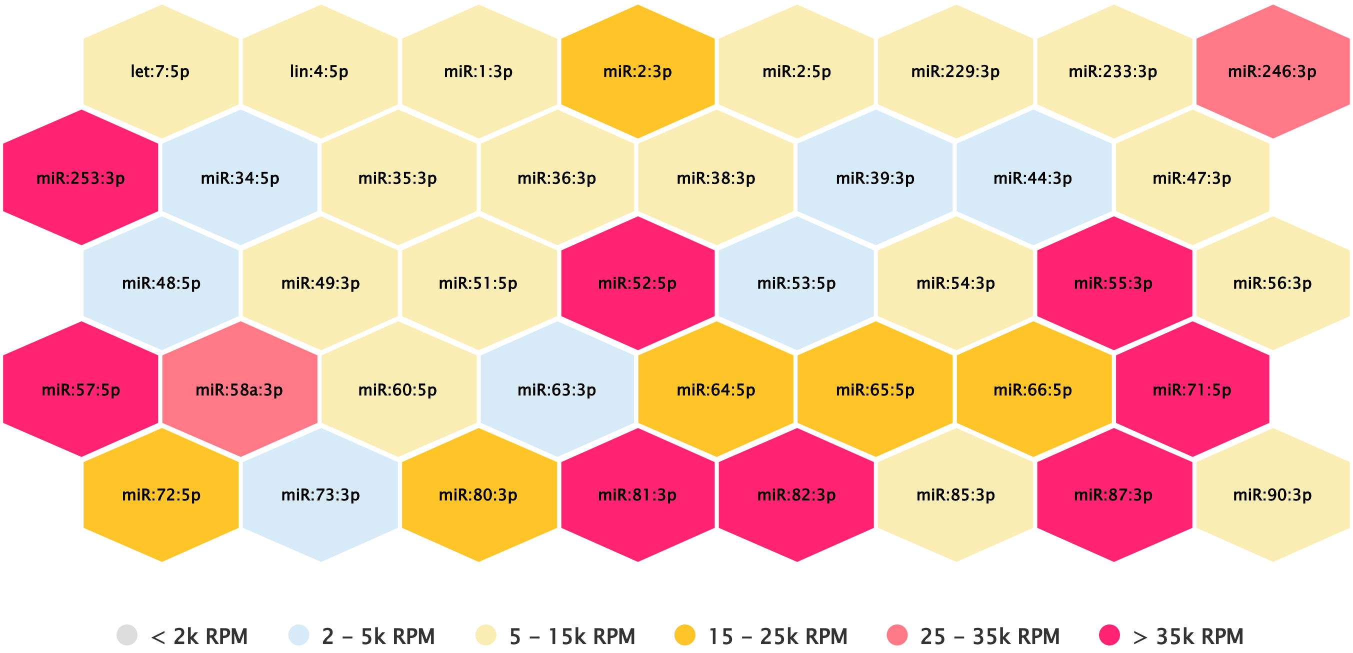
